# Promoter architecture decodes response regulator phosphorylation into distinct transcriptional logics in *Pseudomonas aeruginosa*

**DOI:** 10.64898/2026.07.22.740031

**Authors:** Ziqi Zhu, Ninglin Zhao, Yingjie Song, Dan Fan, Jianping Ying, Cheng Nong, Tonggen Liu, Chenxi Duan, Yaping Zheng, Shuang Zou, Xingyu Mou, Haichuan Ma, Huanxiang Liu, Rui Bao

**Affiliations:** Center of Infectious Diseases, Division of Infectious Diseases in State Key Laboratory of Biotherapy, West China Hospital, Sichuan University, Chengdu 610041, China; College of Life Science, Sichuan Normal University, Chengdu 610101, China; Centre for Artificial Intelligence Driven Drug Discovery, Faculty of Applied Sciences, Macao Polytechnic University, Macao 999078, China

**Keywords:** *Pseudomonas aeruginosa;* two-component system, ParR, response regulator phosphorylation, promoter architecture

## Abstract

Two-component systems convert environmental signals into transcriptional responses, but how one response-regulator phosphorylation state produces different outputs at different promoters remains unclear. Here we show that promoter architecture determines how the ParR response regulator in Pseudomonas aeruginosa interprets phosphorylation. Phenotypic, transcriptomic, biochemical and promoter-engineering analyses showed that phosphorylation of the conserved ParR receiver residue Asp57 lowered DNA-occupancy thresholds across target promoters. Individual promoters, however, converted this shared increase in binding into activation, repression or phosphorylation-dependent sign switching. Chromosomally D57 ParR variants reproduced these behaviours at endogenous loci and altered antibiotic susceptibility, biofilm formation and virulence-associated phenotypes. Mapping and engineering of representative promoters identified a two-tier cis-regulatory code: half-site sequence compatibility determines productive ParR engagement, whereas spacer length determines regulatory sign and magnitude. Spacer swaps were sufficient to reprogram promoter logic between regulatory modes. These findings separate regulator state from promoter decoding and show how bacterial promoter architecture can diversify the outputs of a shared phosphorylation signal.

**Importance:** Pseudomonas aeruginosa is an opportunistic pathogen that survives antimicrobial treatment and adapts to diverse host environments by rapidly changing gene expression. How a single sensory pathway generates different, and sometimes opposing, responses across many genes has remained unclear. We show that the regulatory protein ParR does not impose one fixed transcriptional program. Instead, the DNA architecture of each target promoter determines whether the same phosphorylation signal activates, represses, or reverses gene expression. The sequence and spacing of ParR recognition sites form a compact regulatory code, and changing spacer length alone can reprogram promoter behavior. Different ParR states were also associated with changes in antibiotic susceptibility, biofilm formation, and virulence-associated phenotypes. Our findings explain how one bacterial signaling pathway can generate diverse physiological outcomes without additional regulators and provide a framework for understanding regulatory flexibility in Pseudomonas aeruginosa and for engineering genetic circuits with multiple outputs.

**Graphical abstract:** 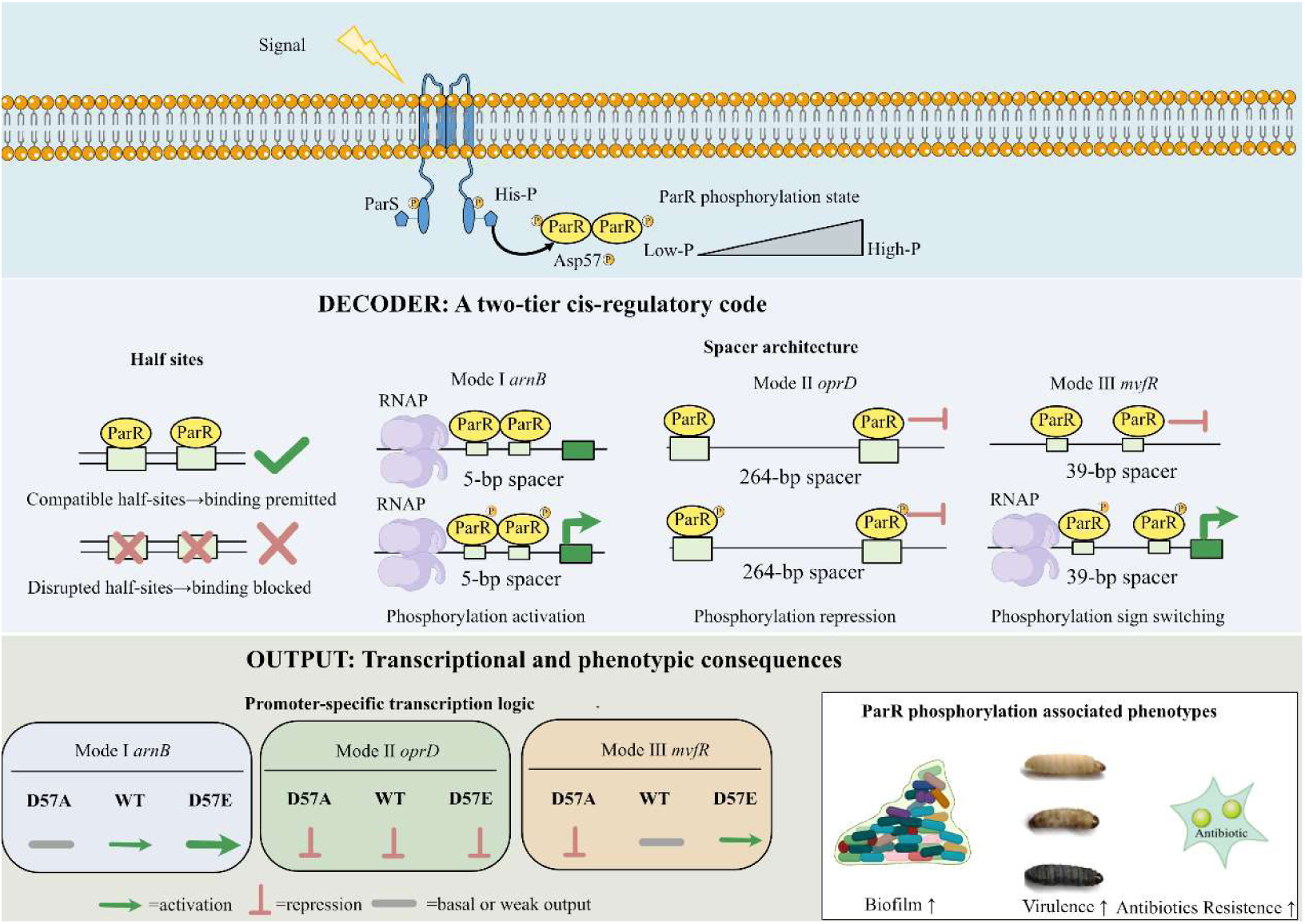

## 1. Introduction

*Pseudomonas aeruginosa* is an opportunistic pathogen that persists in diverse host niches and often withstands antimicrobial therapy. Its adaptability depends not only on intrinsic resistance determinants, such as low outer-membrane permeability and broad efflux capacity, but also on reversible tolerance states that arise in biofilms and chronic-infection microenvironments(1, 2). A recurring constraint in this lifestyle is resource allocation. The bacterium must balance resistance programmes with metabolically costly virulence and biofilm modules, priorities that can impose context-dependent fitness trade-offs under host pressure (3). Two-component systems (TCSs) are central to this coordination, translating environmental cues into cellular responses and contributing repeatedly to *P. aeruginosa* virulence and stress adaptation (4, 5). Although the core TCS pathway is well established, with histidine kinase activation followed by response regulator (RR) phosphorylation, it remains unclear how a common phosphorylation input is decoded by different promoters to generate distinct, and sometimes opposing, transcriptional outputs.

Among envelope stress-responsive TCSs in *P. aeruginosa*, ParRS provides a useful model for this decoding problem. In addition to its established role in lipopolysaccharide remodelling and polymyxin adaptation, ParRS has been linked to efflux induction, porin repression and other envelope-associated defence programmes(6–8). Our previous work further showed that host-derived peptide environments intersect with ParRS-centred envelope stress signalling to reshape pathogenic outputs, supporting ParRS as an infection-relevant decision node rather than a narrow polymyxin module(9). The biophysical rules that permit this regulatory plasticity remain unresolved. Phosphorylation is classically viewed as a binary switch that stabilizes active response-regulator conformations and promotes dimerization for DNA engagement. However, recent biochemical and structural studies of OmpR/PhoB-family transcription complexes indicate that promoter DNA arrangement and RNA polymerase (RNAP) contact geometry can reshape the output of a shared RR state (10, 11). The key question is how RR target promoters encode cis-regulatory rules that convert one signalling state into activation, repression or switching across a heterogeneous regulon.

We hypothesized that the specificity of ParR output is not intrinsic to the RR’s phosphorylation state alone but is encrypted within the *cis*-regulatory architecture of its target promoters. In this model, phosphorylation of the conserved receiver aspartate (D57) primarily controls signal strength and serves as a state variable, whereas the architecture of the binding motifs, including their sequence compatibility and spatial arrangement relative to the core promoter, determines the direction and magnitude of the transcriptional response. This two-tier decoding mechanism would allow a common activation signal to be differentially interpreted across the genome, enabling the precise coordination of defense and offense strategies. This hypothesis is motivated by a broader emerging principle in bacterial transcription: the same transcription factor can exhibit markedly different apparent functions depending on promoter context, with quantitative rules linking basal promoter properties to regulatory fold change(12). Synthetic biology studies likewise show that local changes in promoter or operator architecture, including strength, position, spacing and insulation, can generate distinct input-output behaviours without changing upstream protein components(13).

Here, we tested this decoding model by combining physiological phenotyping, transcriptomics, biochemical binding assays and cis-element engineering. We used chromosomal editing to lock ParR into defined nonphosphorylatable (ParRD57A) and phosphomimetic (ParRD57E) states, and linked these states to antibiotic susceptibility, biofilm formation and host-infection phenotypes. We then integrated global transcriptional profiling with promoter-level validation to define operational rules for ParR-DNA engagement. Finally, synthetic promoter engineering showed that half-site compatibility and spacer architecture can be separated experimentally, and that spacer changes can convert ParR-dependent activation into repression or sign switching. These analyses provide a promoter-architecture framework for ParR signal decoding and suggest a route by which TCS response regulators diversify outputs across a regulon.

## 2. Materials and methods

### 2.1. Bacterial strains, plasmids and growth conditions

All *P. aeruginosa* strains used in this study were derived from PA14 wild-type (WT). Strains, plasmids and primers are listed in Tables S1 and S2. Unless stated otherwise, bacteria were cultured aerobically in lysogeny broth (LB) at 37°C with shaking at 220 rpm. For MIC assays, strains were cultured in Mueller-Hinton broth (MHB). For RNA sequencing (RNA-seq), cultures were grown under identical conditions to mid-log phase (OD_600_ of approximately 0.4–0.6). Cells were harvested rapidly by centrifugation, flash-frozen in liquid nitrogen and stored at −80°C until RNA extraction.

### 2.2. Construction of PA14 mutants and complemented strain

The in-frame deletion mutant Δ*parR*, the chromosomally complemented strain C-Δ*parR*, and the chromosomal phosphorylation site mutants *parR*D57A and *parR*D57E were generated in the *P. aeruginosa* PA14 background by homologous recombination using the suicide vector pEX18Gm. To construct Δ*parR*, approximately 500-bp upstream and downstream homologous arms flanking *parR* were amplified from PA14 genomic DNA and cloned into pEX18Gm. To construct of C-Δ*parR*, the wild-type *parR* allele with its upstream and downstream homologous regions was cloned into pEX18Gm. To construct *parR*D57A and *parR*D57E, the corresponding point mutations were introduced into *parR* by overlap-extension PCR using mutagenic primers (Table S2), and the mutant alleles with approximately 500-bp flanking homologous arms were cloned into pEX18Gm. Recombinant plasmids were verified by Sanger sequencing, introduced into *Escherichia coli* S17-1, and transferred into PA14 by biparental mating on LB agar at 30°C for 12-16 h. Single-crossover integrants were selected on LB agar containing gentamicin (50 μg/mL) and triclosan (26 μg/mL).

Double-crossover recombinants were obtained by counterselection on NaCl-free LB agar supplemented with 15% (w/v) sucrose. Candidate colonies were screened by colony PCR using flanking primers (Table S2). Edited loci were confirmed by Sanger sequencing to verify the intended deletion, complementation or point mutation and to exclude unintended sequence changes within the recombined region.

### 2.3. Crystal violet biofilm assay

Biofilm formation was quantified by crystal violet (CV) staining in standard 96-well microtiter plates (Biosharp). Overnight cultures were diluted 1:100 into fresh LB medium, and 100 μL aliquots were added to each well. Plates were incubated statically at 37°C for 24 h. Planktonic cells were removed, and wells were rinsed gently three times with distilled water. Attached biofilms were fixed with methanol for 10 min, stained with 0.1% (w/v) CV for 10 min, and washed thoroughly to remove excess dye. Bound CV was solubilized with 30% (v/v) glacial acetic acid, and the absorbance was measured at 570 nm using a SuPerMax 3500 microplate reader (Shanghai Flash Bio-Tech Co., Ltd.). Each experiment included sterile medium controls and was performed with at least three independent biological replicates, each containing three technical replicates. Data are presented as mean ± SD. Statistical significance was evaluated by one-way ANOVA followed by Tukey’s post hoc test.

2.4. *Galleria mellonella* infection assay

Virulence was evaluated using a *Galleria mellonella* wax moth larval model. Bacterial cultures were grown in LB at 37°C with shaking to mid-log phase (OD_600_ of approximately 0.4–0.6), harvested by centrifugation and washed three times with sterile phosphate buffered saline (PBS). Washed cells were resuspended in PBS and adjusted to 4 × 10^3^ CFU/mL. For each strain, larvae (n = 16) were injected with 10 μL bacterial suspension (approximately 40 CFU per larva) into the hemocoel via the last left proleg using a Hamilton microsyringe fitted with a 30-gauge needle. Control larvae were injected with 10 μL sterile PBS. Larvae were maintained at 37°C and monitored for survival; death was scored as lack of movement in response to tactile stimulation accompanied by complete melanization. Survival differences were analyzed using the log-rank (Mantel-Cox) test.

### 2.5. RNA sequencing and transcriptomic analysis

Strand-specific RNA-seq libraries were prepared from high-quality total RNA after removal of genomic DNA and depletion of ribosomal RNA using the RiboCop rRNA Depletion Kit for Mixed Bacterial Samples (Lexogen, USA). The rRNA-depleted RNA was fragmented into approximately 200-nt fragments and reverse-transcribed with random hexamer primers, and strand specificity was preserved by incorporation of dUTP during second-strand cDNA synthesis. After end repair, phosphorylation, and A-tailing, libraries were constructed using the Illumina Stranded mRNA Prep, Ligation kit and sequenced on the Illumina NovaSeq 6000 platform to generate paired-end reads. Raw reads were filtered to remove adaptor-containing reads, low-quality reads, and reads containing more than 10% ambiguous bases. Clean reads were mapped to the *P. aeruginosa* UCBPP-PA14 reference genome (ASM1462v1; RefSeq assembly GCF_000014625.1; GenBank assembly GCA_000014625.1) using Bowtie2, and rRNA contamination was assessed by aligning 10,000 randomly selected raw reads from each sample to the Rfam database using BLAST. Transcript abundance, including reference genes, novel transcripts, and sRNAs, was quantified using RSEM and normalized as transcripts per million (TPM). No batch correction was applied. Differential expression analysis was performed using DESeq2 with pairwise comparisons among the WT, Δ*parR*, *parR*D57A, and *parR*D57E groups. Genes with a Benjamini–Hochberg-adjusted *P* value < 0.05 and a fold change ≥ 1.5 were considered differentially expressed. Functional enrichment analyses were performed using GOATOOLS for Gene Ontology analysis and KOBAS 2.0 for KEGG pathway analysis.

### 2.6. Minimum inhibitory concentration (MIC) determination

MICs were determined by broth microdilution in sterile 96-well plates. Overnight cultures grown in MHB were diluted into fresh MHB to yield a final inoculum of approximately 5 × 10⁵ CFU/mL per well (final volume 200 μL). Antibiotics were prepared as two-fold serial dilutions in MHB. Wells containing inoculated MHB without antibiotic served as growth controls, and wells containing MHB only served as sterility controls. Plates were incubated statically at 37°C for 16-20 h. The MIC was recorded as the lowest antibiotic concentration with no visible growth. Unless otherwise stated, MIC assays were performed with at least three independent biological replicates.

### 2.7. RNA isolation and quantitative RT-PCR

Total RNA was extracted from bacterial cultures using TRIzol reagent (Vazyme) according to the manufacturer’s instructions. Residual genomic DNA was removed using the gDNA Eraser component of the PrimeScript RT reagent kit (Vazyme). cDNA was synthesized from 1 μg total RNA using the same kit. qRT-PCR was performed in 20 μL reactions containing 10 ng cDNA, 200 nM gene-specific primers (Table S2), and 2× ChamQ SYBR qPCR Master Mix (Vazyme). The *oprL* gene was used as an internal reference gene for normalization of RT-qPCR data. Reactions were run on a QuantStudio 6 Pro system with the following cycling conditions: 95°C for 30 s; 40 cycles of 95°C for 10 s and 60°C for 30 s; followed by melt-curve analysis. Each sample was analyzed with three technical replicates, and at least three independent biological replicates were performed. Relative expression was calculated using the 2^−ΔΔCt^ method. Statistical comparisons were performed using one-way ANOVA followed by the appropriate post hoc test for experiments involving three or more groups, or by two-tailed unpaired Student’s *t*-test for predefined two-group comparisons.

### 2.8. Heterologous expression and purification of ParR and variants

The *parR* coding sequence was amplified from PA14 genomic DNA and cloned into pET-22b (Novagen) using the ClonExpress II One Step Cloning Kit (Vazyme) to generate a construct expressing ParR fused to a C-terminal 6×His tag (pET 22b-ParR). ParR phosphorylation-site variants, ParRD57A and ParRD57E, were generated by PCR-based site-directed mutagenesis and confirmed by Sanger sequencing. The resulting plasmids were transformed into *E. coli* BL21(DE3). Cells were grown in LB supplemented with ampicillin (50 μg/mL) at 37°C to OD_600_ of approximately 0.6-0.8. Protein expression was induced with 0.4 mM IPTG for 14 h at 16°C. Cells were harvested by centrifugation (3,500 rpm, 10 min, 4°C), resuspended in lysis buffer (25 mM Tris-HCl, pH 7.5, 150 mM NaCl) supplemented with 1 mM PMSF, and lysed by high-pressure homogenization. The soluble fraction was obtained by centrifugation (15,000 rpm, 30 min, 4°C) and purified by native Ni^2+^ affinity chromatography using a nickel affinity column (Qiagen), with elution in 300 mM imidazole. Proteins were buffer-exchanged and concentrated using Amicon Ultra centrifugal filters (10-kDa cutoff), followed by size-exclusion chromatography on an ÄKTA system (Cytiva) equipped with a Bio 70 column equilibrated in storage buffer (25 mM Tris-HCl, pH 7.5, 150 mM NaCl, 10% glycerol). Protein purity was assessed by SDS-PAGE. Purified proteins were concentrated, flash-frozen in liquid nitrogen, and stored at −80°C.

### 2.9. Electrophoretic mobility shift assays (EMSA)

Protein-DNA interactions were analyzed by EMSA. Promoter DNA fragments were amplified from PA14 genomic DNA and purified. Binding reactions (10 μL) were assembled in EMSA buffer (25 mM Tris-HCl, pH 7.5, 150 mM NaCl, 10% glycerol) containing 0.1 μM DNA probe and the indicated ParR concentrations. For promoter-wide occupancy mapping and phosphorylation-state comparisons, ParR was titrated from 0 to 0.8 μM. For in vitro phosphorylation, ParR was pre-incubated with 20 mM acetyl phosphate (AcP) for 30 min before binding. After incubation at 4°C for 30 min, samples were mixed with native loading dye and resolved on 12% native polyacrylamide gels in pre-chilled 0.5× TBE at 130 V for 90 min at 4°C (with buffer circulation). Gels were stained with ethidium bromide (0.5 μg/mL), destained in water and imaged using a ChemiDoc MP system. Each EMSA was repeated at least three times. Promoter probes were generated either by PCR amplification from PA14 genomic DNA or by annealing commercially synthesized oligonucleotides for short engineered ParR-binding constructs, as indicated in Table S2.

### 2.10. â-Galactosidase reporter assays

Promoter activity was measured using a transcriptional *lacZ* reporter. Promoter regions were cloned upstream of promoterless *lacZ* in pRG970Km. Effector plasmids encoding ParR or variants were constructed in pET 22b. Reporter and effector plasmids were co-transformed into *E. coli* DH5α and selected on LB agar containing kanamycin (50 μg/mL) and ampicillin (100 μg/mL). Single colonies were inoculated into LB with antibiotics and grown at 37°C to OD_600_ of approximately 0.6-0.7. Cells were pelleted (3,500 rpm, 15 min, 4°C) and resuspended in 1 mL Z-buffer (60 mM Na₂HPO₄, 40 mM NaH₂PO₄, 10 mM KCl, 1 mM MgSO₄, pH 7.0). Cells were permeabilized with 50 μL 1% (w/v) SDS and 100 μL chloroform, vortexed for 15 s, and reactions were initiated by adding 100 μL ONPG (4 mg/mL) at 28°C. After 10 min, reactions were stopped with 150 μL 1 M Na₂CO₃. After clarification (12,000 rpm, 10 min), A_420_ and A_550_ were measured. β-galactosidase activity (Miller units) was calculated as 1000 × [A_420_-(1.7 × A_550_)] / (T × V × OD_600_), where T is reaction time (min) and V is culture volume (mL). Each condition was assayed with three technical replicates and at least three independent biological replicates. Statistical comparisons were performed by one-way ANOVA followed by Dunnett’s post hoc test against the specified control.

### 2.11. Motif discovery and phylogenetic analyses

For motif discovery, upstream promoter regions of ParR-regulated genes were extracted from the PA14 genome and analyzed using MEME to identify enriched conserved motifs (motif width set to 8 bp; both strands allowed). The resulting consensus motif was used for downstream ParR binding site engineering and probe design. ParR homologs were identified using DIAMOND BLAST against the Pseudomonas Genome Database and BLASTP against UniProt using PA14 ParR as query (query coverage >50% and identity >50%). The top UniProt hits were aligned in BioEdit, phylogenetic trees were reconstructed using the Maximum Likelihood method in MEGA12, and trees were visualized and annotated using ChiPlot/tvBOT(14).

### 2.12. Statistical analysis

Unless stated otherwise, data are presented as mean ± SD from at least three independent biological replicates and were analyzed using GraphPad Prism 9.0. Multiple-group comparisons were performed using one-way ANOVA followed by Tukey’s post hoc test, or Dunnett’s post hoc test when comparisons were made against a single control. Two-group comparisons were performed using two-tailed unpaired Student’s *t*-test. Survival curves were analyzed using the log-rank (Mantel-Cox) test. Exact n values, statistical tests, and significance information for each experiment are provided in the figure legends or main text. A *P* value < 0.05 was considered statistically significant.

## 3. Results

### 3.1 ParR coordinates antibiotic resistance and virulence-associated phenotypes

To assess whether ParR affects baseline antimicrobial susceptibility in *P. aeruginosa* PA14, we determined MICs for the *parR* deletion mutant (Δ*parR*) in Mueller-Hinton (MH) medium. Deletion of *parR* reduced MICs by one two-fold dilution step across the tested polymyxins, aminoglycosides, and β-lactams. Genetic complementation restored these values to WT levels (Table 1). These results indicate that ParR contributes to intrinsic defense profiles, consistent with reports linking ParRS to adaptive polymyxin resistance and cross-class susceptibility phenotypes(6, 8, 15).

**Table 1.**
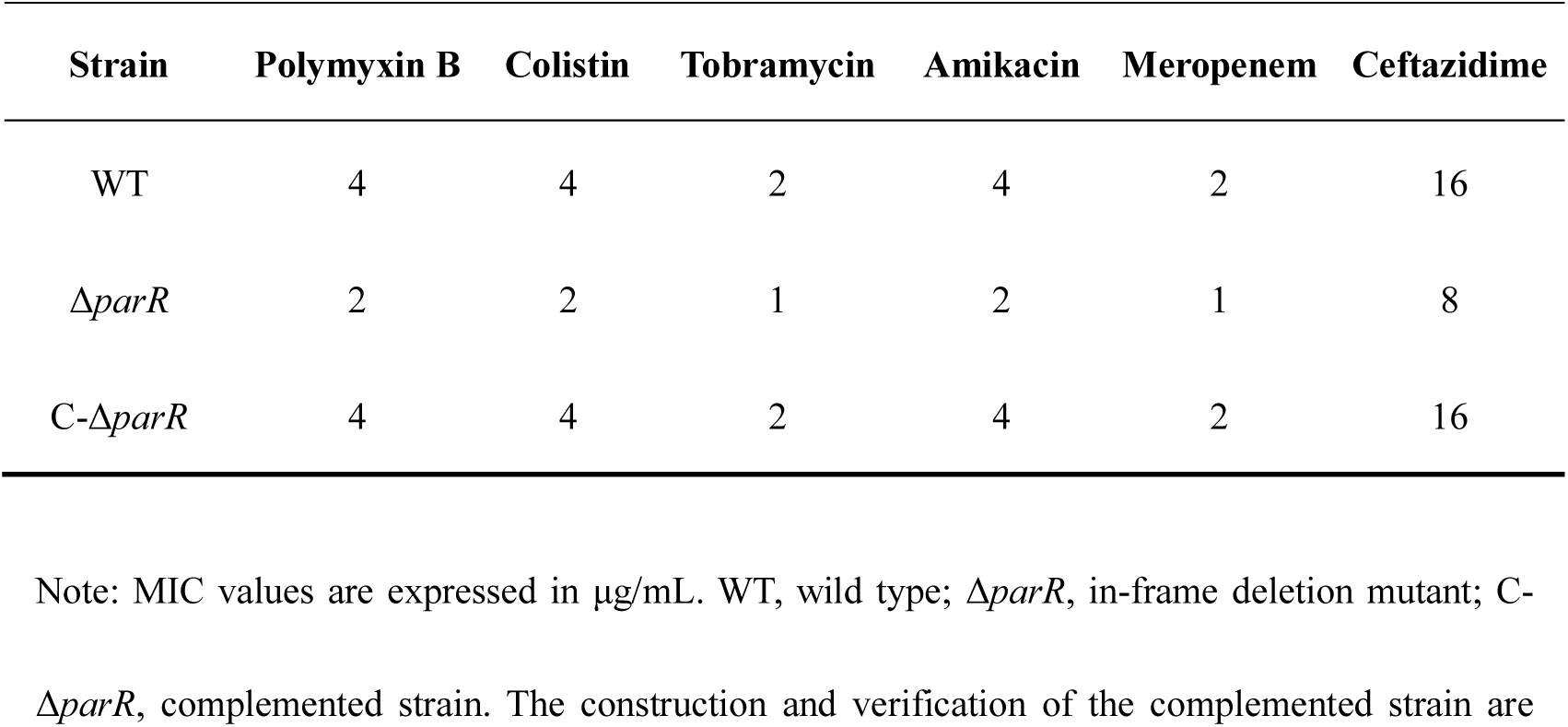
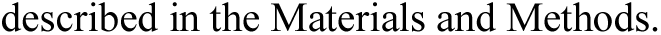
Minimum inhibitory concentrations (MICs) of WT, Δ*parR*, and the complemented strain in Mueller-Hinton broth.

| Strain | Polymyxin B | Colistin | Tobramycin | Amikacin | Meropenem | Ceftazidime |
| --- | --- | --- | --- | --- | --- | --- |
| WT | 4 | 4 | 2 | 4 | 2 | 16 |
| Δ*parR* | 2 | 2 | 1 | 2 | 1 | 8 |
| C-Δ*parR* | 4 | 4 | 2 | 4 | 2 | 16 |

Note: MIC values are expressed in μg/mL. WT, wild type; $\Delta$ *parR*, in-frame deletion mutant; C-Δ *parR*, complemented strain. The construction and verification of the complemented strain are
described in the Materials and Methods.

We next examined the role of ParR in biofilm formation using a static crystal violet microtiter-plate assay. The Δ*parR* mutant exhibited a modest but reproducible increase in surface-associated biomass relative to WT (approximately 25% higher; Fig. 1A). Complementation shifted the biofilm signal back toward WT levels (Fig. 1A), supporting a *parR*-dependent phenotype. This directionality is consistent with previous observations that *parS*/*parR* mutants exhibit elevated quorum-sensing and secondary-metabolite outputs, and that *parS* mutations frequently arise during antibiotic driven evolution in *in vivo* biofilm models(16, 17).

**Fig. 1.**
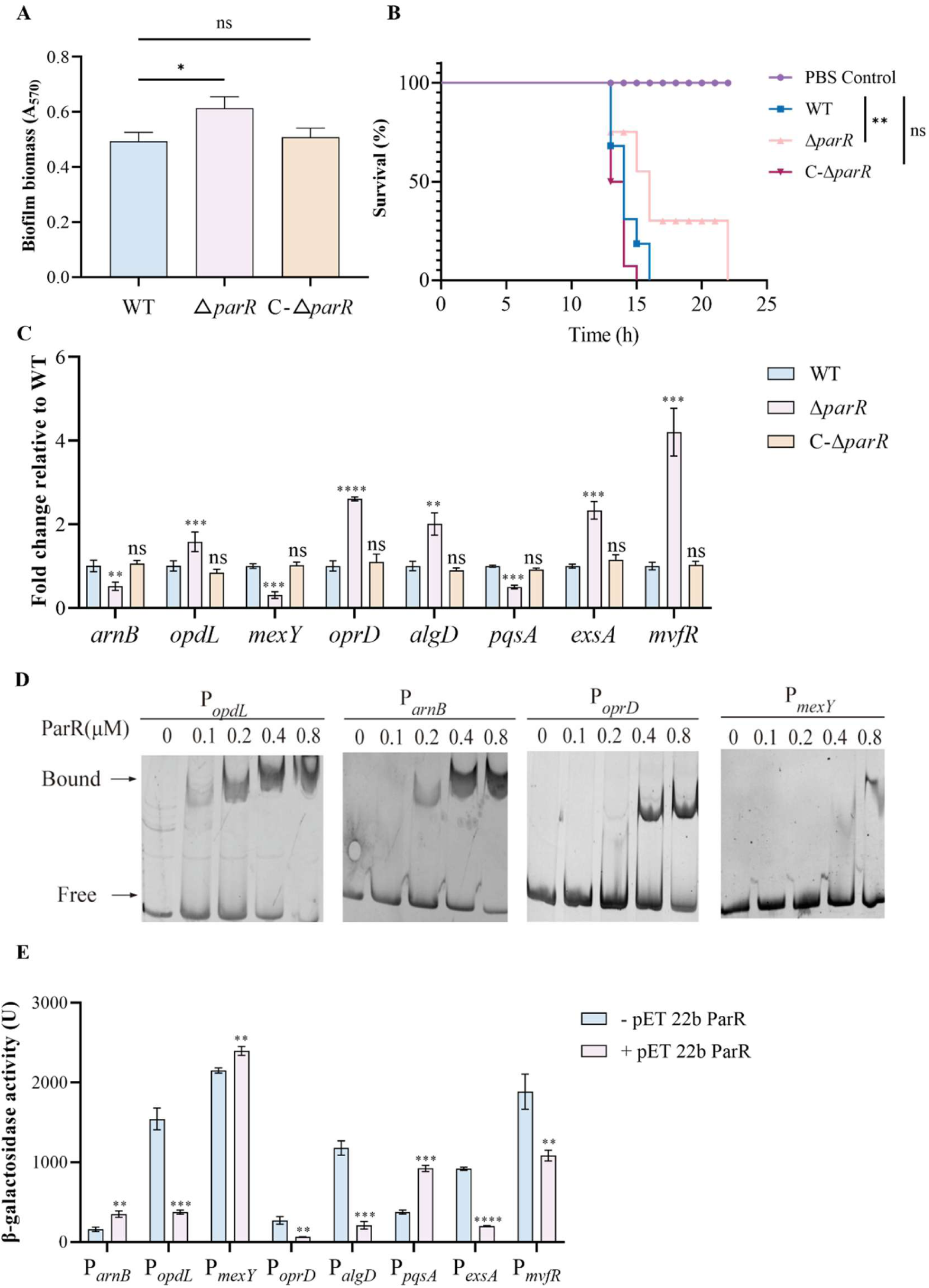
ParR links physiological phenotypes to direct promoter binding and bidirectional transcriptional outputs. (A) Biofilm biomass of WT PA14, Δ*parR*, and C-Δ*parR*, quantified by crystal violet staining and measured as absorbance at 570 nm. (B) Survival of *Galleria mellonella* larvae after infection with WT PA14, Δ*parR*, or C-Δ*parR*. Larvae were injected with 10 μL bacterial suspension at 4 × 10³ CFU/mL (approximately 40 CFU per larva) and monitored over time. PBS-injected larvae served as a negative control. (C) RT-qPCR analysis of eight representative targets spanning resistance, biofilm, and virulence-associated modules in WT PA14, Δ*parR*, and C-Δ*parR*. Transcript levels were normalized to the internal reference gene and are shown relative to WT. The complete 20-gene panel is provided in Fig. S2. (D) Representative EMSAs showing ParR binding to selected promoter probes, including P*_opdL_*, P*_arnB_*, P*_oprD_*, and P*_mexY_*, across the indicated ParR concentrations (0-0.8 μM). Positions of free probe and ParR-DNA complexes are indicated. Additional EMSAs for the full promoter panel are shown in Fig. S3. (E) Promoter-*lacZ* reporter assays for the same representative promoter subset shown in (C), measured by β-galactosidase activity in the absence or presence of ParR expression. The full reporter panel is provided in Fig. S4. Data in (A), (C), and (E) are mean ± SD from three independent experiments. Statistical significance is indicated in the figure. One-way ANOVA with Dunnett’s multiple-comparisons test was used for (A), (C) and (E), and survival differences in (B) were analyzed by the log-rank test. ns, not significant; \**P* < 0.05, \*\**P* < 0.01, \*\*\**P* < 0.001, \*\*\*\**P* < 0.0001.

To determine whether these ParR-linked physiological shifts extend to host relevant outcomes, we assessed virulence in a *Galleria mellonella* infection model. Under a standardized inoculum, WT caused rapid larval mortality, whereas the Δ*parR* mutant showed significantly delayed killing (Fig. 1B). Complementation reversed this attenuation and restored a WT-like survival trajectory (Fig. 1B; log-rank test as indicated). Together with the antimicrobial and biofilm phenotypes, this attenuation prompted us to map the downstream regulatory network associated with the pleiotropic effects of ParR.

To place these phenotype-level shifts into a genome-wide context, we compared RNA-seq profiles of WT and Δ*parR* under matched growth conditions. Across 5,891 annotated genes, 76 met the preset significance cutoff (Fold Change > 1.5, FDR-adjusted *P* < 0.05; 72 upregulated and 4 downregulated in Δ*parR*) (Fig. S1A). The differentially expressed genes (DEGs) were consistent with the observed phenotypes. Altered expression of envelope-remodelling and permeability factors, including the carbapenem porin *oprD*(18), aligned with multidrug susceptibility shifts. Dysregulation of the T3SS regulator *exsA*(19) and various TCSs (*phoP, rssB, amgR*)(20) provided a molecular basis for the attenuated host killing. KEGG enrichment also detected repression of the pyochelin biosynthesis and uptake module (*pchA, pchE, pchF, fptA*)(21) (Fig. S1B).

Steady-state transcriptomics, however, captures both direct regulatory targets and indirect cascade effects. To define a tractable set of candidate direct ParR targets, we used an intersectional prioritization strategy. We cross-referenced our RNA-seq DEGs with established ParRS evidence streams, including an experimentally mapped TCS regulatory network(22), a prior ParRS transcriptome linking to quorum-sensing circuitry(16), and our host peptide stress model (9). This filtering strategy yielded a curated panel of 20 candidate genes spanning resistance, biofilm and virulence-associated axes for stepwise validation (Table 2).

**Table 2.**
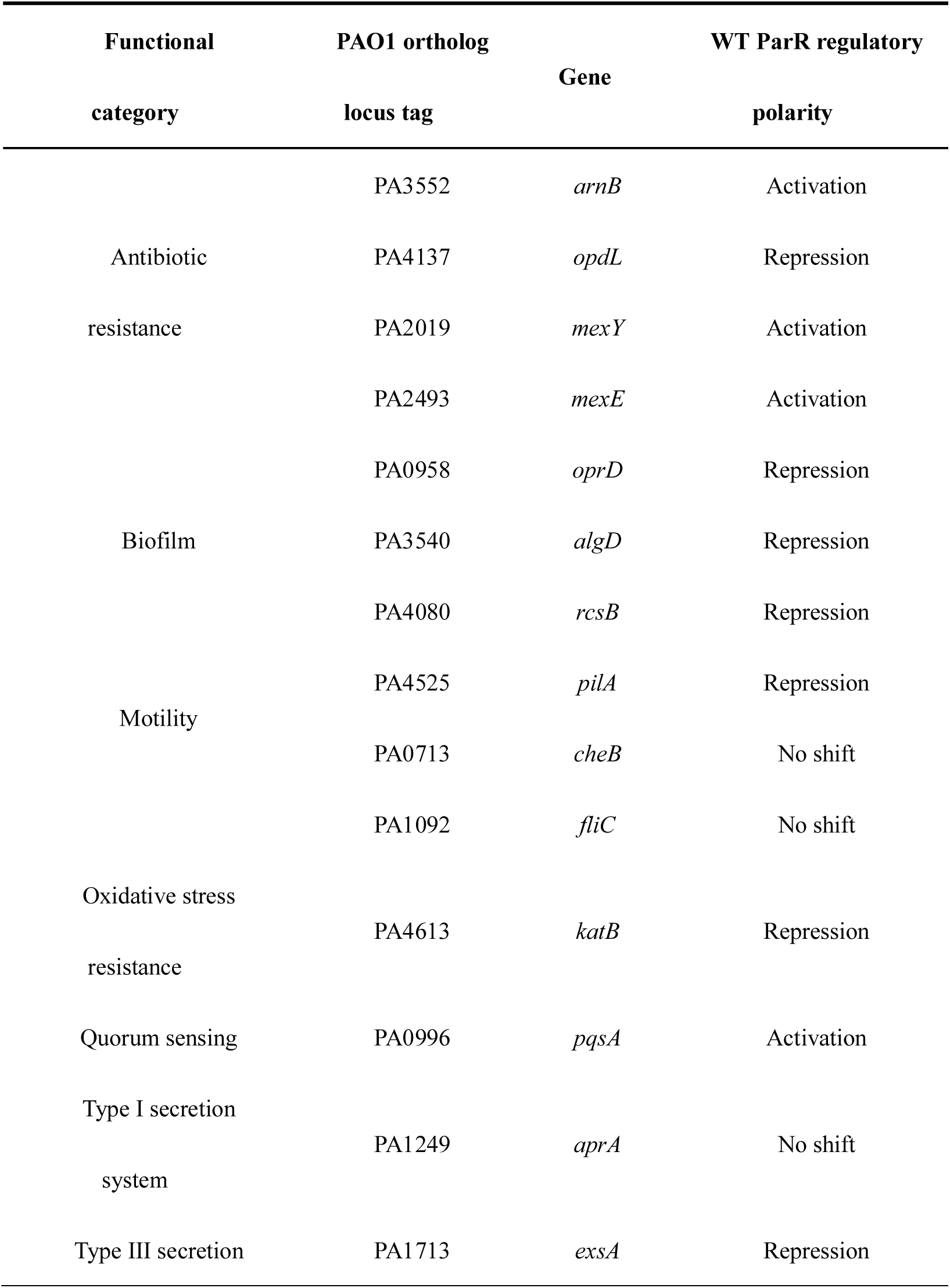
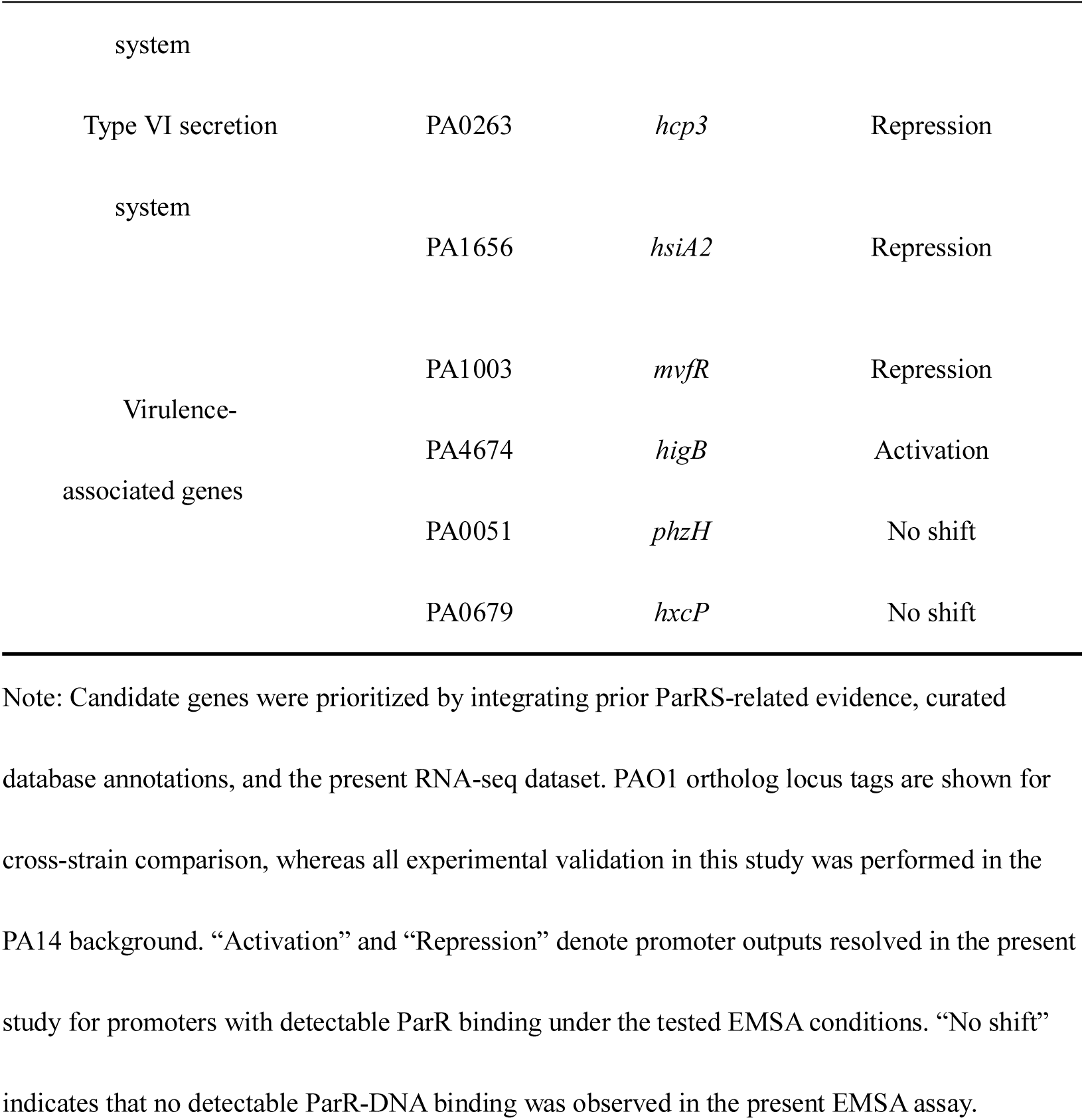
Curated candidate ParR-associated genes and their validation outcomes in the PA14 background.

Targeted RT-qPCR confirmed that *parR* deletion significantly altered the expression of 16 out of the 20 candidates, with complementation largely restoring WT like levels (Fig. S2). Eight representative targets are shown in Fig. 1C to summarize the major regulatory trends. Within the resistance module, Δ*parR* reduced the expression of the polymyxin-resistance determinant *arnB* (approximately 0.5-fold of WT) and the efflux-associated genes *mexY* and *mexE* (approximately 0.3-fold and 0.4-fold, respectively). By contrast, expression of the porins *oprD* and *opdL* increased (approximately 2.6-fold and 1.6-fold, respectively). Biofilm- and virulence-linked genes were also rewired in the mutant, including induction of *algD*, *rcsB* and *mvfR*, induction of secretion and stress-associated genes *exsA*, *hcp3*, *hsiA2*, *katB*, and *hxcP*, and reduced *pqsA*. Motility-associated genes *cheB*, *fliC*, the type I secretion linked gene *aprA* and the phenazine linked gene *phzH* remained unchanged, indicating selective ParR-dependent rewiring (Fig. S2).

We next asked whether ParR directly engages these loci at the promoter level. Using EMSA with approximately 200 bp promoter probes, ParR produced clear, concentration dependent DNA-protein complexes for 15 of the 20 targets (75%) (Table 3; Fig. 1D; Fig. S3). Binding affinities, estimated by the minimum ParR: DNA molar ratio required for a visible shift, varied widely. P*_opdL_* exhibited the strongest affinity (1:1 ratio), whereas most promoters, including P*_arnB_*, P*_algD_*, and P*_mvfR_* clustered at a 2:1 ratio. A smaller subset, such as P*_mexY_* and P*_oprD_*, required higher ratios (4:1 or 8:1) for detectable complex formation (Table 3). These EMSA data confirm broad, direct promoter engagement by ParR across diverse functional modules.

**Table 3.**
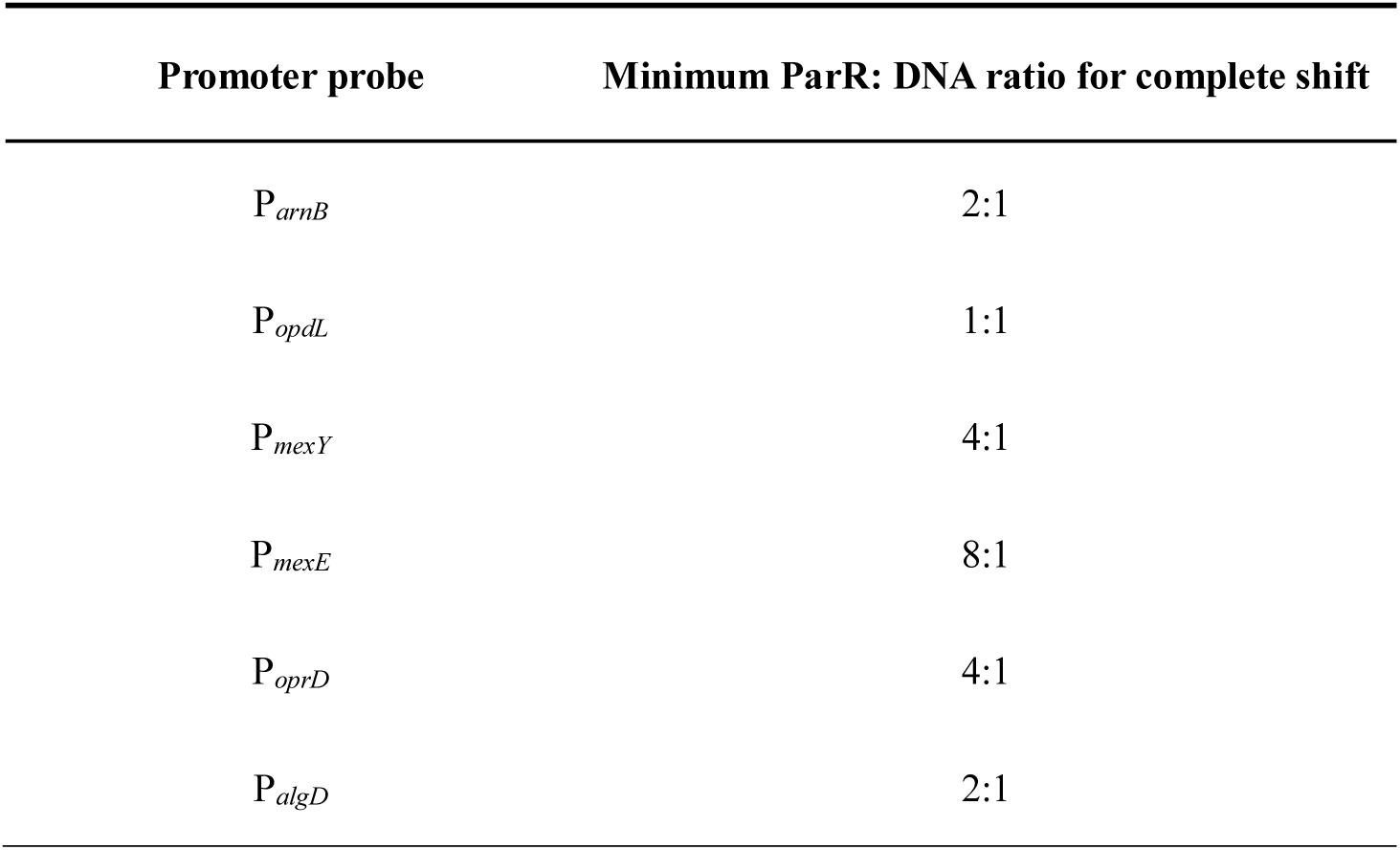
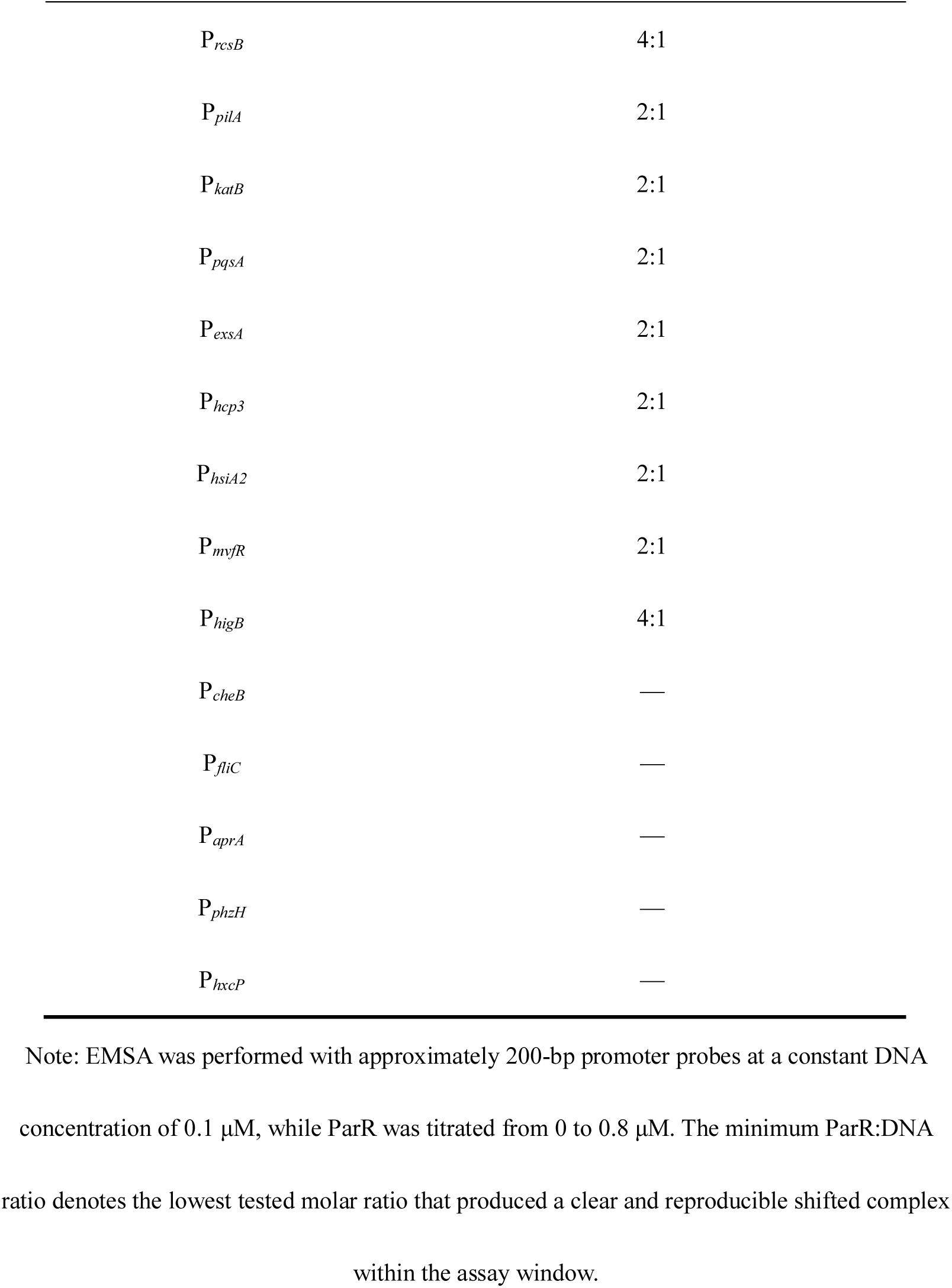
Apparent EMSA shift thresholds for WT ParR across the tested promoter probes.

Given this direct binding, we quantified the corresponding promoter outputs using a β-galactosidase reporter assay under ParR (+) versus ParR (−) conditions (Fig. 1E; Fig. S4). Direct ParR binding did not impose a uniform regulatory polarity. Instead, ParR activated a subset of targets, including P*_arnB_*, P*_mexY_*, and P*_pqsA_*, with 1.3-to 2.7-fold induction. it repressed other promoters, including P*_oprD_*, P*_algD_*, and P*_mvfR_*, reducing promoter activity to approximately 18-67% of basal ParR-free levels.

Together, these data identify ParR as a coordinator that directly engages a diverse promoter repertoire linked to multidrug resistance, biofilm formation and virulence-associated traits. The observation that one transcription factor can drive both activation and repression across its regulon raises a mechanistic question. Because TCS RR activity is commonly governed by phosphorylation state, we next asked whether bidirectional transcriptional outputs are hardwired into promoter sequences or are decoded dynamically by the phosphorylation status of ParR.

### 3.2. Target promoters decode ParR phosphorylation into distinct transcriptional logics

In canonical TCS, phosphorylation of the conserved receiver domain acts as an allosteric switch that reshapes receiver-domain conformational equilibria and dimerization, thereby altering promoter occupancy(23–25). Evidence that unphosphorylated RRs can exert distinct regulatory activities further supports phosphorylation state as a bidirectional determinant of output, rather than only a quantitative activity dial(26–28). To test whether phosphorylation dictates the bidirectional outputs observed across the ParR regulon, we focused on the conserved receiver-domain phosphoacceptor, Asp57 (D57) (Fig. 2A)(24).

**Fig. 2.**
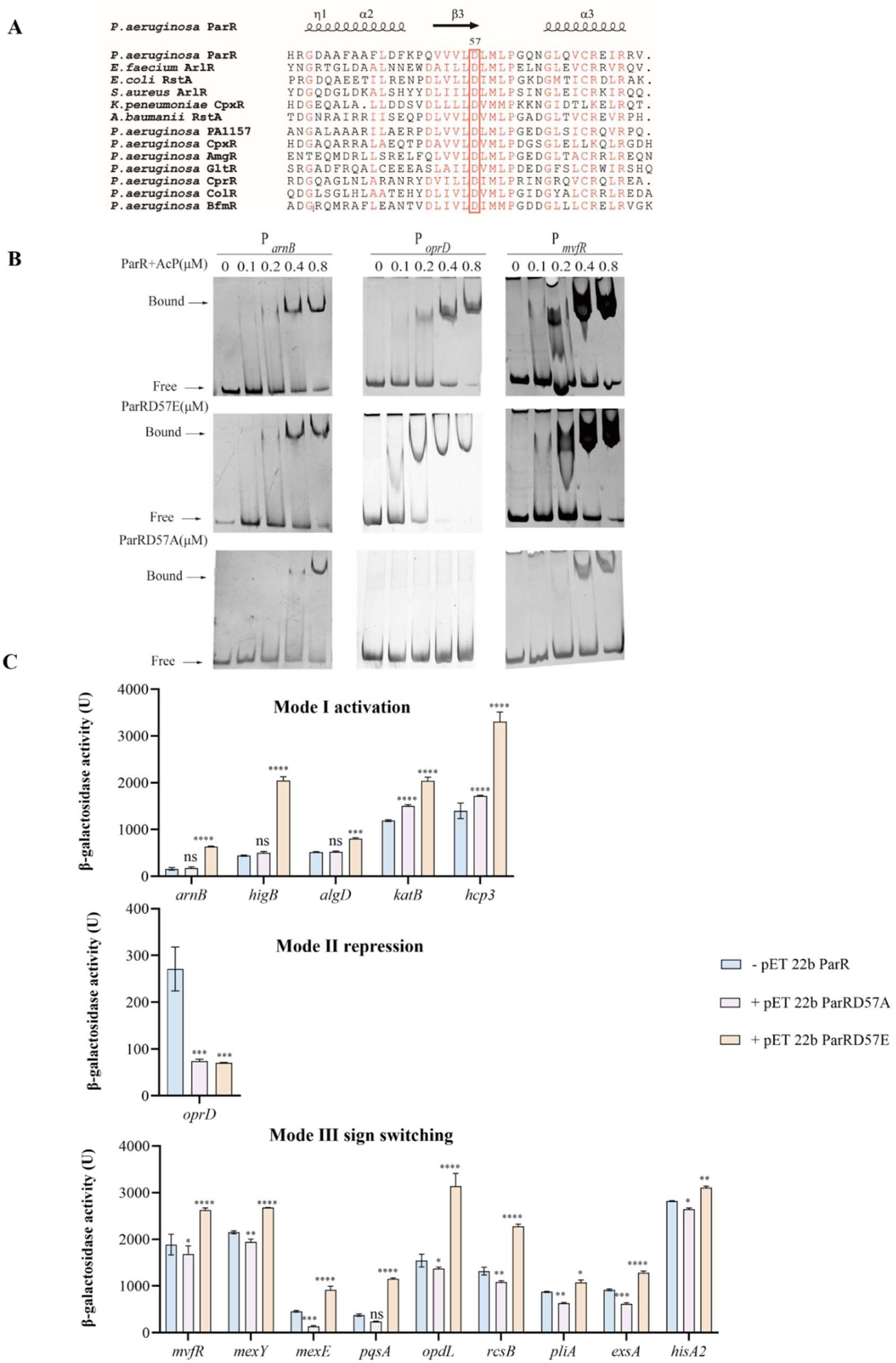
State-locked ParR D57 variants reveal promoter-specific decoding of phosphorylation-enhanced DNA binding. (A) Multiple-sequence alignment of *P. aeruginosa* ParR with representative RRs, highlighting the conserved receiver-domain phosphoacceptor Asp57 (red box). Secondary-structure elements of ParR are shown above the alignment. All aligned protein sequences were retrieved from UniProt. (B) Representative EMSAs showing binding of ParR+AcP, ParRD57E and ParRD57A to the indicated target promoters. Increasing protein concentrations (0, 0.1, 0.2, 0.4, and 0.8 μM) were incubated with the corresponding promoter fragments. Bound and free DNA species are indicated. These gels illustrate that AcP-treated WT ParR and the ParRD57E lower promoter occupancy thresholds, whereas ParRD57A weakens or abolishes binding in a promoter-dependent manner. (C) β-galactosidase activities of heterologous promoter lacZ reporters co-expressed with pET 22b ParR, pET 22b ParRD57A, or pET 22b ParRD57E. Promoters are grouped into three regulatory classes according to their transcriptional responses to the ParR receiver-domain state: Mode I, activation; Mode II, repression; and Mode III, sign switching. Bars show mean ± SD from three independent biological replicates. Statistical significance was assessed by one-way ANOVA followed by Dunnett’s multiple comparisons test within each promoter, using WT ParR as the reference group where indicated. Exact comparisons are annotated in the plot. ns, not significant; \**P* < 0.05, \*\**P* < 0.01, \*\*\**P* < 0.001, \*\*\*\**P* < 0.0001.

To uncouple ParR activation from dynamic upstream histidine kinase signalling, we generated genetically stable D57 substitution variants: ParRD57A as a nonphosphorylatable allele and ParRD57E as a commonly used phosphomimetic proxy(29, 30). Because Asp to Glu substitutions do not always perfectly recapitulate authentic phosphorylation, we benchmarked ParRD57E against wild-type ParR phosphorylated in vitro using acetyl phosphate (AcP), a standard small molecule phosphodonor(30, 31). This dual genetic and biochemical approach enables precise, promoter-resolved comparisons of ParR state dependent outputs.

We first mapped how the D57 state reshapes ParR-DNA occupancy across the 15 direct target promoters using EMSA. The D57 state altered occupancy thresholds in a promoter-dependent manner (representative gels in Fig. 2B; promoter-resolved summary in Table 4; remaining EMSAs in Fig. S5). Activated-state conditions universally lowered the binding threshold. ParR+AcP produced detectable shifts at a 1:1 minimum binding ratio for 11 promoters and at 2:1 for the remaining four promoters (P*_algD_*, P*_rcsB_*, P*_pilA_*, and P*_hcp3_*). The binding profile of ParRD57E was closely matched to that of ParR+AcP, shifting 12 promoters at 1:1 and three promoters (P*_algD_*, P*_rcsB_*, and P*_hcp3_*) at 2:1. This close agreement supports the use of ParRD57E as a practical activated-state proxy for downstream assays, with a minor difference at P*_pilA_*, where ParRD57E shifted at 1:1 whereas ParR+AcP required 2:1.

**Table 4.**
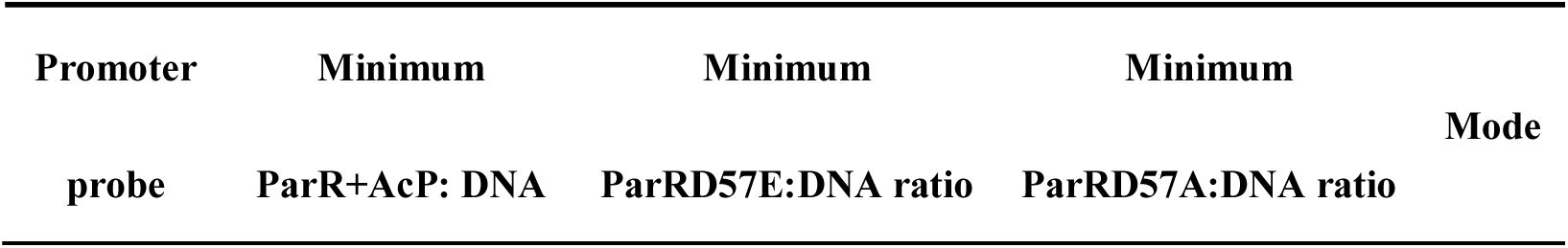
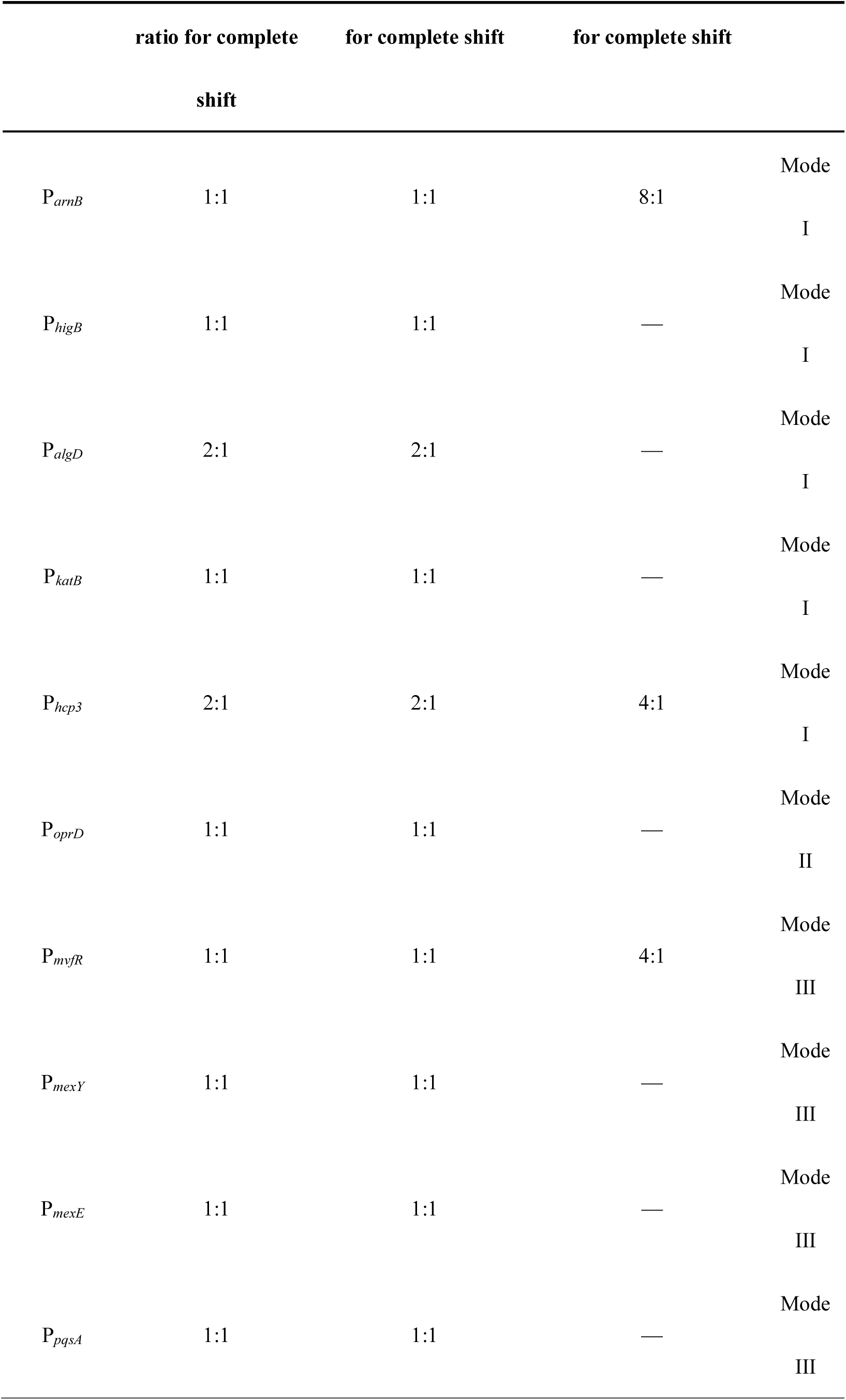
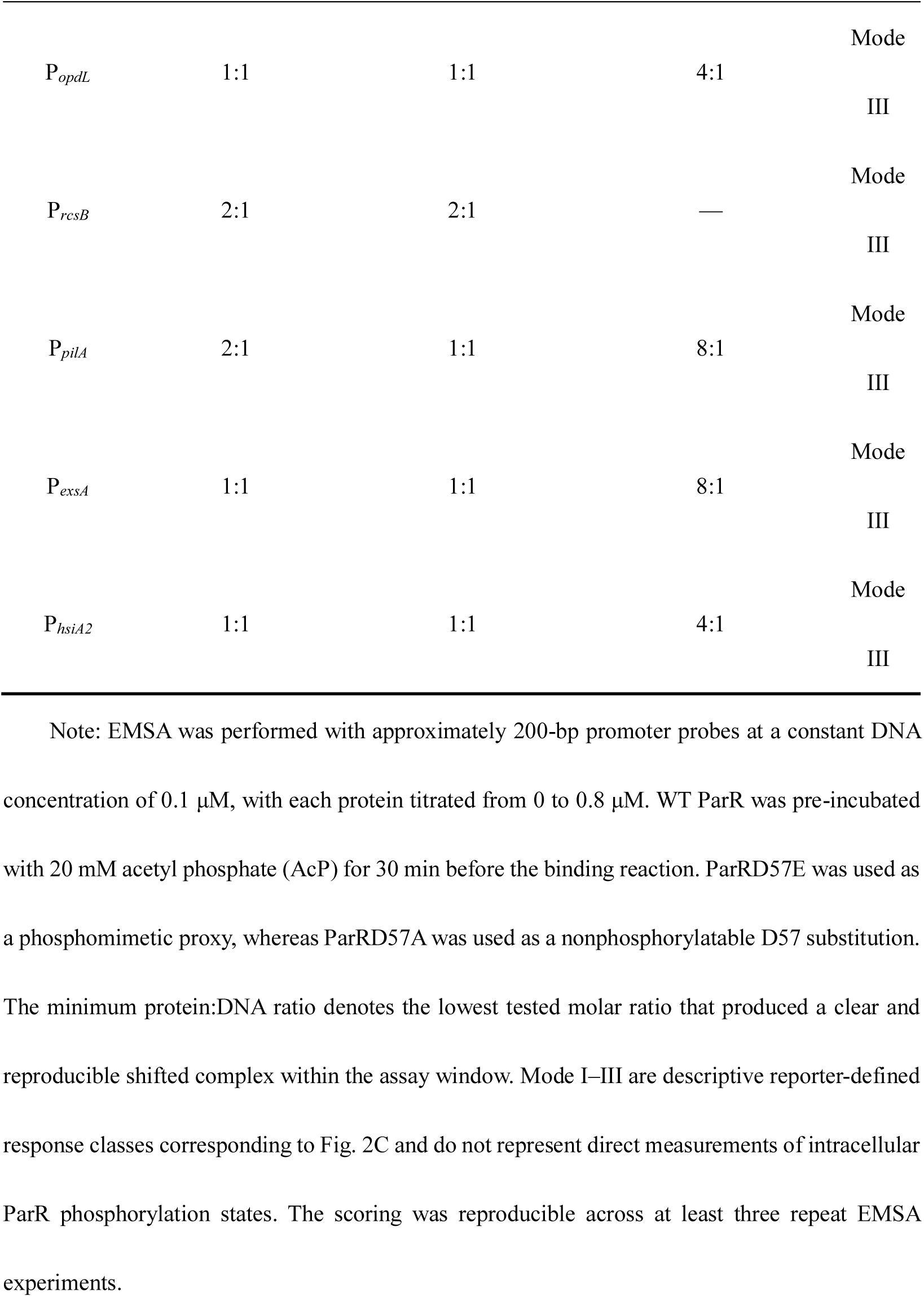
D57-state response classes of the candidate promoters determined using wild-type ParR, ParR D57A, and ParR D57E.

By contrast, the phosphorylation deficient proxy ParRD57A weakened or abolished binding and partitioned the promoters into three distinct regimes. Eight promoters, including P*_mexY_*, P*_mexE_*, P*_oprD_*, P*_algD_*, P*_rcsB_*, P*_katB_*, P*_pqsA_*, and P*_higB_*, showed no detectable shift within the tested concentration window. These promoters therefore require an activated receiver-domain state for stable occupancy under the assay conditions. Four promoters (P*_opdL_*, P*_mvfR,_* P*_hsiA2,_* P*_hcp3_*) retained measurable ParRD57A engagement only at an elevated threshold of 4:1, whereas three promoters (P*_arnB_*, P*_pilA_*, and P*_exsA_*) required an even higher threshold of 8:1.

These EMSA data show that phosphorylation enhances DNA binding by lowering occupancy thresholds, but this shared biophysical gain alone cannot explain how ParR activates some target genes and represses others. We hypothesized that individual promoters decode this phosphorylation signal into distinct transcriptional outputs(32). To test this, we quantified promoter activity using a heterologous promoter β-Galactosidase reporter assay, where each target promoter drives *lacZ* and ParR variants (WT, D57A, or D57E) are supplied *in trans* (Fig. 2C)(33). This configuration isolates promoter logic from native feedback loops.

Based on the sign and magnitude of transcriptional output across the phosphorylation-deficient (ParRD57A) and phosphomimetic (ParRD57E) states, we grouped the ParR regulon into three distinct regulatory modes (Fig. 2C). Mode I, phosphorylation-enhanced activation, comprises promoters for which ParR acts as an activator and phosphorylation increases this activation. The *arnB* promoter exemplified this behaviour: output increased markedly under ParRD57E relative to WT, whereas ParRD57A remained near basal levels. This pattern is consistent with phosphorylation increasing productive RNAP recruitment at activation-competent architectures(32). Mode II, phosphorylation-robust repression, is exemplified by the porin promoter *oprD*. These targets remained strongly repressed regardless of ParR state. Even weak binding by ParRD57A or basal WT levels was sufficient to maintain repression, suggesting a promoter configuration in which steric hindrance of RNAP dominates across a broad range of regulator phosphorylation levels(32). Mode III, phosphorylation-dependent sign switching, comprises promoters whose outputs invert between unphosphorylated and phosphorylated states. The virulence regulator *mvfR* illustrates this switching: it is repressed by the inactive ParRD57A but strongly activated by the phosphomimetic ParRD57E (Fig. 2C). Such sign switching indicates that D57 phosphorylation changes more than a quantitative activity dial and can alter the functional relationship between the RR and promoter machinery (34). Together, these mode definitions support a model in which ParR phosphorylation functions as a global state variable that is differentially decoded by local promoter architectures.

### 3.3 Chromosomal ParR D57 substitutions are associated with transcriptional and phenotypic differences

To determine whether the promoter decoding logic defined in the heterologous system operates at endogenous loci, we engineered the state-locked *parRD57A* and *parRD57E* alleles directly into the *P. aeruginosa* chromosome. This allele-based strategy enforces a homogeneous receiver-site condition across the cell population and reduces confounding from mixed phosphorylated and unphosphorylated ParR pools that can arise from variable kinase stoichiometry or host-derived modifications(35, 36).

Using these chromosomal D57 substitution strains, we first quantified representative direct target transcripts by RT-qPCR (Fig. 3A). The chromosomal expression patterns were broadly consistent with the three regulatory modes identified in the heterologous assays. Consistent with Mode I activation, ParRD57E produced the strongest induction across this subset, with *arnB* ncreasing by approximately 6.83-fold and *higB* by approximately 3.73-fold. *algD*, *katB*, and *hcp3* showed a similar phosphorylation-graded pattern (ParRD57A, approximately 1.59- to 1.65-fold; ParRD57E, approximately 1.90- to 2.71-fold) (Fig. 3A). Mode II repression was maintained at *oprD*, whose transcript abundance was suppressed in both ParRD57A (approximately 0.27-fold) and ParRD57E (approximately 0.30-fold) relative to WT (Fig. 3A). . Mode III sign switching was also reinforced at the mRNA level. As shown by *mvfR* (ParRD57A approximately 0.79-fold vs. ParRD57E approximately 2.39-fold), the Mode III set, including *mexY*, *mexE*, *pqsA*, *opdL*, *rcsB*, *pilA*, *exsA* and *hsiA2*, generally showed basal or repressed transcription in the D57A condition and induction in the D57E condition (Fig. 3A). These data support an association between RR phosphorylation state and derepression or sign switching at selected endogenous targets(37).

**Fig. 3.**
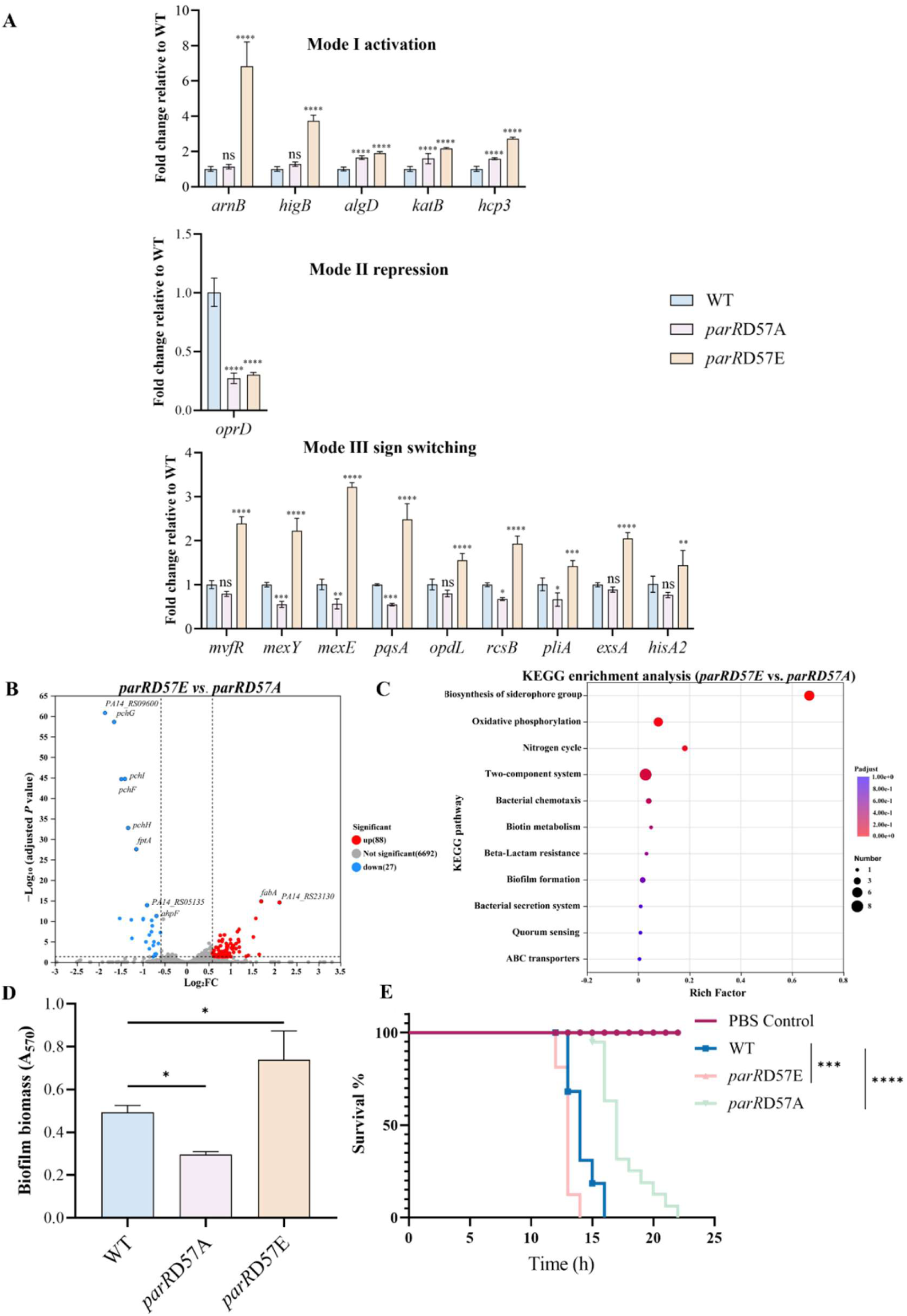
State-locked ParR phosphorylation rewires endogenous transcription and downstream phenotypes. (A) RT-qPCR analysis of representative direct targets in WT, *parRD57A* and *parRD57E* strains. Genes are grouped according to the three promoter decoding behaviors defined in the preceding analyses: Mode I, activation (*arnB, higB, algD, katB,* and *hcp3*); Mode II, repression (*oprD*); and Mode III, sign switching (*mvfR, mexY, mexE, pqsA, opdL, rcsB, pilA, exsA,* and *hsiA2*). Transcript levels are shown as fold change relative to WT. (B) Volcano plot showing differential gene expression between *parRD57E* and *parRD57A*. Each dot represents one gene. Significantly upregulated and downregulated genes are highlighted in red and blue, respectively, according to the predefined RNA-seq thresholds fold change > 1.5 and FDR-adjusted *P* < 0.05. (C) KEGG enrichment analysis of differentially expressed genes from the *parRD57E* versus *parRD57A* comparison. Dot size indicates gene number, and dot color indicates adjusted *P* value. (D) Static biofilm biomass measured by crystal violet staining (A_570_) in WT, *parRD57E* and *parRD57A* strains. (E) Survival of *Galleria mellonella* larvae following infection with WT, *parRD57A* and *parRD57E*. Larvae were injected with 10 μL bacterial suspension at an inoculum of 4 × 10³ CFU/mL (approximately 40 CFU per larva) and monitored over time. PBS-injected larvae served as a negative control. Data in (A) and (D) are mean ± SD from three independent experiments. Statistical significance is indicated in the figure. One-way ANOVA with Tukey’s multiple-comparisons test was used for (A) and (D), and survival differences in (E) were analyzed by the log-rank test. ns, not significant; \**P* < 0.05, \*\**P* < 0.01, \*\*\**P* < 0.001, \*\*\*\**P* < 0.0001.

We next expanded our view to the global transcriptome using RNA-seq. Compared with WT PA14, the state-locked mutants yielded relatively small sets of DEGs under stringent cutoffs (ParRD57A: 5 DEGs; ParRD57E: 12 DEGs) (Fig. S6). This pattern is consistent with the biophysics of wild-type TCS signalling. WT ParR is not a static entity, but a dynamic mixed-state ensemble that samples a continuum of phosphorylation levels set by endogenous environmental cues. The WT transcriptome therefore represents an average over mixed states, which can compress statistical fold changes when either locked extreme is used as the comparator.

To resolve the regulatory amplitude of the ParR regulon, we directly compared the two homogeneous extremes (*parRD57E* versus *parRD57A*). This contrast increased transcriptional separation and revealed a broader set of differentially expressed genes (115 DEGs; 88 upregulated and 27 downregulated) (Fig. 3B). KEGG enrichment highlighted suppression of the pyochelin module (*pchF-I,* and *fptA*) and multiple chaperone or stress genes (*grpE, htpG, clpB*), together with the induction of respiratory or energy-metabolism nodes (*cioA, ccoO, napA, sdhC*) and a T6SS sheath gene (*tssB*). This broader transcriptional divergence indicates that altering the D57 state is associated with substantial changes in cellular transcriptional programs (Fig. 3C), prompting us to test whether these locked-state differences translate into phenotypic outcomes.

If ParR phosphorylation functions as a global state variable for transcriptional reprogramming, toggling this state should affect macroscopic survival phenotypes. Chromosomal D57 mutations bidirectionally shifted antibiotic tolerance (Table 5). Relative to PA14, *parRD57A* sensitized bacteria and lowered MICs by 2- to 4-fold across polymyxins, aminoglycosides and β-lactams. Conversely, *parRD57E* increased MICs up to 8-fold relative to *parRD57A*. This class-spanning resistance profile is consistent with the Mode I transcriptional rewiring of *arnB* activation described above(8).

**Table 5.** Minimum inhibitory concentrations (MICs) of WT, *parRD57A*, and *parRD57E* strains in Mueller-Hinton (MH) medium.

| Strain | Polymyxin B | Colistin | Tobramycin | Amikacin | Meropenem | Ceftazidime |
| --- | --- | --- | --- | --- | --- | --- |
| WT | 4 | 4 | 2 | 4 | 2 | 16 |
| *parRD57A* | 1 | 2 | 1 | 1 | 1 | 4 |
| *parRD57E* | 8 | 8 | 4 | 8 | 4 | 32 |

Note: MIC values are reported in $\mu\text{g/mL}$ . WT, wild type; *parRD57A*, nonphosphorylatable chromosomal ParR mutant; *parRD57E*, phosphomimetic chromosomal ParR mutant. MICs were determined by broth microdilution in Mueller-Hinton broth as described in Materials and Methods, with at least three independent biological replicates.

State-dependent stratification also extended to multicellular and host-pathogen interactions. In static biofilm assays, biomass tracked with the D57 allele: *parRD57E* accumulated significantly more biofilm than WT, whereas *parRD57A* formed less (Fig. 3D). This trend was consistent with state-biased induction of biofilm-associated nodes such as *algD* and *rcsB*. In a *Galleria mellonella* infection model, the locked states resolved into a clear killing hierarchy (Fig. 3E). *parRD57E* caused faster killing (median survival, 13 h) than WT (14 h), whereas *parRD57A* was attenuated, delaying death onset and extending survival (17 h).

Collectively, these data support a model in which ParR phosphorylation acts as a regulatory state variable whose effects are resolved differently across the regulon, with measurable consequences for antibiotic susceptibility, biofilm formation and host-associated virulence.

### 3.4 Divergent promoter architectures decode a shared ParR binding interface into distinct transcriptional logics

Our data revealed a mechanistic paradox: ParR phosphorylation enhances DNA binding broadly, yet this shared biophysical event is translated into activation, repression, or sign switching. In *E. coli*, quantitative models suggest that transcriptional fold changes are strongly shaped by basal promoter architecture (12). Because ParR belongs to the OmpR/PhoB family, in which DNA recognition often relies on paired half-sites separated by a spacer(38, 39), we hypothesized that the spatial arrangement of these *cis* elements encodes the regulatory logic. To test this, we selected one representative target from each regulatory mode, *arnB* for Mode I, *oprD* for Mode II and *mvfR* for Mode III, and used a stepwise pipeline combining de novo motif discovery (MEME Suite), EMSA mapping and minimal-box reporter assays to identify the ParR-binding region driving each output.

For the Mode I exemplar *arnB*, MEME analysis and tiled EMSA identified a single dominant binding region (P*_arnB-2_*) (Fig. 4A-D; Fig. S7). Sequence analysis revealed a compact, dyad-symmetric architecture consisting of two 8-bp inverted half-sites (TTGTTAAG) separated by a 5-bp spacer (Fig. 4B). When cloned into the reporter assays, this isolated element recapitulated Mode I behaviour: WT ParR activated transcription (approximately 1.9-fold), ParRD57E further increased activation (approximately 3.7-fold), and ParRD57A failed to activate (Fig. 4D). These results identify the compact symmetric half-site as an architecture compatible with phosphorylation-amplified activation.

**Fig. 4.**
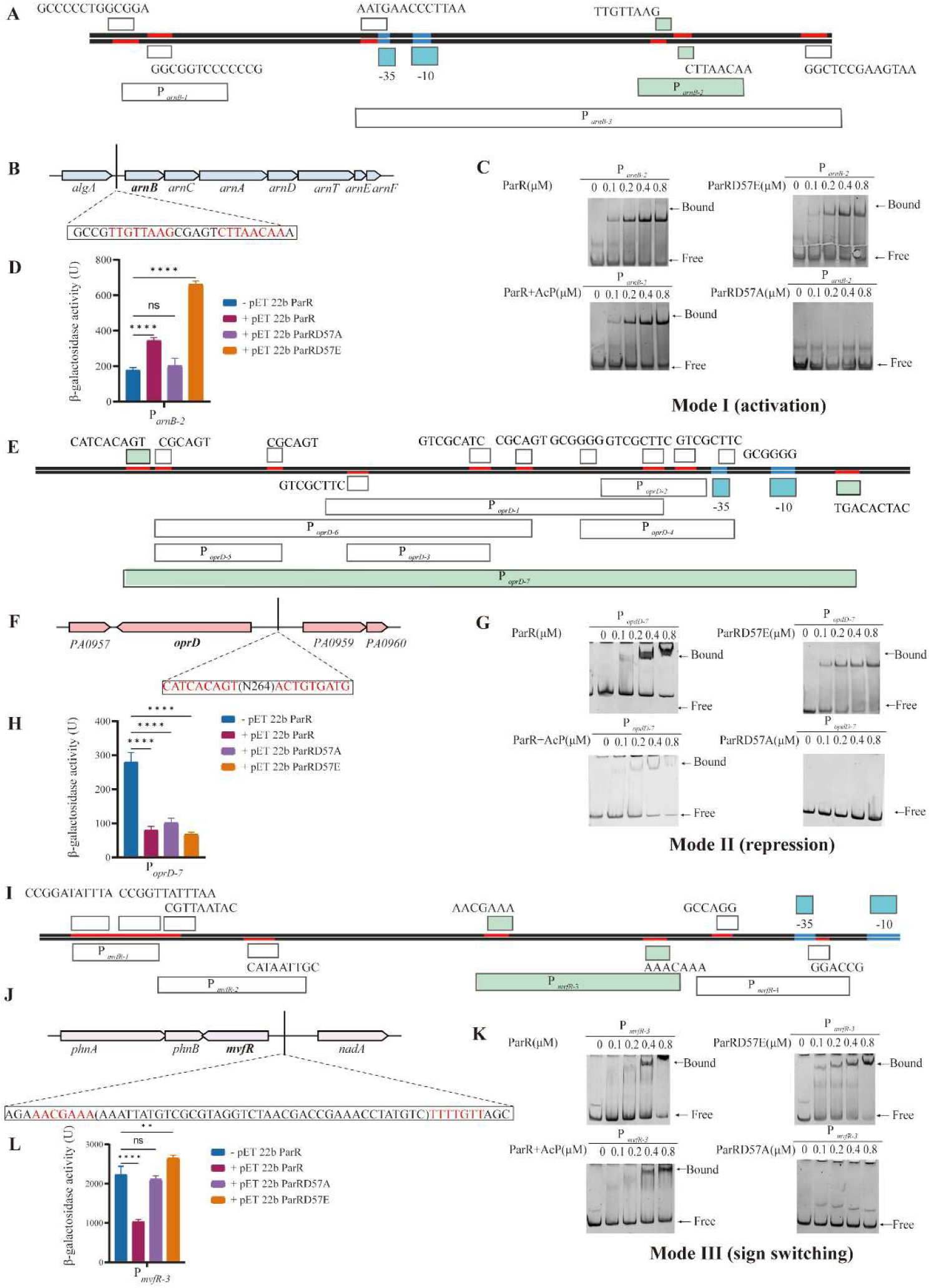
**Representative ParR boxes from *arnB*, *mvfR*, and *oprD* reveal architecture-dependent transcriptional outputs**. (A-D) Identification and functional validation of the *arnB* ParR binding site. (A) Schematic of the P*_arnB_* promoter region showing candidate sequence elements, core promoter features, and promoter fragments used for mapping; the validated ParR binding site, P*_arnB_*_-2_, is highlighted. (B) Genomic context of the *arn* locus and sequence of P*_arnB-2_*, with the putative paired half-sites highlighted in red. (C) EMSAs of P*_arnB_*_-2_ with ParR, ParRD57E, ParR+AcP and ParRD57A at the indicated protein concentrations. Bound and free DNA are indicated. (D) β-galactosidase activities of the minimal P*_arnB-2_* reporter in the indicated expression backgrounds, showing Mode I output (activation). (E-H) Identification and functional validation of the *oprD* ParR binding site. (E) Schematic of the *oprD* promoter region showing candidate sequence elements and promoter fragments used for mapping; the validated ParR binding site, P*_oprD-7_*, is highlighted. (F) Genomic context of the *oprD* locus and sequence of P*_oprD-7_*, with the putative paired half-sites highlighted in red and the long intervening spacer indicated. (G) EMSAs of P*_oprD_*_-7_ with ParR, ParRD57E, ParR+AcP, and ParRD57A at the indicated protein concentrations. Bound and free DNA are indicated. (H) β-galactosidase activities of the minimal P*_oprD_*_-7_ reporter in the indicated expression backgrounds, showing Mode II output (repression). (I-L) Identification and functional validation of the *mvfR* ParR binding site. (I) Schematic of the *mvfR* promoter region showing candidate sequence elements and promoter fragments used for mapping; the validated ParR binding site, P*_mvfR-3_*, is highlighted. (J) Genomic context of the *mvfR* locus and sequence of P*_mvfR-3_*, with the putative paired half-sites highlighted in red and the intervening spacer indicated. (K) EMSAs of P*_mvfR_*_-3_ with ParR, ParRD57E, ParR+AcP, and ParRD57A at the indicated protein concentrations. Bound and free DNA are indicated. (L) β-galactosidase activities of the minimal P*_mvfR_*_-3_ reporter in the indicated expression backgrounds, showing Mode III output (sign switching). Together, these representative ParR binding sites illustrate that distinct *cis*-architectures are associated with different transcriptional response patterns in the reporter assay framework used here. Bars represent mean ±SD from three biologically independent experiments. Statistical significance is indicated as shown in the plots; \**P* < 0.05, \*\**P* < 0.01, \*\*\**P* < 0.001, \*\*\*\**P* < 0.0001.

The ParR binding sites for Mode II and Mode III loci showed different spatial geometries. For the Mode II exemplar *oprD*, its mapped ParR binding site P*_oprD-7_* consists of an imperfect inverted-repeat core (CATCACAGT) separated by a 264-bp spacer (Fig. 4E-H; Fig. S8). Despite this long separation, the minimal element maintained repression across ParR phosphorylation states (Fig. 4G). For the Mode III exemplar *mvfR*, mapping identified P*_mvfR-3_*, which contains a conserved half-site core (AACGAAA) and a 39-bp spacer (Fig. 4I-L; Fig. S9). This isolated binding site as sufficient to support phosphorylation-dependent recoding: WT ParR repressed the reporter (approximately 2.1-fold), whereas ParRD57E shifted the output towards activation (approximately 1.3-fold) (Fig. 4L). Together, these data reveal an architectural stratification. Although the targets share a paired half-site scaffold, spacer length and symmetry are associated with how the phosphorylation signal is interpreted(40, 41).

This architectural diversity raised the question of whether ParR uses distinct protein conformations or cryptic domains to engage structurally divergent promoters. To test this, we performed structure-guided mutagenesis on the ParR DNA-binding domain (DBD). Using an AlphaFold3-predicted structure, we identified conserved residues on the putative recognition helix and wing (S202, L204, R205, K207, and L208) among OmpR/PhoB-family RRs (Fig. S10A-C). Alanine substitution of any single residue in this interface abolished detectable EMSA binding across all three structurally diverse ParR binding site (P*_arnB-2_*, P*_mvfR-3_*, and P*_oprD-7_*) (Fig. S10 D-F). Consistently, these DBD mutants completely lost regulatory activity in the corresponding reporter assays, collapsing outputs to the empty-vector baseline (Fig. S10 G).

These mutagenesis results indicate that representative promoters from distinct regulatory classes rely on a common ParR DNA-contact surface. The divergent transcriptional outputs are therefore unlikely to arise from alternative protein-DNA binding modes. Instead, they are decoded by promoter *cis* geometry. By varying half-site symmetry and spacer length, the genome can wire one phosphorylation input into distinct logic gates and thereby support differential execution of survival-associated programmes.

### 3.5 A two-tier *cis*-regulatory code programs the transcriptional logic of ParR

Our data point to a bipartite *cis-*regulatory code governing the ParR regulon: half-site sequence gates initial binding compatibility, whereas spacer geometry sets the topological constraints that decode phosphorylation into specific transcriptional behaviours. To test this model, we engineered synthetic ParR binding site by varying half-site composition and spacer length across the three exemplars, P*_arnB_*, P*_mvfR_*, and P*_oprD_*. We then asked whether minimal promoter edits were sufficient to reprogram transcriptional logic.

We first examined the role of the half-site core as a binding-compatibility gate. Scrambling the palindromic sequences in the three exemplars (P*_arnB-2-scr8_*, P*_mvfR-3-scr7_*, and P*_oprD-7-scr9_*) abolished detectable ParR engagement in vitro across all phosphorylation states (Fig. 5A). Thus, D57 phosphorylation lowers the occupancy threshold but cannot bypass the requirement for sequence compatibility.

**Fig. 5.**
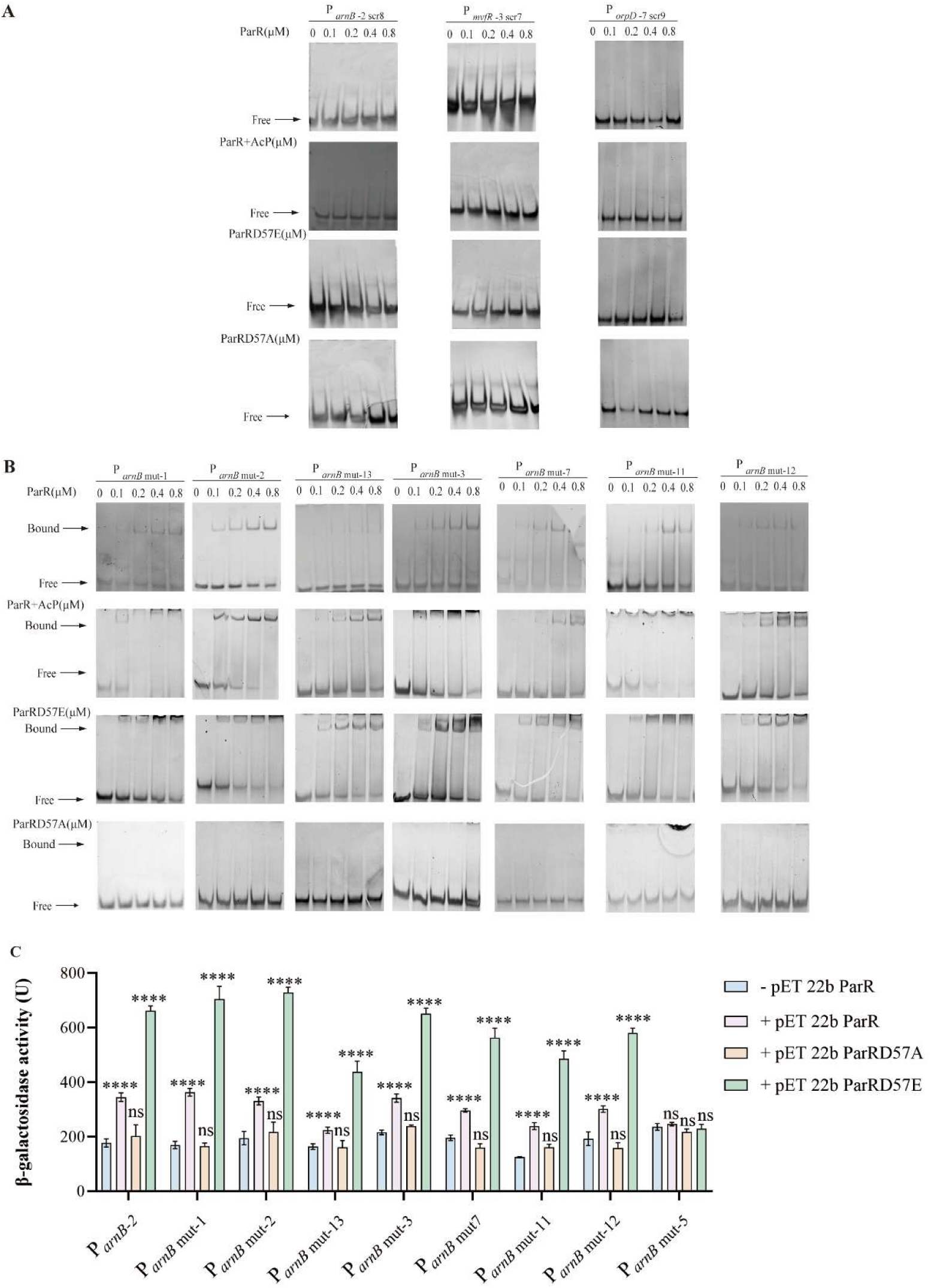
Half-site compatibility defines the first *cis*-regulatory gate for ParR occupancy and transcriptional output. (A) EMSAs using scrambled derivatives of three representative probes, P*_arnB-2-scr8_, P_mvfR_*_-3-*scr7*_, and P*_oprD-7-scr9_,* incubated with increasing concentrations (0, 0.1, 0.2, 0.4, and 0.8 μM) of ParR, ParR+AcP, ParRD57E, or ParRD57A. Scrambling the half-site sequence abolished detectable discrete binding across all tested ParR states. (B) EMSAs of selected nucleotide variants derived from the high-affinity P*_arnB-2_* probe. The indicated mutant probes were incubated with the same protein series as in panel A. Most perturbations weakened binding or increased the minimum protein: DNA ratio required for detection of a discrete shifted complex, whereas binding-permissive variants retained the state hierarchy described in the main text. Mutant definitions and full probe sequences are provided in Table 6 and Fig. S11. Bound and free DNA species are indicated. (C) β-Galactosidase reporter assay of the WT P*_arnB-2_* constructed and selected half-site variants in the presence of the indicated control or ParR state expression construct. Variants that remained binding-permissive in EMSA retained activation by ParR, especially ParRD57E, whereas the binding-defective variant P*_arnB-mut5_* collapsed toward baseline activity across all tested states. Data are mean ± SD (n ≥ 3). Statistical analysis: one-way ANOVA with Dunnett’s post hoc test relative to −pET 22b ParR. ns, no significant, \**P* < 0.05, \*\**P* < 0.01, \*\*\**P* < 0.001, \*\*\*\**P* < 0.0001.

**Table 6.**
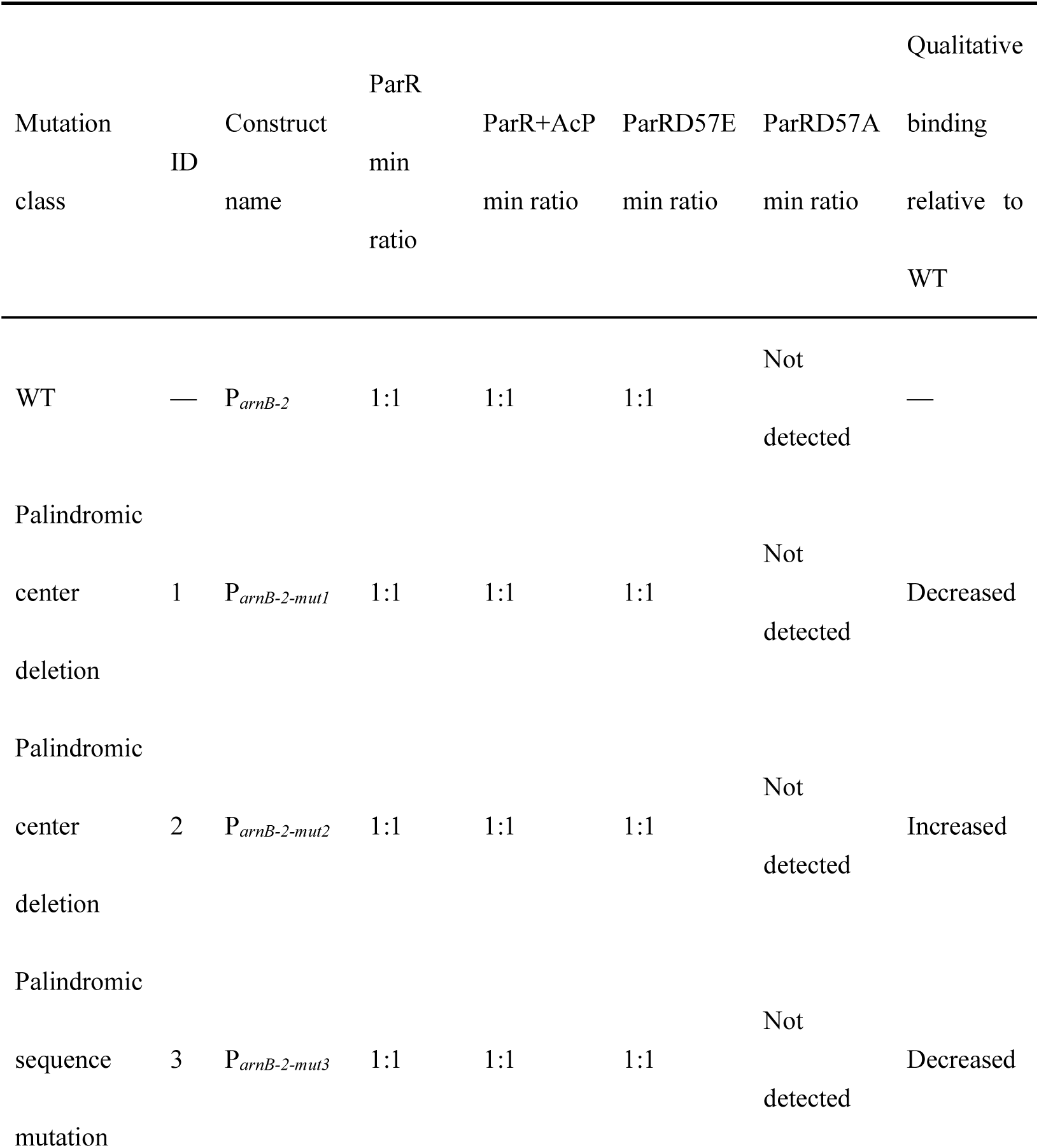
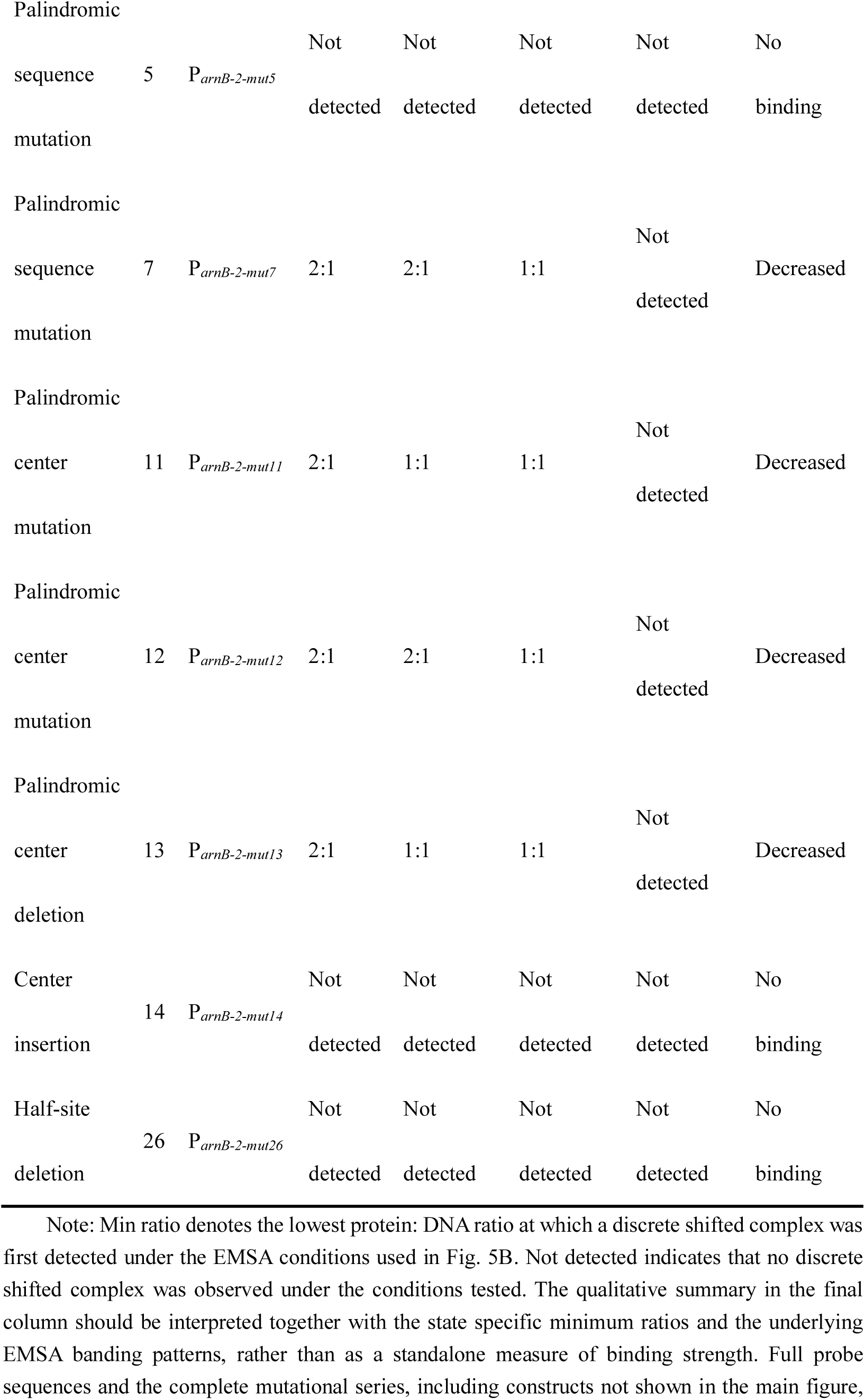
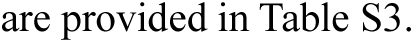
Representative nucleotide perturbations within P*_arnB-2_* define sequence compatibility and state-dependent EMSA occupancy thresholds.

| Mutation<br>class | ID | Construct<br>name | ParR | ParR+AcP | ParRD57E | ParRD57A | Qualitative<br>binding<br>relative to<br>WT |
| --- | --- | --- | --- | --- | --- | --- | --- |
|  |  |  | min<br>ratio | min ratio | min ratio | min ratio |  |
|  |  |  | ratio |  |  |  |  |
| WT | — | $P_{arnB-2}$ | 1:1 | 1:1 | 1:1 | Not<br>detected | — |
| Palindromic<br>center<br>deletion | 1 | $P_{arnB-2-mut1}$ | 1:1 | 1:1 | 1:1 | Not<br>detected | Decreased |
| Palindromic<br>center<br>deletion | 2 | $P_{arnB-2-mut2}$ | 1:1 | 1:1 | 1:1 | Not<br>detected | Increased |
| Palindromic<br>sequence<br>mutation | 3 | $P_{arnB-2-mut3}$ | 1:1 | 1:1 | 1:1 | Not<br>detected | Decreased |
| Palindromic |  |  |  |  |  |  |  |
| sequence | 5 | *P<sub>arnB-2-mut5</sub>* | Not detected | Not detected | Not detected | Not detected | No binding |
| mutation |  |  |  |  |  |  |  |
| Palindromic |  |  |  |  |  |  |  |
| sequence | 7 | *P<sub>arnB-2-mut7</sub>* | 2:1 | 2:1 | 1:1 | Not detected | Decreased |
| mutation |  |  |  |  |  |  |  |
| Palindromic |  |  |  |  |  |  |  |
| center | 11 | *P<sub>arnB-2-mut11</sub>* | 2:1 | 1:1 | 1:1 | Not detected | Decreased |
| mutation |  |  |  |  |  |  |  |
| Palindromic |  |  |  |  |  |  |  |
| center | 12 | *P<sub>arnB-2-mut12</sub>* | 2:1 | 2:1 | 1:1 | Not detected | Decreased |
| mutation |  |  |  |  |  |  |  |
| Palindromic |  |  |  |  |  |  |  |
| center | 13 | *P<sub>arnB-2-mut13</sub>* | 2:1 | 1:1 | 1:1 | Not detected | Decreased |
| deletion |  |  |  |  |  |  |  |
| Center |  |  |  |  |  |  |  |
| insertion | 14 | *P<sub>arnB-2-mut14</sub>* | Not detected | Not detected | Not detected | Not detected | No binding |
| Half-site |  |  |  |  |  |  |  |
| deletion | 26 | *P<sub>arnB-2-mut26</sub>* | Not detected | Not detected | Not detected | Not detected | No binding |

Note: Min ratio denotes the lowest protein: DNA ratio at which a discrete shifted complex was first detected under the EMSA conditions used in Fig. 5B. Not detected indicates that no discrete shifted complex was observed under the conditions tested. The qualitative summary in the final column should be interpreted together with the state specific minimum ratios and the underlying EMSA banding patterns, rather than as a standalone measure of binding strength. Full probe sequences and the complete mutational series, including constructs not shown in the main figure,
are provided in Table S3.

To define the boundaries of this compatibility, we performed systematic mutagenesis on the high-affinity P*_arnB-2_* scaffold (Fig. 5B; Table 6; Table S3; Fig. S11). Most single nucleotide variations within the half-site eliminated discrete complex formation. A small subset of mutations, including mut7/11/12/13, retained binding at elevated thresholds while preserving the state-dependent hierarchy (ParRD57E ≈ ParR+AcP > WT > ParRD57A). These biophysical constraints were recapitulated in reporter assays. Binding-permissive variants preserved Mode I activation, whereas half-site-disrupting mutations such as mut5 abolished transcriptional activity across all ParR states and reduced output to baseline levels (Fig. 5C). Together, these data identify half-site compatibility as the first essential gate for regulon entry.

We next tested whether spacer geometry determines the input-output transfer function. To do so, we performed spacer-swap engineering by exchanging spacer lengths among the *arnB* (5-bp), *mvfR* (39-bp) and *oprD* (264-bp) scaffolds (Fig. 6A). This design allowed us to ask whether a promoter could be shifted into a different regulatory mode by altering spacer length alone.

**Fig. 6.**
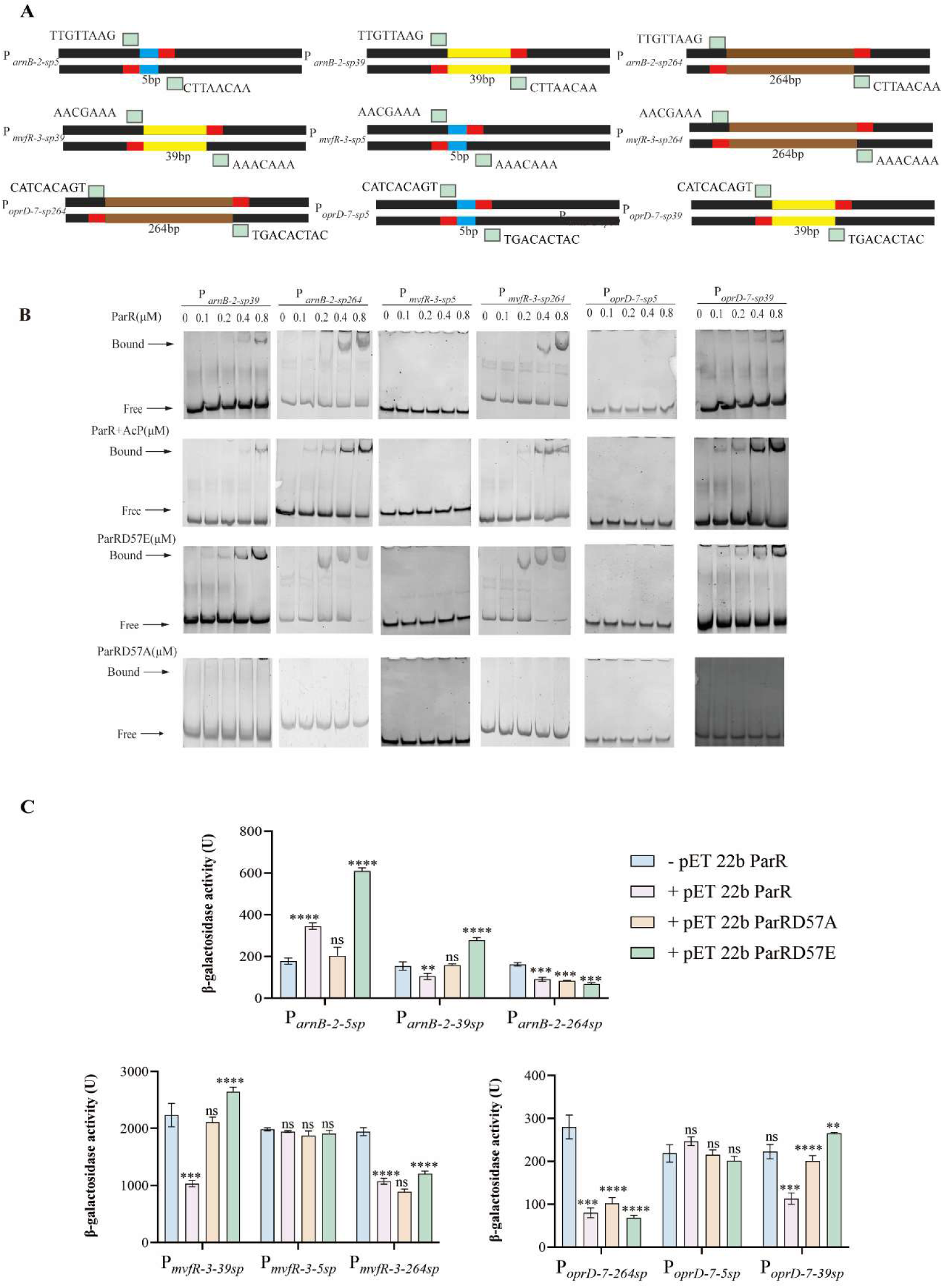
Spacer length reprograms ParR transcriptional output. (A) Schematic of spacer-swap constructs generated by exchanging the spacer segment between the paired half-sites of three representative ParR binding sites, P*_arnB-2_, P_mvfR_*_-3_, and P*_oprD_*_-7_. Native and engineered constructs carrying 5-bp, 39-bp, or 264-bp spacers are shown. Spacer modules are color-coded as follows: blue, 5-bp; yellow, 39-bp; brown, 264-bp. (B) EMSAs of representative spacer-swap probes incubated with increasing concentrations (0, 0.1, 0.2, 0.4, and 0.8 μM) of ParR, ParR+AcP, ParRD57E, or ParRD57A. Spacer expansion in the *arnB* scaffold increased the WT binding threshold while preserving preferential binding by the activated states, whereas compression of the *mvfR* and *oprD* architectures to a 5-bp spacer abolished detectable binding. Bound and free DNA species are indicated. (C) β-galactosidase reporter activities of native and spacer-swapped promoter constructs in the presence of the indicated control or ParR-state expression construct. In the *arnB* scaffold, spacer extension rewired the native activation mode into sign switching or repression. In the *oprD* scaffold, introduction of an intermediate 39-bp spacer unlocked phosphorylation-dependent sign switching from a natively repressed architecture. Each scaffold is plotted on an independent y-axis scale. Data are mean ± SD (n=3). Statistical analysis: one-way ANOVA with Dunnett’s post hoc test relative to −pET 22b-ParR. ns, no significant, \**P* < 0.05, \*\**P* < 0.01, \*\*\**P* < 0.001, \*\*\*\**P* < 0.0001.

In EMSA, spacer alteration shifted binding thresholds substantially (Fig. 6B). Expanding the *arnB* spacer to 39 bp or 264 bp increased the WT ParR binding threshold (≥2:1) while maintaining high apparent affinity for activated states (approximately 1:1). Conversely, compressing the *mvfR* and *oprD* spacers to 5 bp abolished detectable binding, suggesting topological constraints that phosphorylation could not overcome.

The reporter assays showed that spacer swapping could reprogram regulatory modes. In the *arnB* scaffold, extending the spacer from 5 bp to 39 bp converted native Mode I activation into Mode III sign switching, whereas extension to 264 bp shifted the output towards Mode II repression. Similarly, grafting an intermediate 39-bp spacer onto the *oprD* architecture produced phosphorylation-dependent sign switching in a promoter that is natively repressed. These results show that spacer geometry can flatten, amplify or invert the transcriptional readout of a constant phosphorylation signal (Fig. 6C).

Together, these data connect promoter-level binding properties with transcriptional and phenotypic outputs in the systems examined here. The ParR regulon operates through a transfer function in which the regulator provides a state-dependent binding gain, whereas promoter architecture acts as the locus-specific logic gate that shapes the final output. This modular *cis*-decoding mechanism explains the pleiotropic control of native bacterial virulence and resistance programmes and offers a framework for synthetic promoter design. By tuning spacer lengths and half-site affinities, minimal promoter edits can reprogram complex input-output behaviours without modifying the trans-acting regulator(41, 42).

## 4. Discussion

Bacteria must balance the metabolic costs of virulence with the survival demands imposed by antibiotic exposure (8, 15, 16). The *P. aeruginosa* ParRS TCS provides a model for this multidimensional control, but how one sensory kinase-RR pair orchestrates pleiotropic reprogramming has remained unclear. Our findings show that ParR does not function as a single uniform transcriptional switch. Instead, it occupies a diverse promoter network that includes *arnB* for envelope charge remodelling, *oprD* for porin-mediated influx and *mvfR* as a virulence regulator, and it exerts locus-specific regulatory effects with different signs(43).

A central biophysical insight of this work is that receiver-domain phosphorylation at D57 functions as a global state variable rather than a simple rheostat. Phosphorylation enhances ParR-DNA binding affinity and supports robust complex formation at lower regulator concentrations, but local promoter architecture determines the transcriptional consequence (26, 32). By separating binding from activation, we show that promoters can translate a common biophysical change, increased occupancy, into stronger activation, stable repression or a switch in regulatory sign. This is consistent with emerging structural paradigms for OmpR/PhoB-family regulators, in which the topological register between the RR dimer and RNA polymerase holoenzyme influences regulatory sign(44, 45).

We distill this mechanism into a two-tier *cis-regulatory* code: half-site sequence gates binding compatibility, whereas spacer geometry programs regulatory logic. In this framework, phosphorylation supplies the thermodynamic driving force, but promoter architecture defines the exact regulatory trajectory. Previous studies have highlighted the importance of *cis* context in bacterial transcription(12, 46, 47). Our spacer-swap experiments add causal support by showing that expanding or compressing the distance between half-sites can shift promoters across functional regimes, including activation, sign switching and repression. On this basis, we propose a working model in which ParR phosphorylation provides a common regulatory input, whereas promoter half-site compatibility and spacer architecture decode this input into distinct transcriptional and phenotypic outputs. These results indicate that modest topological rewrites can be sufficient to invert how a shared phosphorylation signal is interpreted in this experimental framework(48).

From an evolutionary perspective, architecture-driven decoding may provide one route by which bacteria diversify output without expanding the number of upstream regulators(49). Sequence comparisons suggest that this decoding framework may extend beyond PA14. Among representative *Pseudomonas* ParR homologs from distinct species, full-length sequence identity relative to PA14 ranged from 50.87% to 74.58%, whereas identity within the C-terminal DNA-contacting region spanning aa201–228 increased to 57.14%–92.86%. The phosphoacceptor residue D57 and the key DNA-contact residues S202, R205, and K207 were fully conserved, while L204 and L208 showed only conservative L/I substitutions (Fig. S12). Because our mutational analyses showed that representative promoters from all three regulatory classes rely on a common ParR DNA-contact surface, these data support the idea that output diversification is more likely to be driven by promoter cis architecture than by major changes in the ParR DNA-recognition mechanism. Adaptation to fluctuating host microenvironments or antibiotic pressure requires phenotypic flexibility. If new trans-acting regulators are costly to evolve, promoter architecture offers an accessible adaptive substrate(50). Insertions or deletions within a spacer could shift a target gene between defence- and virulence-associated outputs without altering the upstream signalling cascade. Such *cis*-regulatory plasticity may contribute to the repeated rewiring of resistance and virulence networks in adaptable pathogens such as *P. aeruginosa* (17).

These findings suggest that ParRS may act as a regulatory hub through which resistance- and virulence-associated programs can be co-modulated. More broadly, the ParR system provides an experimentally tractable example of how minimal *cis-regulatory* changes can reshape the output of a shared upstream signal. Disrupting the ParRS signalling hub, either by dampening kinase activation or by inhibiting RR-DNA engagement, could therefore affect both high-defence envelope states and virulence-associated programmes(51–53). For synthetic biology, the ParR system illustrates a compact design principle (53). If RR phosphorylation is treated as a shared input, promoter libraries can be designed in which minimal cis edits generate different activation thresholds and dynamic ranges (54). This architecture-first strategy provides a mechanistic foundation for building multi-output, state-dependent genetic circuits with a small protein footprint and shifts TCS engineering towards promoter-centric logic design.

## Conflict of interest

The authors declare that they have no conflict of interest.

## Acknowledgements

This work was supported by the National Natural Science Foundation of China (32370187 to Rui Bao) and the China Postdoctoral Science Foundation (GZC20241164 and 2025M771483 to Ninglin Zhao; GZC20241162 and 2025HXBH108 to Xingyu Mou). We thank Prof. Zhenling Wang (State Key Laboratory of Biotherapy, Sichuan University) for providing the *Pseudomonas aeruginosa* strains PAO1 and PA14, as well as the allelic-exchange vector pEX18Gm. We thank Prof. Yongxing He (School of Life Sciences, Lanzhou University) for kindly providing the β-galactosidase reporter plasmid pRG970Km, and Prof. Kelei Zhao (School of Pharmacy, Chengdu University) for the generous gift of the overexpression plasmid pL2a.

## Data availability

The RNA-seq data generated in this study have been deposited in the Genome Sequence Archive of the National Genomics Data Center, China National Center for Bioinformation, under accession number CRA041726. Other data supporting the findings of this study are available from the corresponding author upon reasonable request.

## Author Contributions

Rui Bao, Yingjie Song and Ziqi Zhu designed research; Ziqi Zhu, Ninglin Zhao and Dan Fan performed the experiments; Jianping Ying, Cheng Nong, Tonggen Liu, Chenxi Duan, Yaping Zheng, Shuang Zou, Xingyu Mou, Haichuan Ma and Huanxiang Liu provided technical support and insights; Ninglin Zhao and Ziqi Zhu analyzed the data; and Ninglin Zhao, Ziqi Zhu and Rui Bao wrote the manuscript.

---

**Fig. S1.**
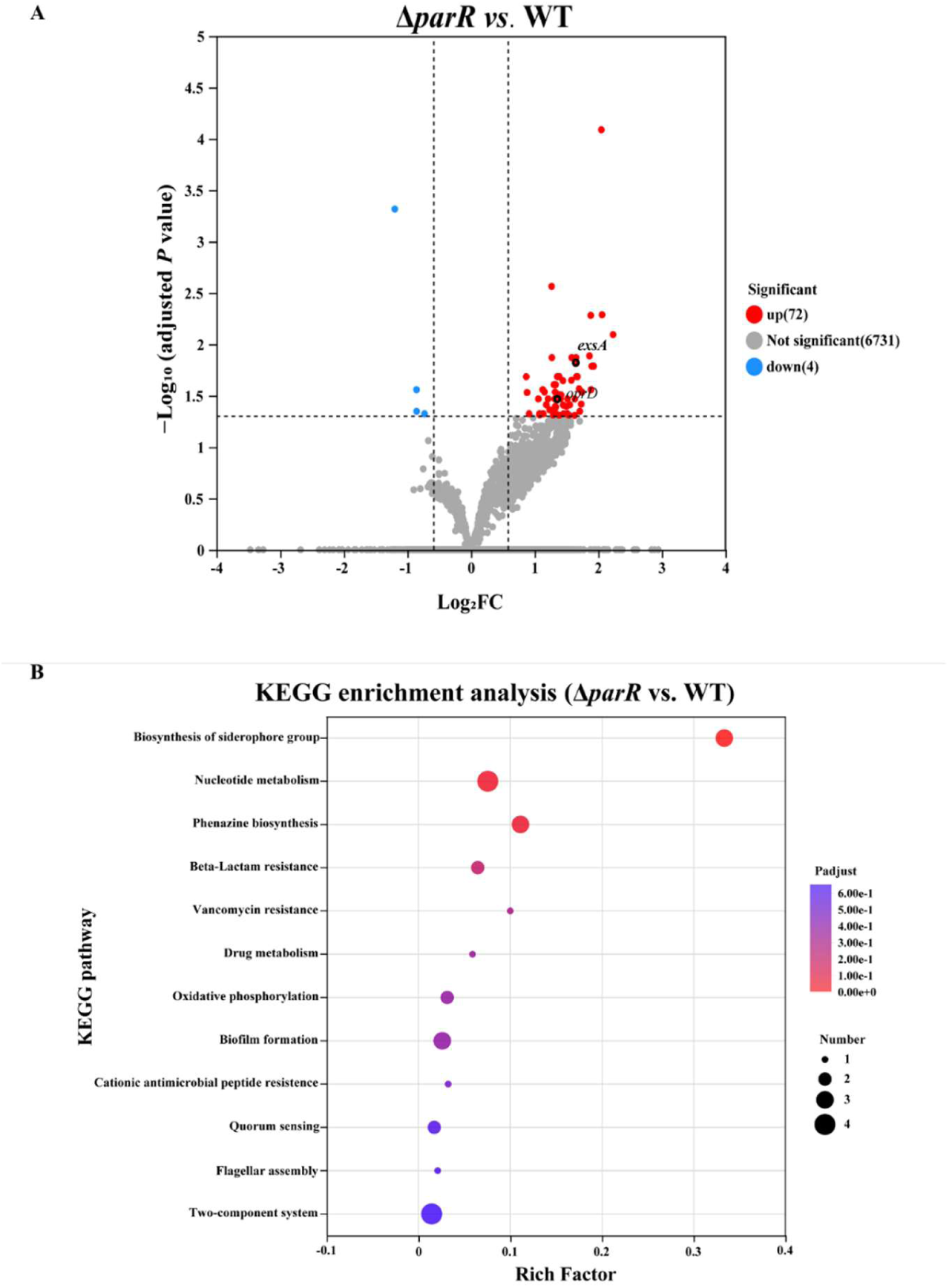
RNA-seq overview and KEGG enrichment analysis of differentially expressed genes in Δ*parR* versus WT. (A) Volcano plot of RNA-seq data comparing Δ*parR* and WT under matched growth conditions. Each dot represents one annotated gene. Differentially expressed genes were defined using the preset thresholds of fold change > 1.5 and FDR-adjusted *P* < 0.05. Upregulated genes are shown in red, downregulated genes in blue, and non-significant genes in gray. Vertical dashed lines indicate the fold-change cutoff, and the horizontal dashed line indicates the adjusted *P* value cutoff. Selected genes discussed in the main text are labeled. (B) KEGG enrichment analysis of differentially expressed genes identified in Δ*parR* versus WT. Bubble size indicates gene number, and bubble color indicates adjusted *P* value. Rich factor represents the ratio of differentially expressed genes to the total number of genes annotated in each pathway.

**Fig. S2.**
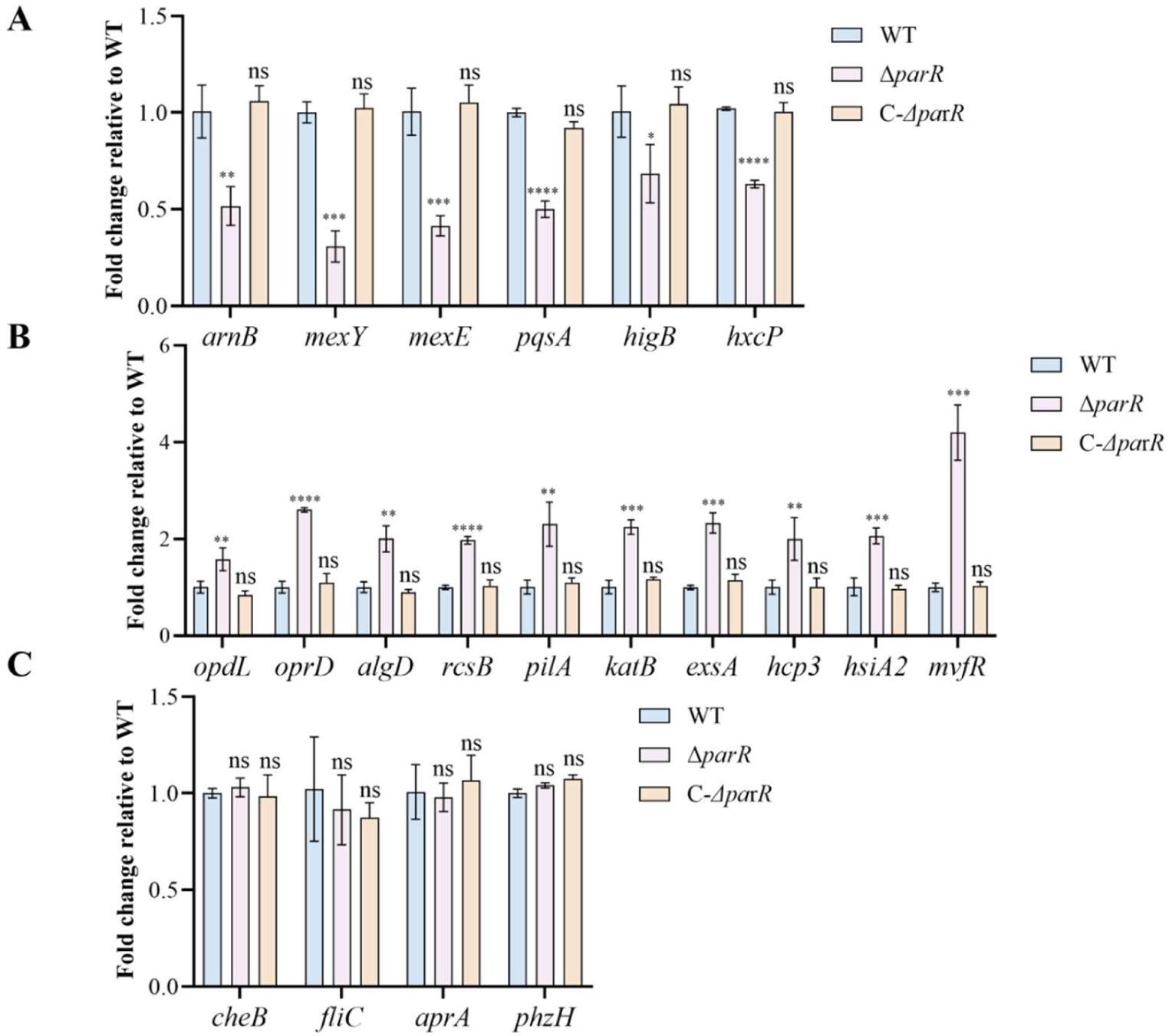
**RT-qPCR validation of the complete 20-genes candidate panel in WT, Δ*parR*, and C-Δ*parR*** Candidate genes were grouped according to their transcriptional behavior in the Δ*parR* mutant for display clarity. (A) Genes reduced in Δ*parR* relative to WT. (B) Genes increased in Δ*parR* relative to WT. (C) Genes showing no significant change under the tested conditions. Transcript levels were normalized to the internal reference gene and are shown relative to WT. The representative subset shown in Fig. 1C is included here as part of the complete 20 genes panel. Data are presented as mean ± SD from three independent experiments. Statistical significance is indicated in the figure. Symbols above the Δ*parR* and C-Δ*parR* bars indicate comparisons with WT within each gene. ns, not significant; \**P* < 0.05, \*\**P* < 0.01, \*\*\**P* < 0.001, \*\*\*\**P* < 0.0001.

**Fig. S3.**
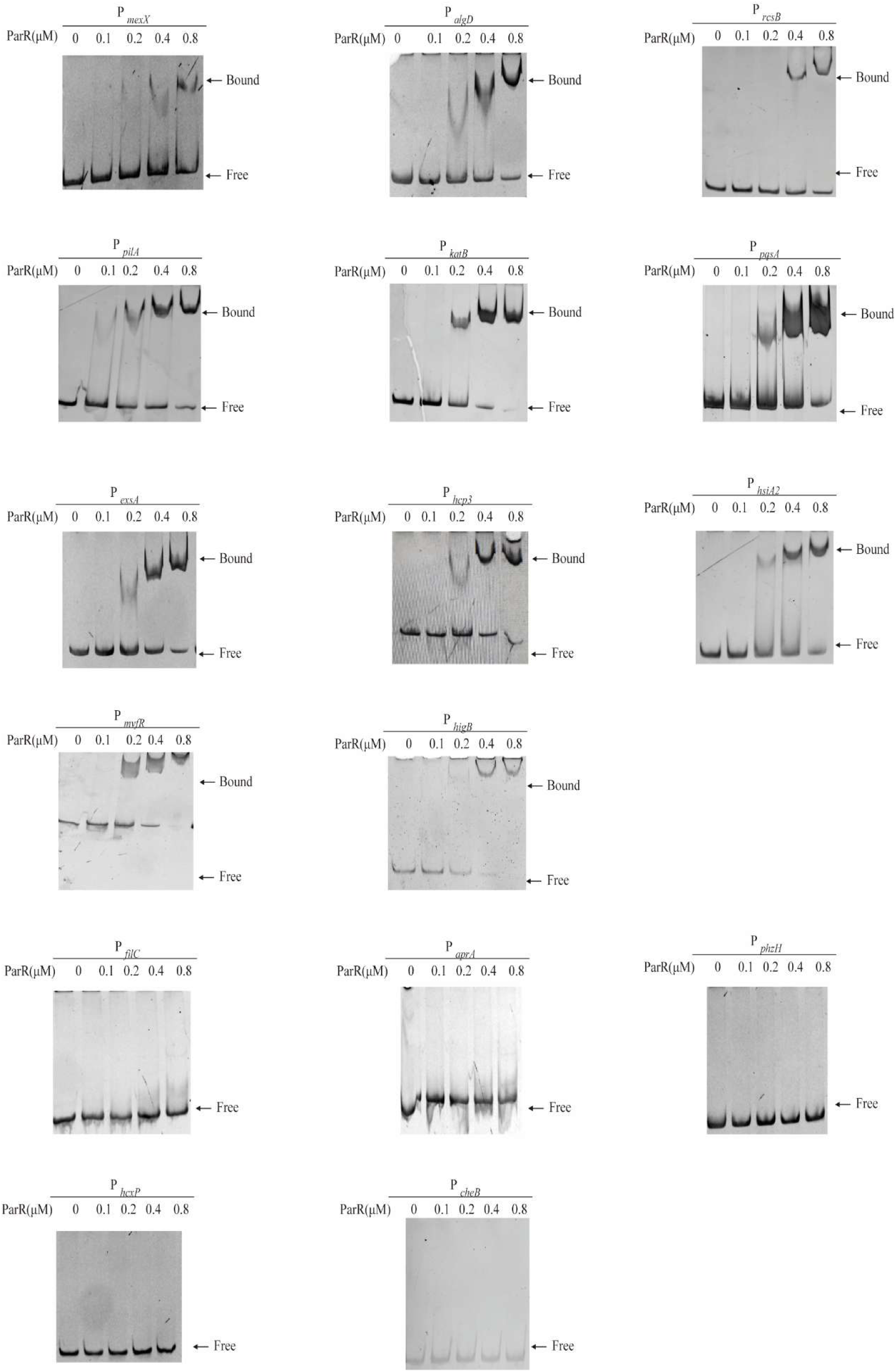
Full EMSA panel of candidate promoter probes, separated into shift-positive and no-shift groups. EMSAs were performed using approximately 200-bp promoter probes incubated with increasing concentrations of ParR (0, 0.1, 0.2, 0.4, and 0.8 μM). Representative promoter probes shown in Fig. 1D are omitted here; the remaining candidate promoter probes are displayed in this supplementary figure. Positions of free probe and ParR-DNA complexes are indicated. (A) Promoter probes showing detectable ParR-DNA shifting under the tested conditions. (B) Promoter probes showing no detectable ParR-DNA shift across the titration series. The semi-quantitative ranking of binding strength based on the minimum ParR: DNA ratio required for visible shifting is summarized in Table 3.

**Fig. S4.**
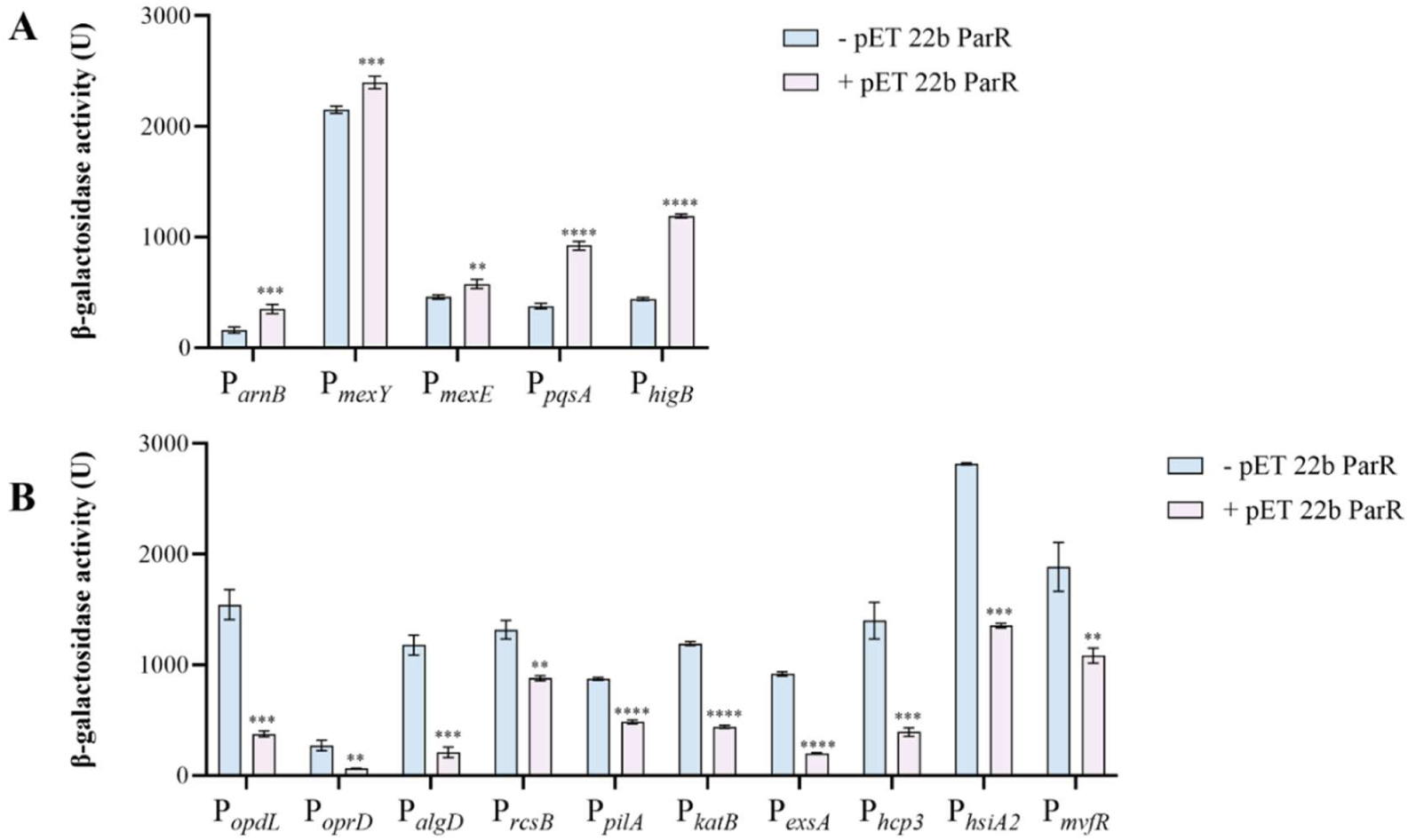
Promoter-*lacZ* reporter assays for EMSA-positive candidate promoters reveal ParR-dependent activation and repression. Reporter activity was measured in cells carrying the empty vector or pET22b-ParR and is shown as β-galactosidase activity. Promoters were grouped according to the direction of ParR-dependent output for display clarity. (A) Promoters activated by ParR expression. (B) Promoters repressed by ParR expression. Data are presented as mean ± SD from three independent experiments. Statistical significance is indicated in the figure and reflects comparisons between the under the empty-vector and ParR-construct conditions within each promoter. \**P* < 0.05, \*\**P* < 0.01, \*\*\**P* < 0.001, \*\*\*\**P* < 0.0001.

**Fig. S5.**
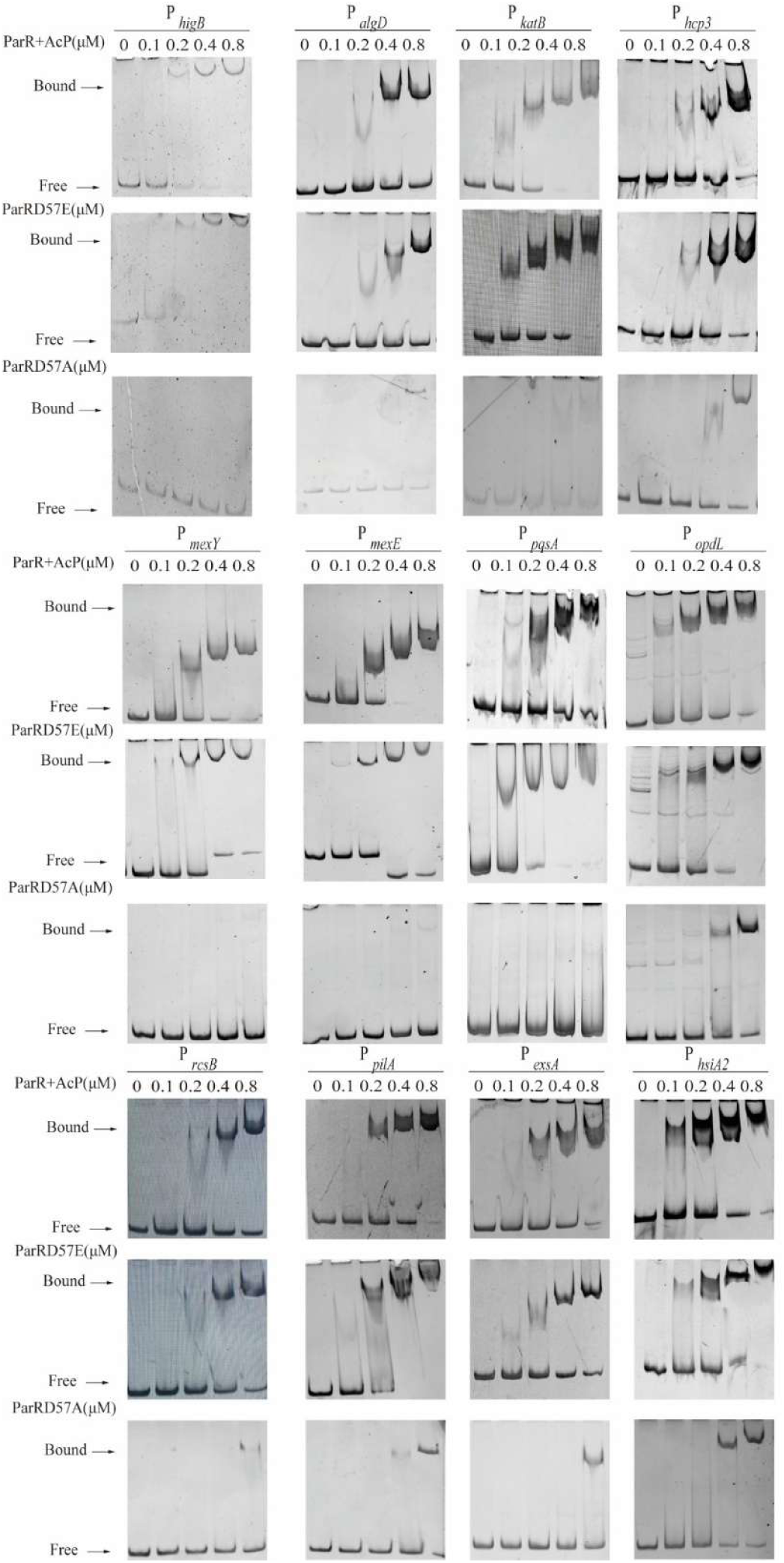
Additional EMSAs for the remaining direct target promoters not shown in. Fig. 2B This figure shows EMSA results for the remaining 12 ParR target promoters not displayed in Fig. 2B, including P*_higB_,* P*_algD_,* P*_katB_,* P*_hcp3_,* P*_mexY_,* P*_mexE_,* P*_pqsA_,* P*_opdL_,* P*_rcsB_,* P*_pilA_,* P*_exsA_,* and P*_hsiA2_*. ParR+AcP, ParRD57E and ParRD57A were incubated with the indicated promoter fragments at increasing protein concentrations (0, 0.1, 0.2, 0.4, and 0.8 μM). Bound and free DNA species are indicated. Together with the representative gels shown in Fig. 2B, these data complete the promoter resolved EMSA dataset underlying the binding-threshold analysis described in the main text.

**Fig. S6.**
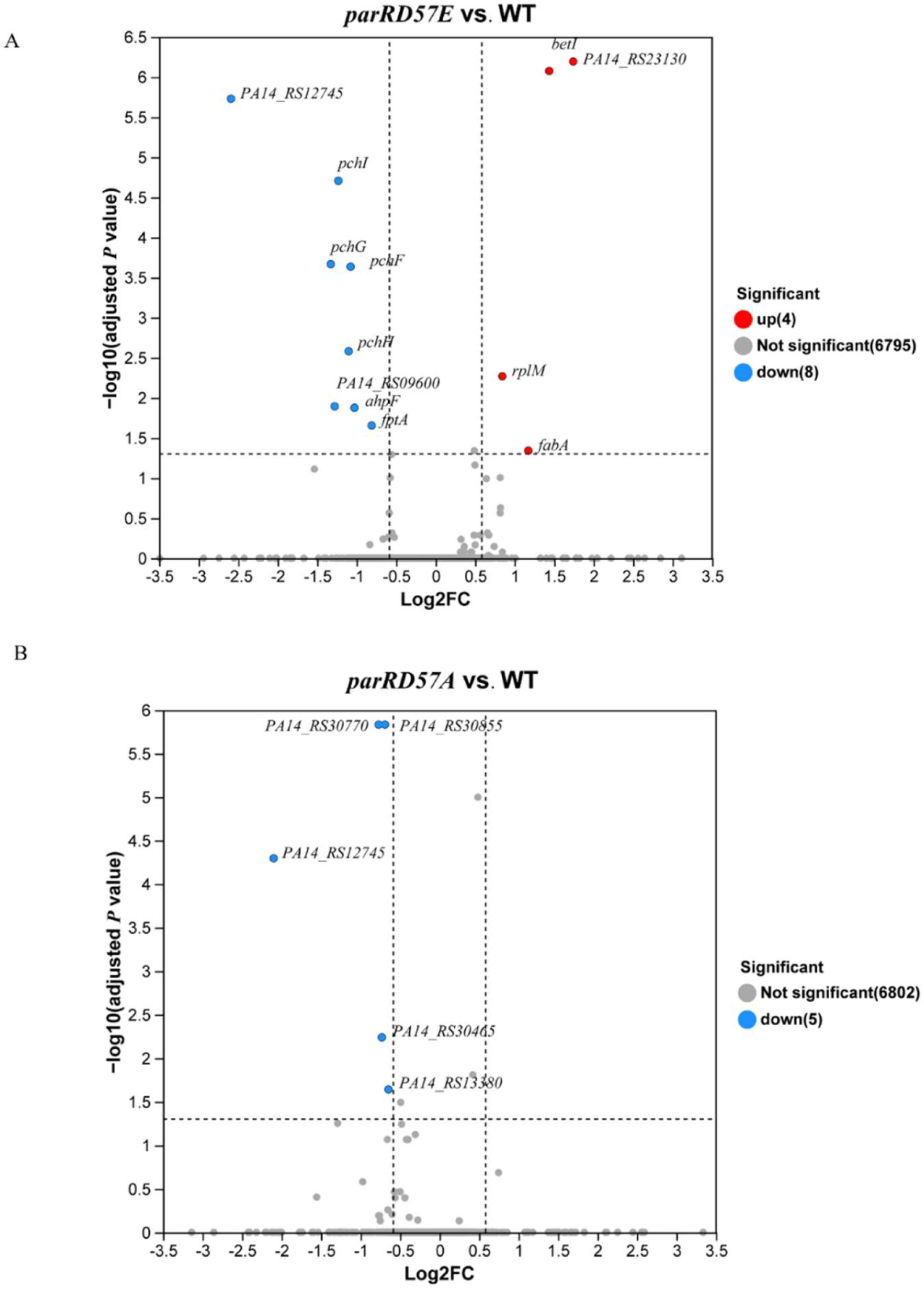
Transcriptomic comparison of *parRD57E* and *parRD57A* against WT by volcano plot analysis. (A) Differentially expressed genes identified in the *parRD57E* versus WT comparison. Upregulated genes are shown in red, downregulated genes in blue, and non-significant genes in gray; no significantly upregulated genes were detected under the applied cutoff. (B) Differentially expressed genes identified in *parRD57A* versus WT comparison. Downregulated genes are shown in blue and non-significant genes in gray; no significantly upregulated genes were detected under the applied cutoff. In both panels, the x-axis indicates log_2_ fold change (Log_2_FC), and the y-axis indicates −log_10_(adjusted *P* value). Vertical dashed lines indicate the fold-change cutoff, and the horizontal dashed line indicates the adjusted *P* value cutoff. Labeled points denote representative significant genes.

**Fig. S7.**
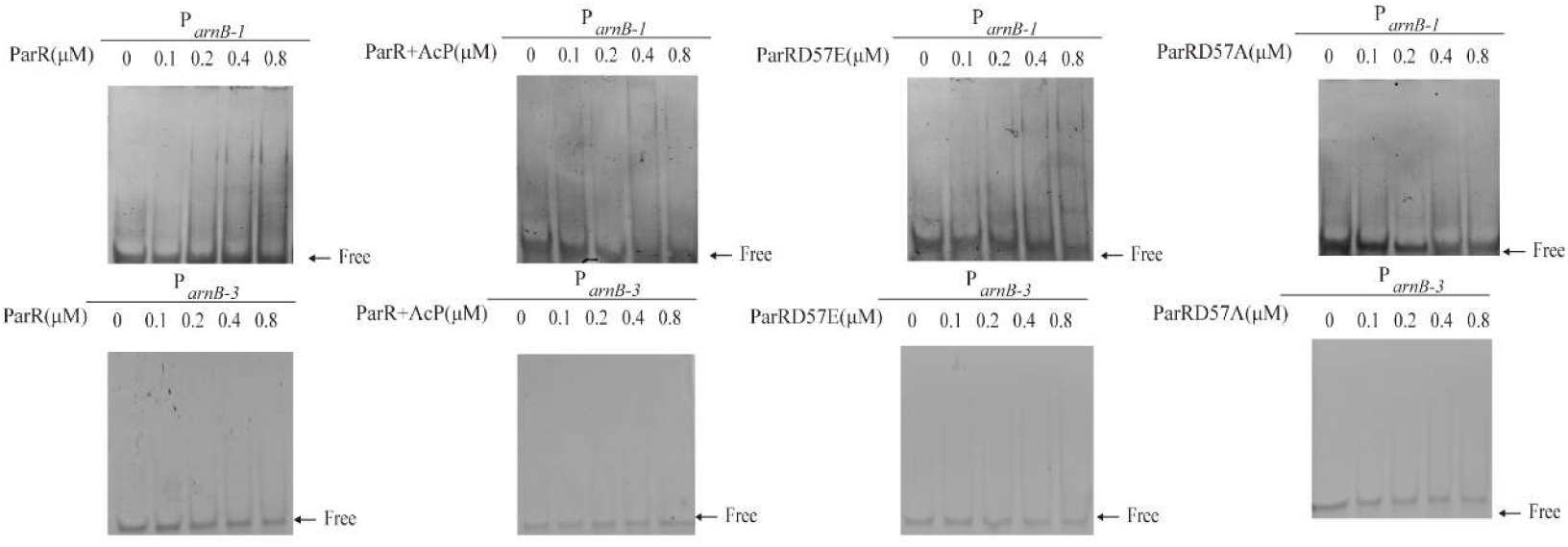
EMSA screening of additional MEME predicted candidates in the *arnB* intergenic region reveals probe-selective ParR binding. MEME identified multiple palindromic candidate elements within the *arnB* intergenic region. EMSAs were performed using probes corresponding to P*_arnB_*_-1_ and P*_arnB_*_-3_ under the same assay conditions used for the validated P*_arnB_*_-2_ probe in Fig. 4C. For each probe, ParR, ParR+AcP, ParRD57E, and ParRD57A were titrated over the indicated concentration range (0, 0.1, 0.2, 0.4, and 0.8 μM). Under these matched conditions, P*_arnB_*_-1_ and P*_arnB_*_-3_ showed no obvious shift or only very weak complex formation. MEME predicted candidates differ in ParR binding capacity and should be interpreted together with the promoter mapping and functional validation data for P*_arnB_*_-2_ in Fig. 4D.

**Fig. S8.**
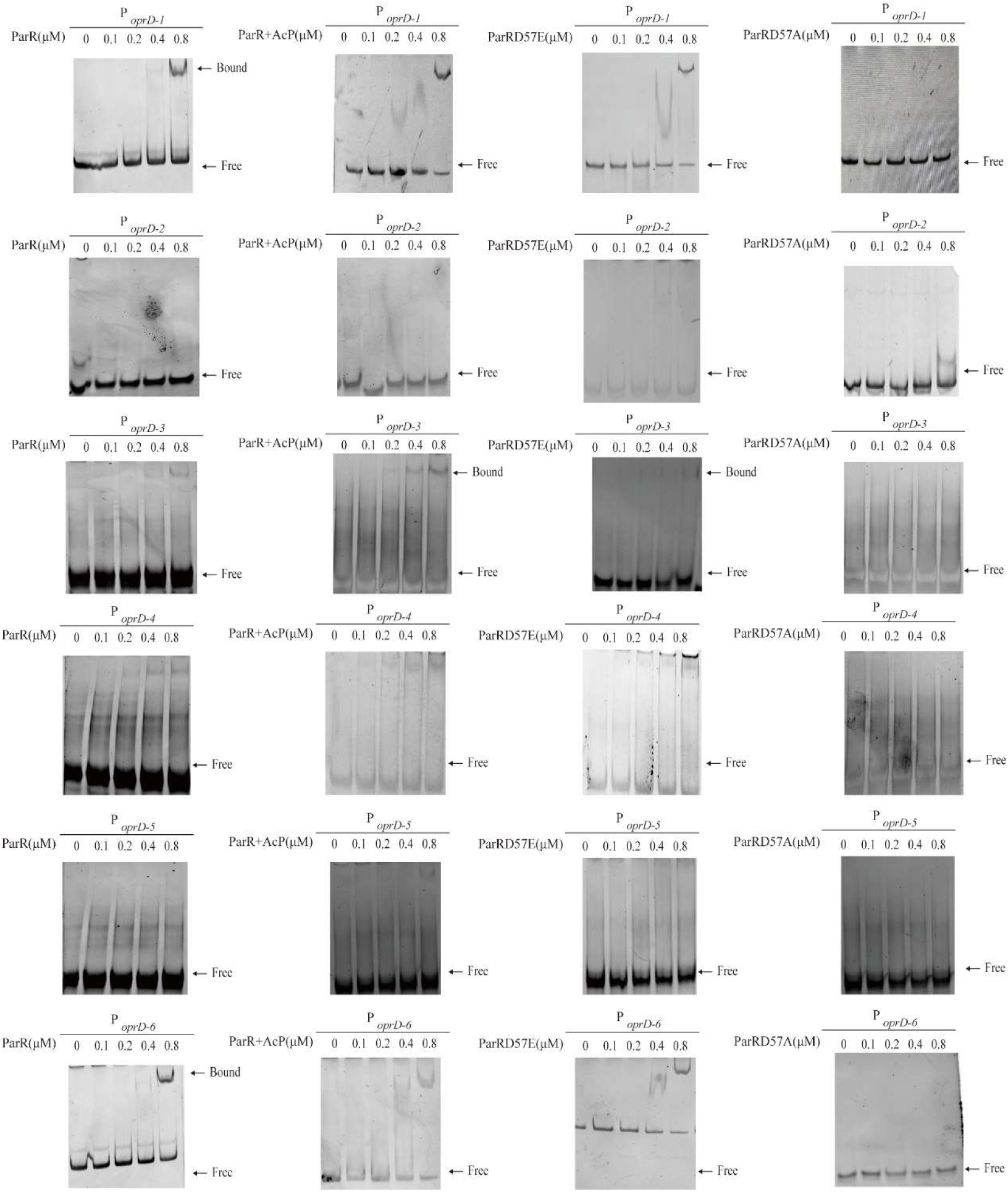
EMSA screening of additional MEME predicted candidates in the *oprD* intergenic region reveals probe-selective ParR binding. MEME identified multiple candidate elements within the *oprD* intergenic region. To examine the binding capacities of additional candidate probes, EMSAs were performed using P*_oprD-1_*, P*_oprD-2_*, P*_oprD-3_*, P*_oprD-4_*, P*_oprD-5_*, and P*_oprD-6_* under the same assay framework used for the validated P*_oprD-7_* probe. For each probe, ParR, ParR+AcP, ParRD57E, and ParRD57A were titrated over the indicated concentration range (0, 0.1, 0.2, 0.4, and 0.8 μM). Under these conditions, the alternative probes showed variable binding behavior, with some probes displaying detectable or condition-dependent complex formation and others showing little or no obvious shift. Free probe is indicated, and bound complex is marked where visible. These data indicate that additional MEME predicted candidates in the *oprD* region differ in ParR binding capacity and should be interpreted together with the mapping and functional validation data for P*_oprD-7_* in Fig. 4H.

**Fig. S9.**
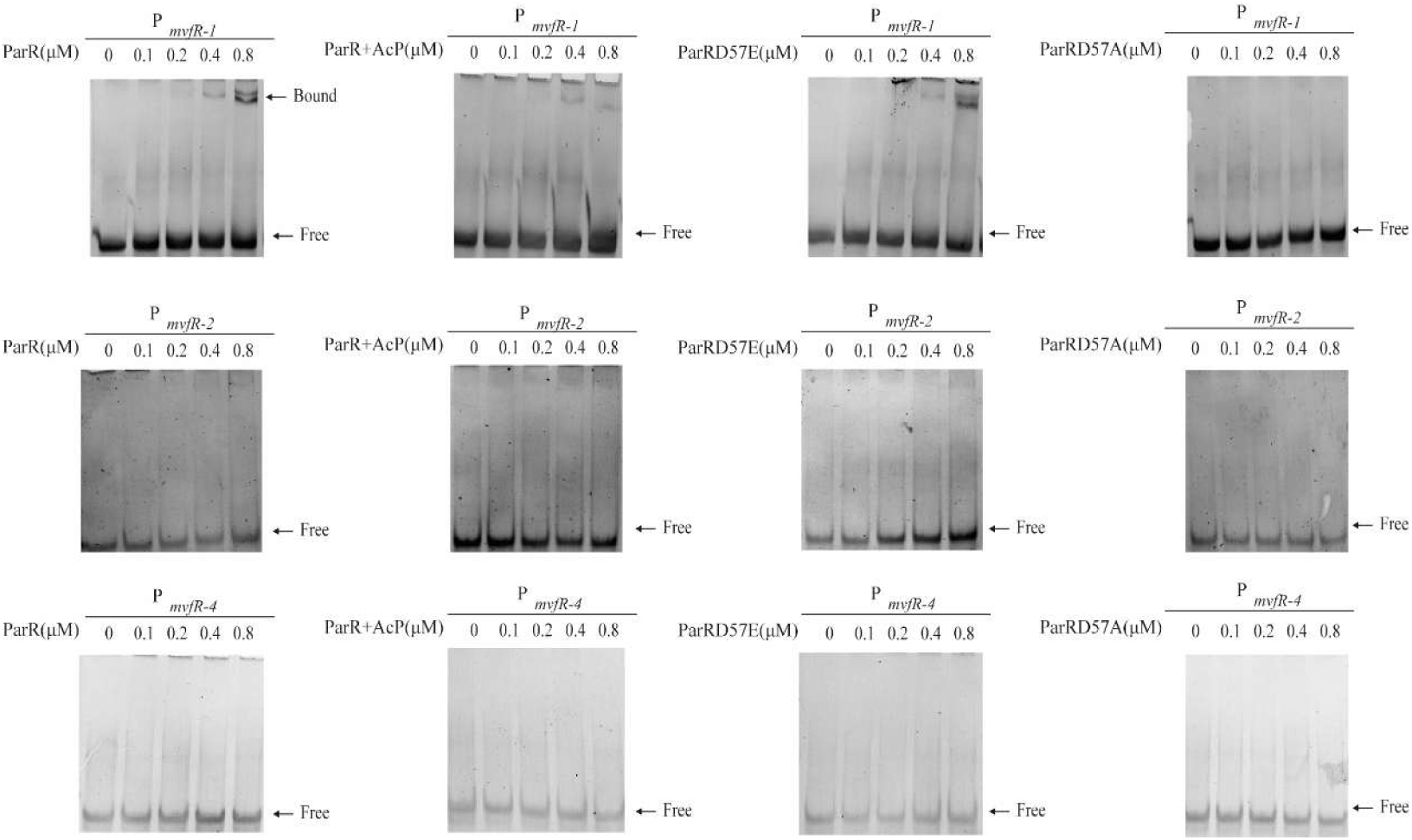
EMSA screening of additional MEME predicted candidates in the *mvfR* intergenic region reveals probe-selective ParR binding. MEME identified multiple candidate elements within the *mvfR* intergenic region. To examine the binding capacities of additional candidate probes, EMSAs were performed using P*_mvfR-1_*, P*_mvfR-2_*, and P*_mvfR-4_* under the same assay framework used for the validated P*_mvfR-3_* probe. For each probe, ParR, ParR+AcP, ParRD57E, and ParRD57A were titrated over the indicated concentration range (0, 0.1, 0.2, 0.4, and 0.8 μM). Under these conditions, P*_mvfR-1_* showed detectable complex formation, whereas P*_mvfR-2_* and P*_mvfR-4_* exhibited little or no obvious shift. Free probe is indicated; bound complex is marked where visible. These results indicate that alternative MEME predicted candidates differ in ParR binding capacity and should be interpreted together with the promoter mapping and functional validation data for P*_mvfR-3_* in Fig. 4L.

**Fig. S10.**
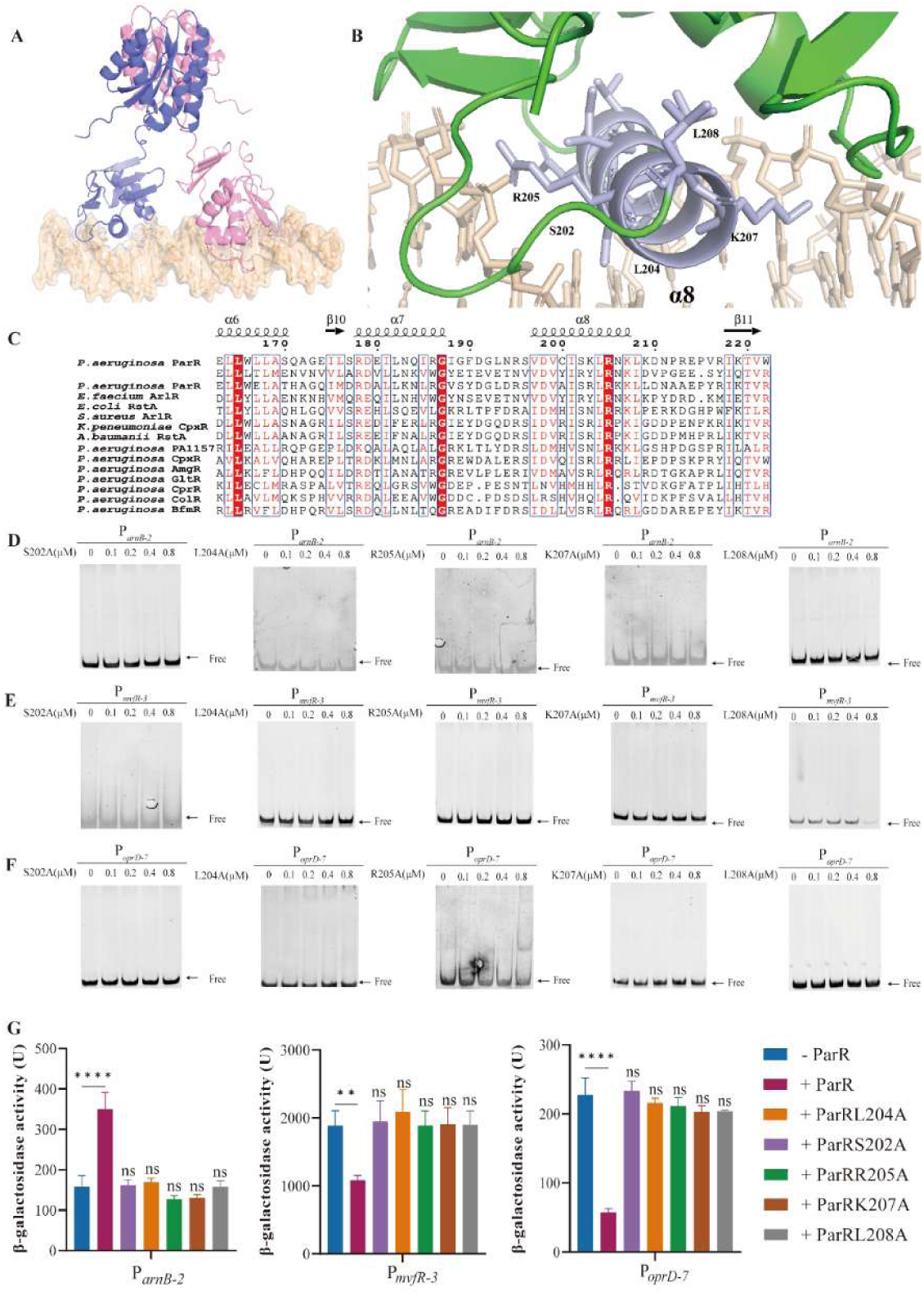
Structure-guided analysis of predicted ParR DNA contact residues supports a common interface for representative target promoters. (A) Overall view of the AlphaFold3 predicted ParR-DNA (P*_arnB-2_*) complex. (B) Enlarged view of the predicted ParR-DNA interface, showing residues selected for alanine-substitution analysis on the putative DNA-binding surface. (C) Multiple-sequence alignment of ParR with representative OmpR/PhoB-family response regulators, illustrating conservation of the corresponding interface region. (D-F) EMSAs of alanine substituted ParR variants using the representative probes P*_arnB-2_* (D), P*_mvfR-3_* (E), and P*_oprD-7_* (F). The indicated mutant proteins were titrated from 0 to 0.8 μM under the same assay framework. Free probe is indicated. (G) β-Galactosidase reporter assay of the corresponding P*_arnB-2_*, P*_mvfR-3_*, and P*_oprD-7_* reporters in the indicated expression backgrounds. Bars represent mean ± SD; *n* = 3 biologically independent experiments. Statistical significance is indicated in the figure. One-way ANOVA with Tukey’s multiple-comparisons test was used for panel G. ns, not significant; \**P* < 0.05, \*\**P* < 0.01, \*\*\**P* < 0.001, \*\*\*\**P* < 0.0001. These data support the interpretation that residues on the predicted ParR DNA contact surface contribute broadly to representative target binding and reporter regulation.

**Fig. S11.**
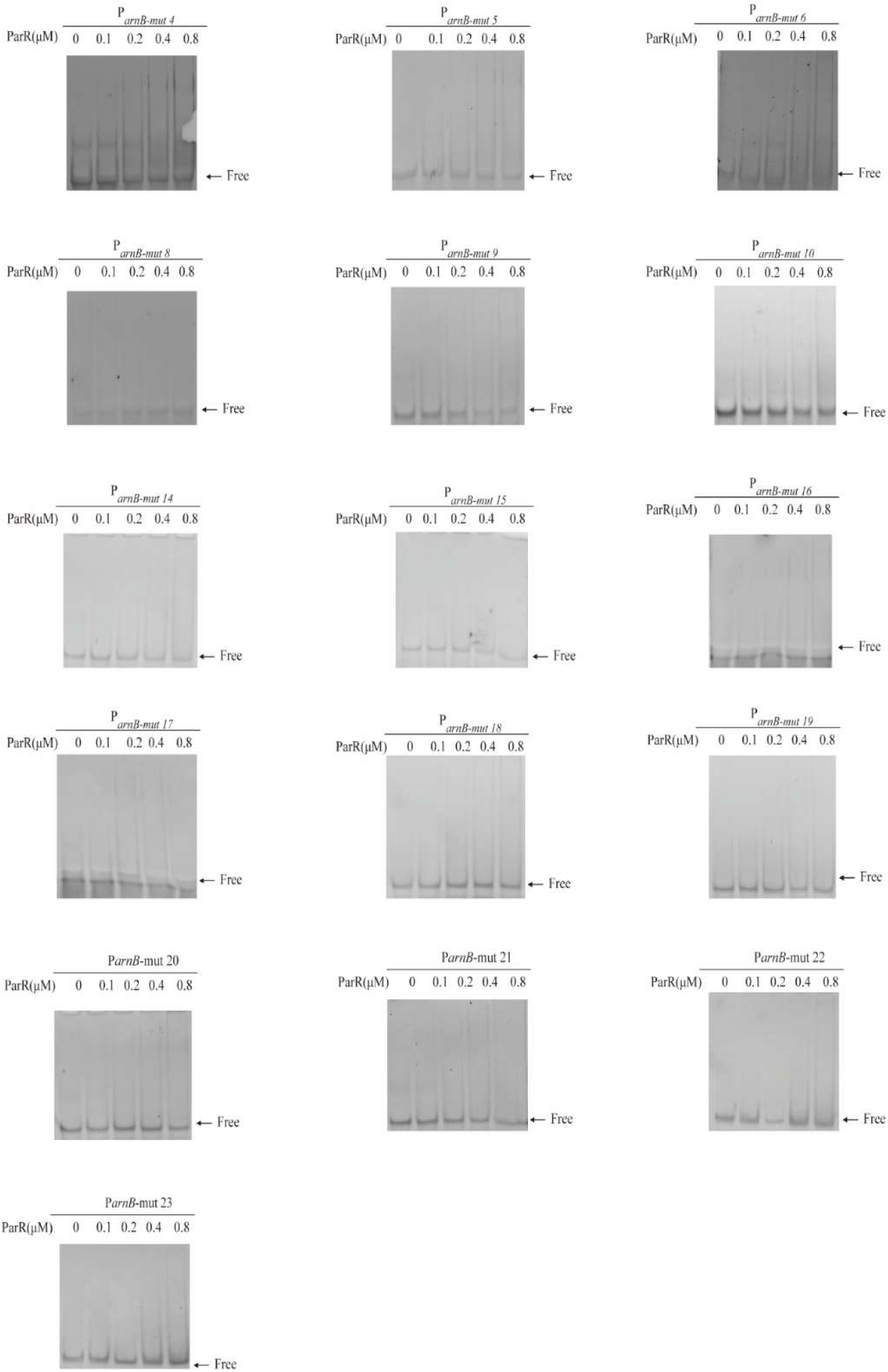
ParR primary EMSA screen of the P*_arnB-2_* mutational series. EMSA screening of ParR binding to the P*_arnB-2_* mutant panel (mut1-mut23), grouped by mutation class (spacer deletions, half-sites substitutions, centre polyN substitutions, and spacer insertions). Each probe was first screened using ParR across the indicated protein: DNA titrations. Only binding-permissive variants (mut1, mut2, mut13, mut3, mut7, mut11, and mut12) were subsequently assayed with additional ParR activity states. Variants with no detectable ParR binding in the primary screen were not further tested with other ParR states.

**Fig. S12.**
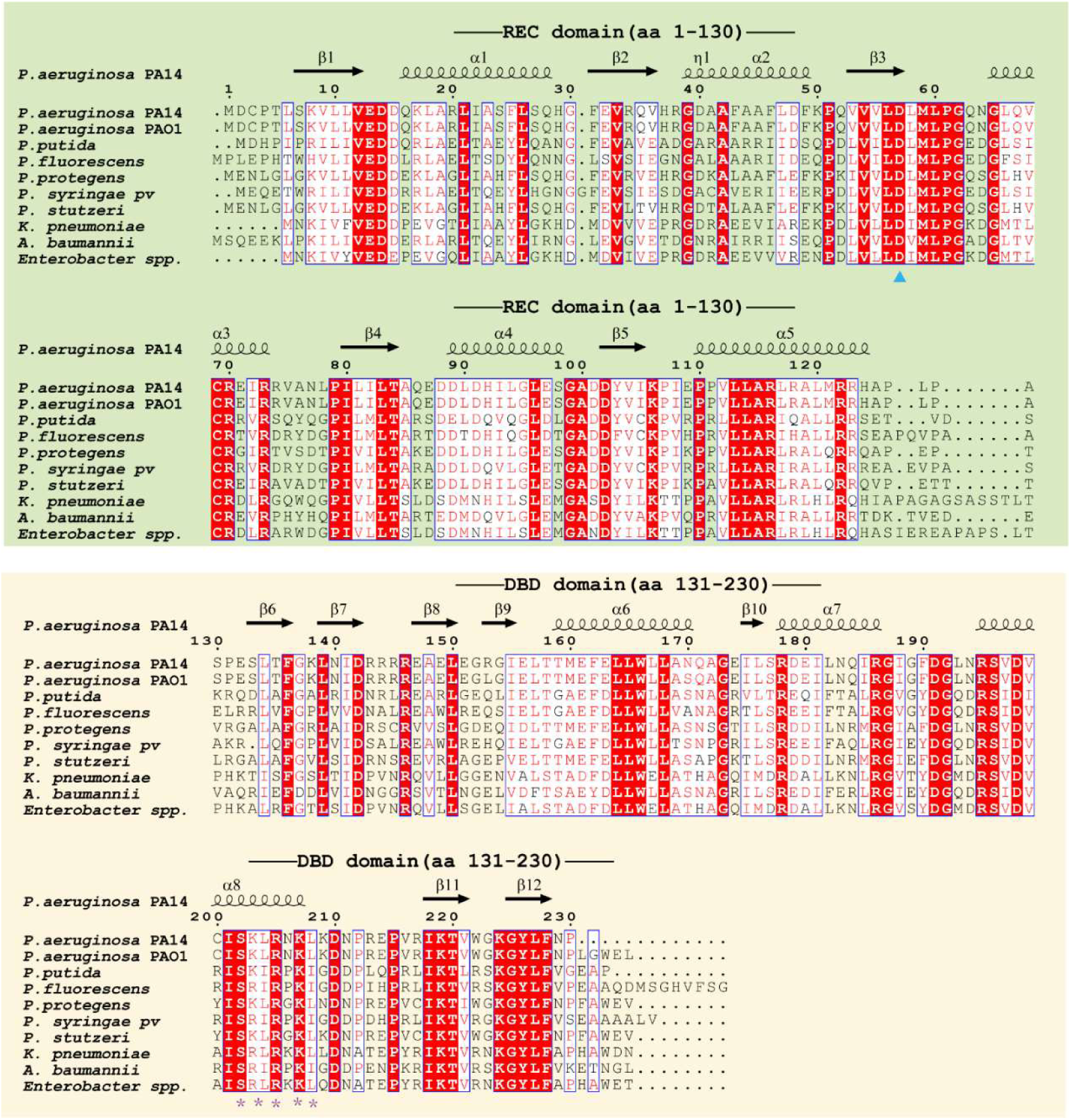
Conservation of the ParR DNA-binding interface across representative *Pseudomonas* homologs and selected non-Pseudomonas homologs Multiple-sequence alignment of ParR homologs from representative *Pseudomonas* species and selected non-*Pseudomonas* Gram-negative bacteria. The N-terminal receiver domain (REC) and the C-terminal DNA-binding domain (DBD) are indicated above the alignment. Compared with the full-length protein sequence, the DBD region shows a higher degree of conservation across the aligned homologs. Black triangle indicates the conserved phosphoacceptor Asp57. Asterisks indicate conserved DNA-contact residues S202, L204, R205, K207, and L208. Residue numbering is based on *P. aeruginosa* PA14 ParR.

**Table S1.** Key resources table.

| Reagent or Resource | Source | Identifier |
| --- | --- | --- |
| **Bacterial cells and plasmids** |  |  |
| *E. coli* BL21(DE3) | Beijing Genesand Biotech Co.,Ltd | Cat# SEC19 |
| *E. coli* DH5α | Beijing Genesand Biotech Co.,Ltd | Cat#SCC01 |
| *E. coli* S17-1 | This study | N/A |
| *P. aeruginosa* PA14 | Prof. Zhenling Wang LAB | N/A |
| *P. aeruginosa* PA14 $\Delta parR$ | This study | |
| *P. aeruginosa* PA14 C-$\Delta parR$ | This study | |
| *P. aeruginosa* PA14 *parRD57E* | This study |  |
| *P. aeruginosa* PA14 *parRD57A* | This study |  |
| pET-22b | This study |  |
| pRG970 | Prof. Yongxing He LAB |  |
| pEX18-Gm | Prof. Zhenling Wang LAB |  |
| pL2a | Prof. Kelei Zhao LAB |  |
| **Chemicals** |  |  |
| Ni-NTA agarose | ThermoFisher | Cat# R90110 |

**Table S2.**
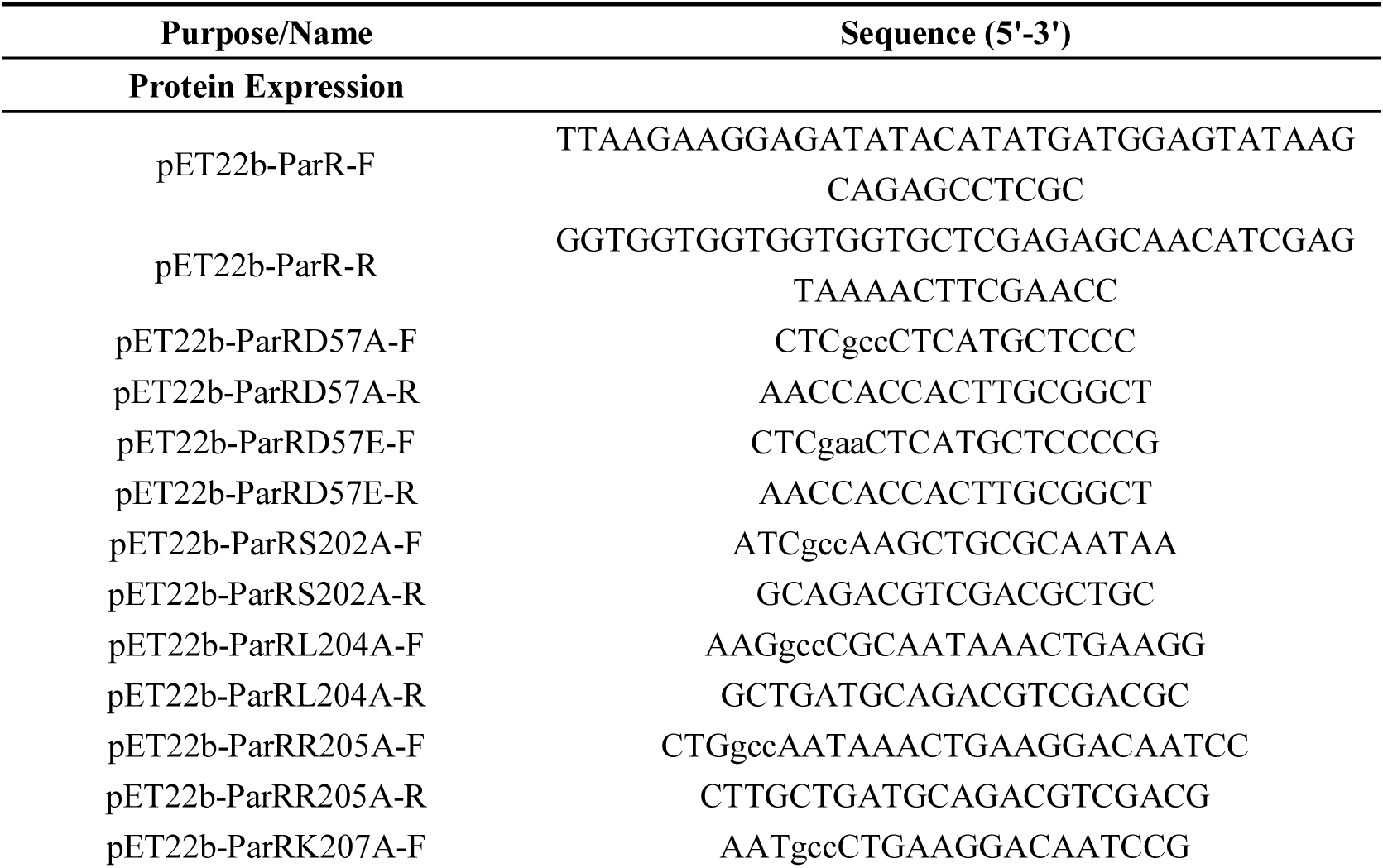
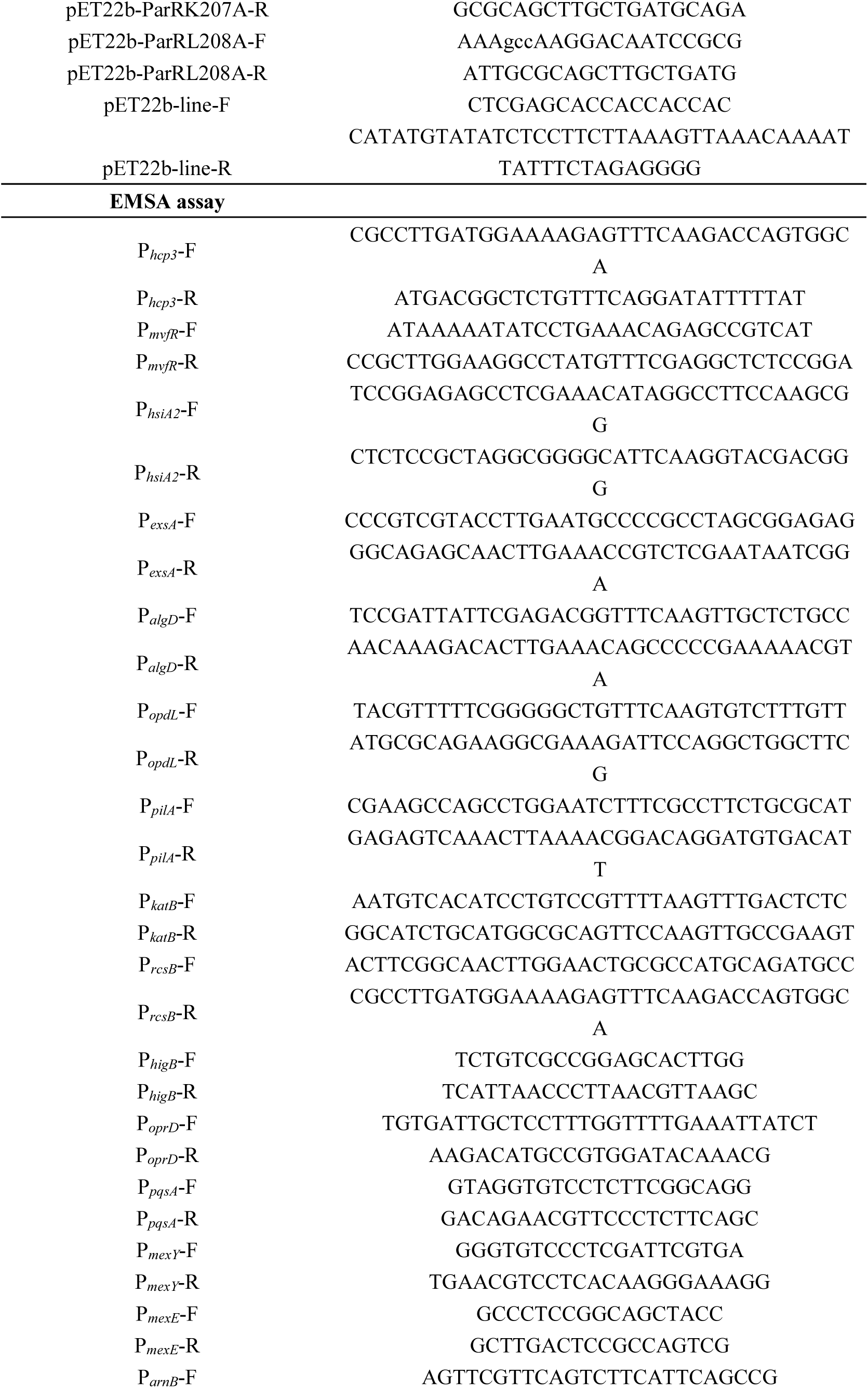
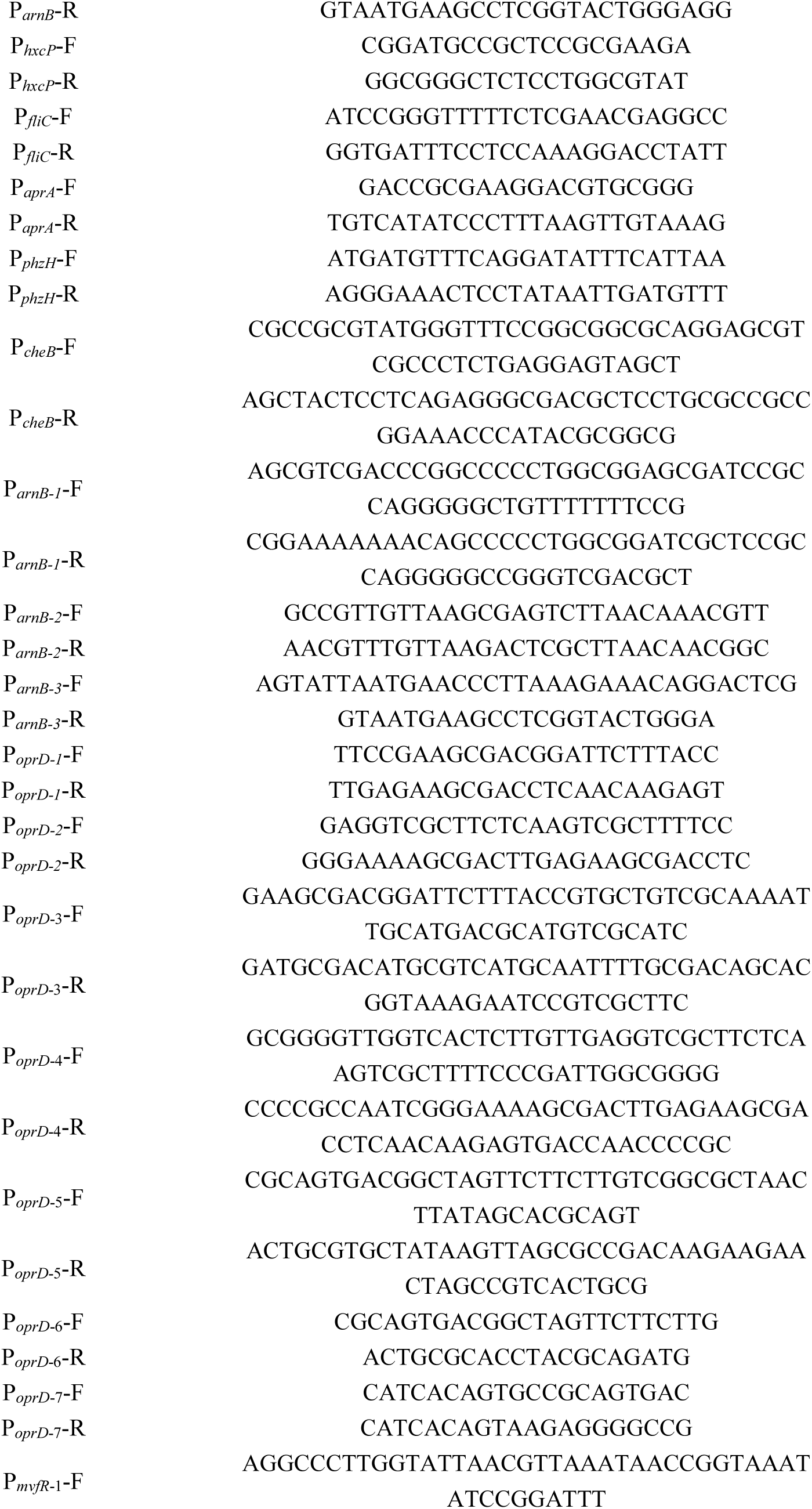
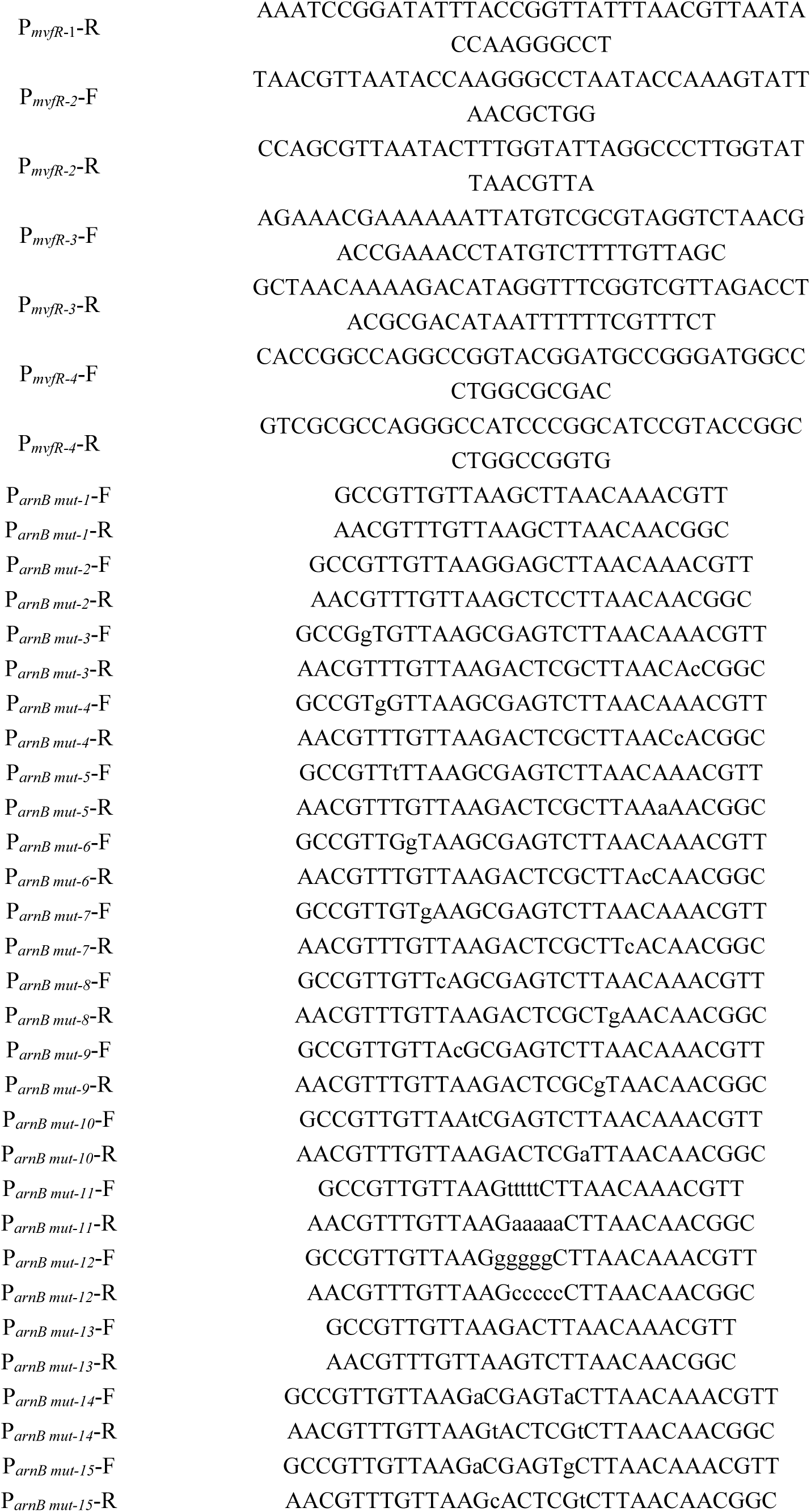
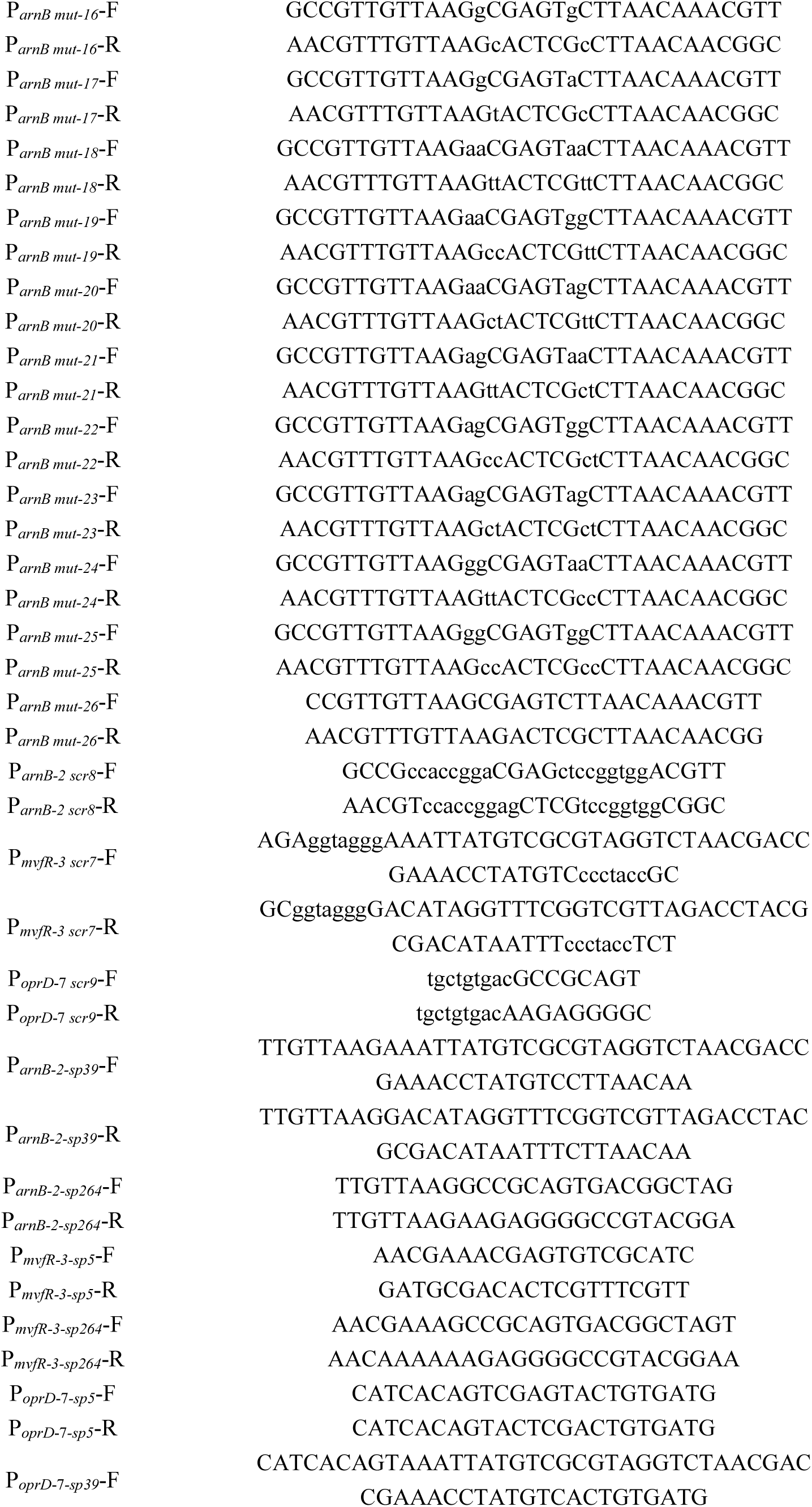
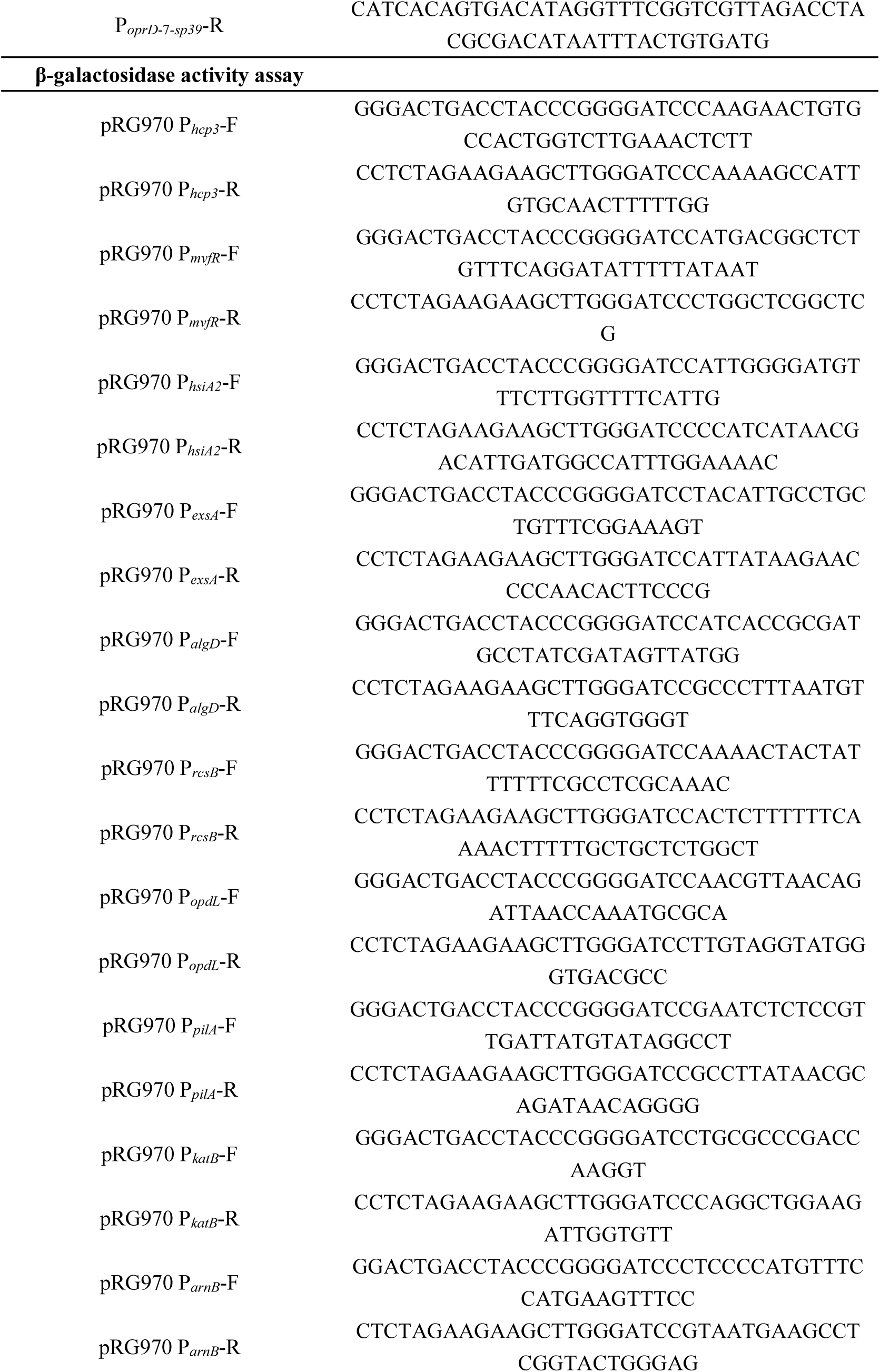
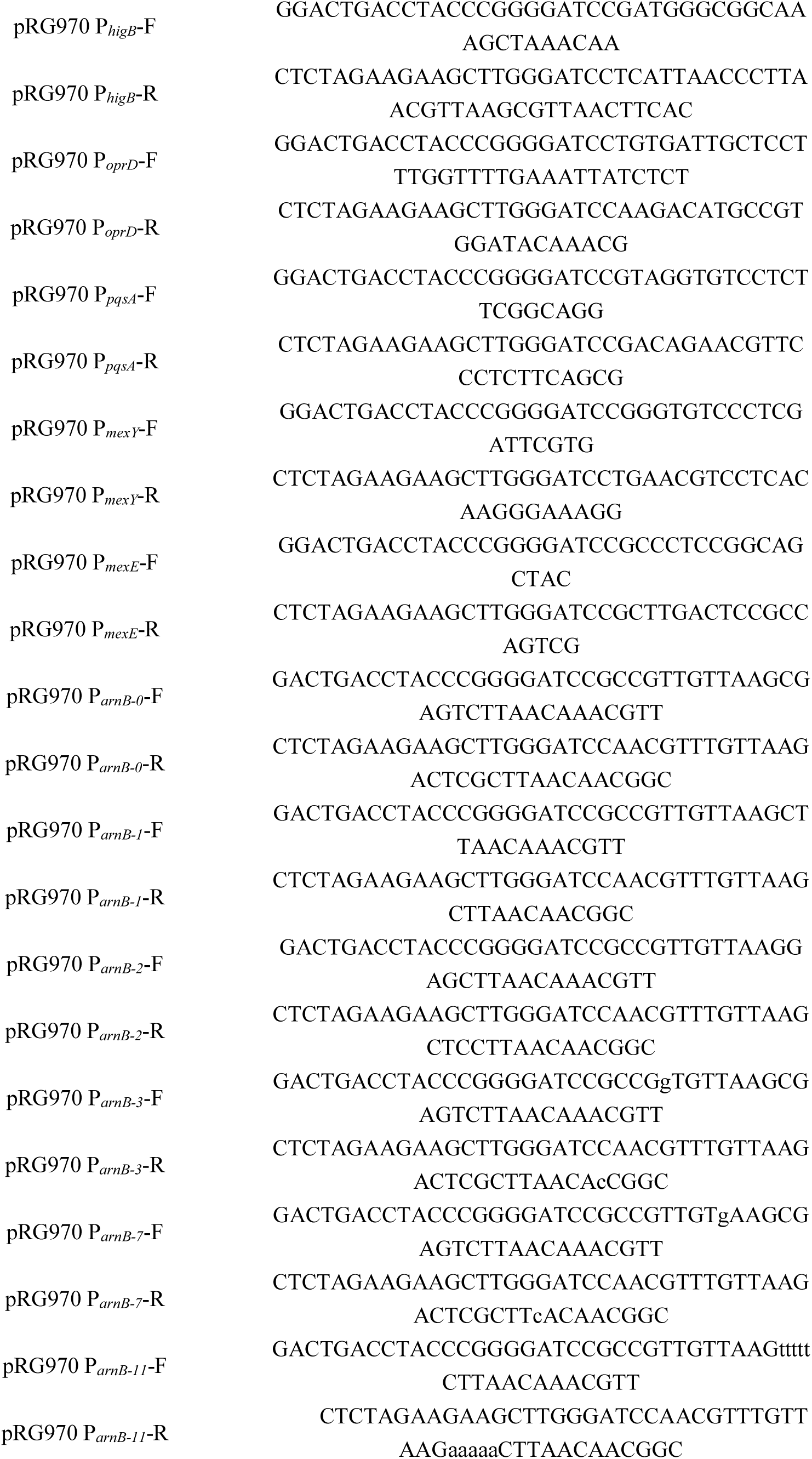
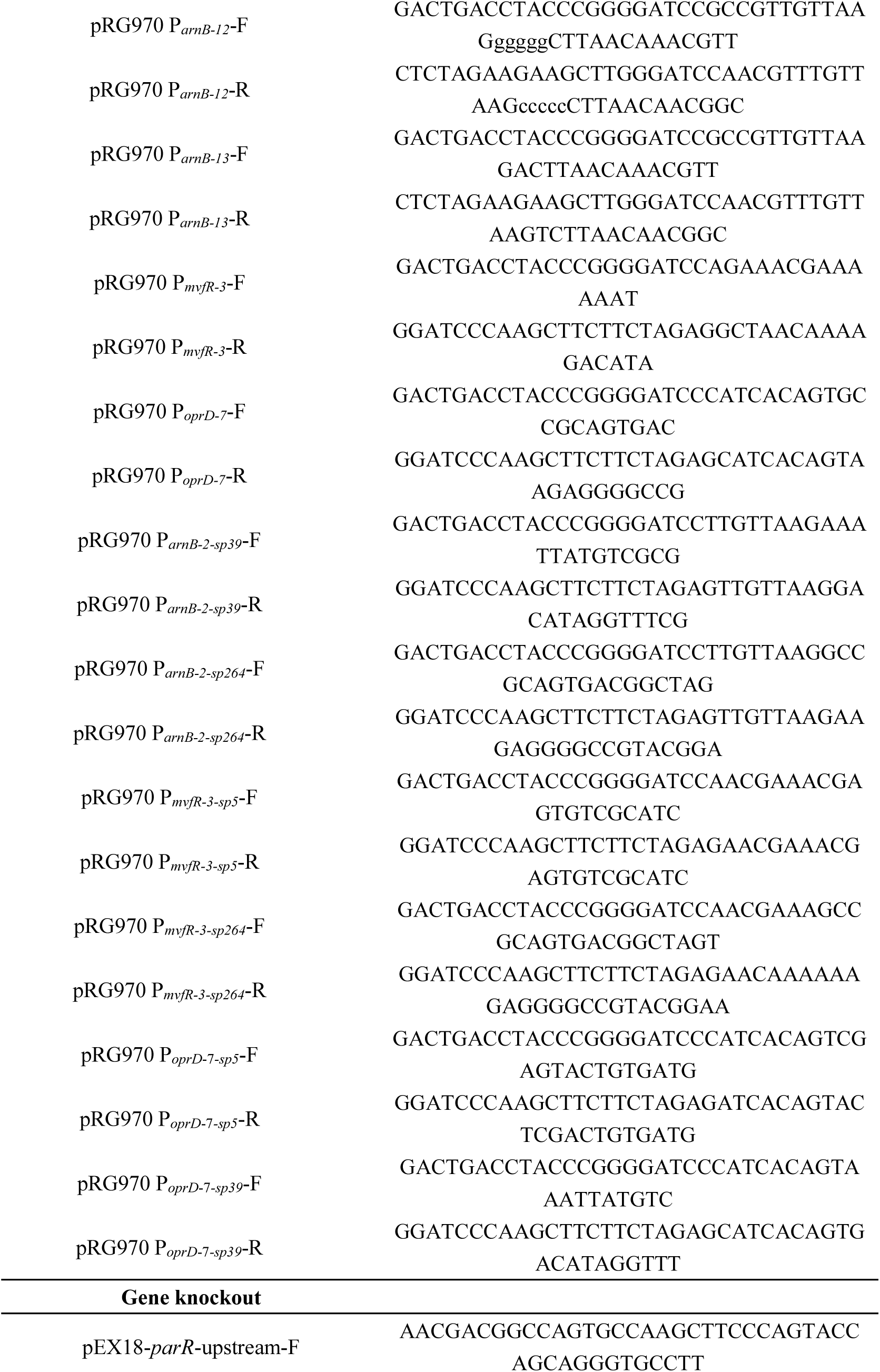
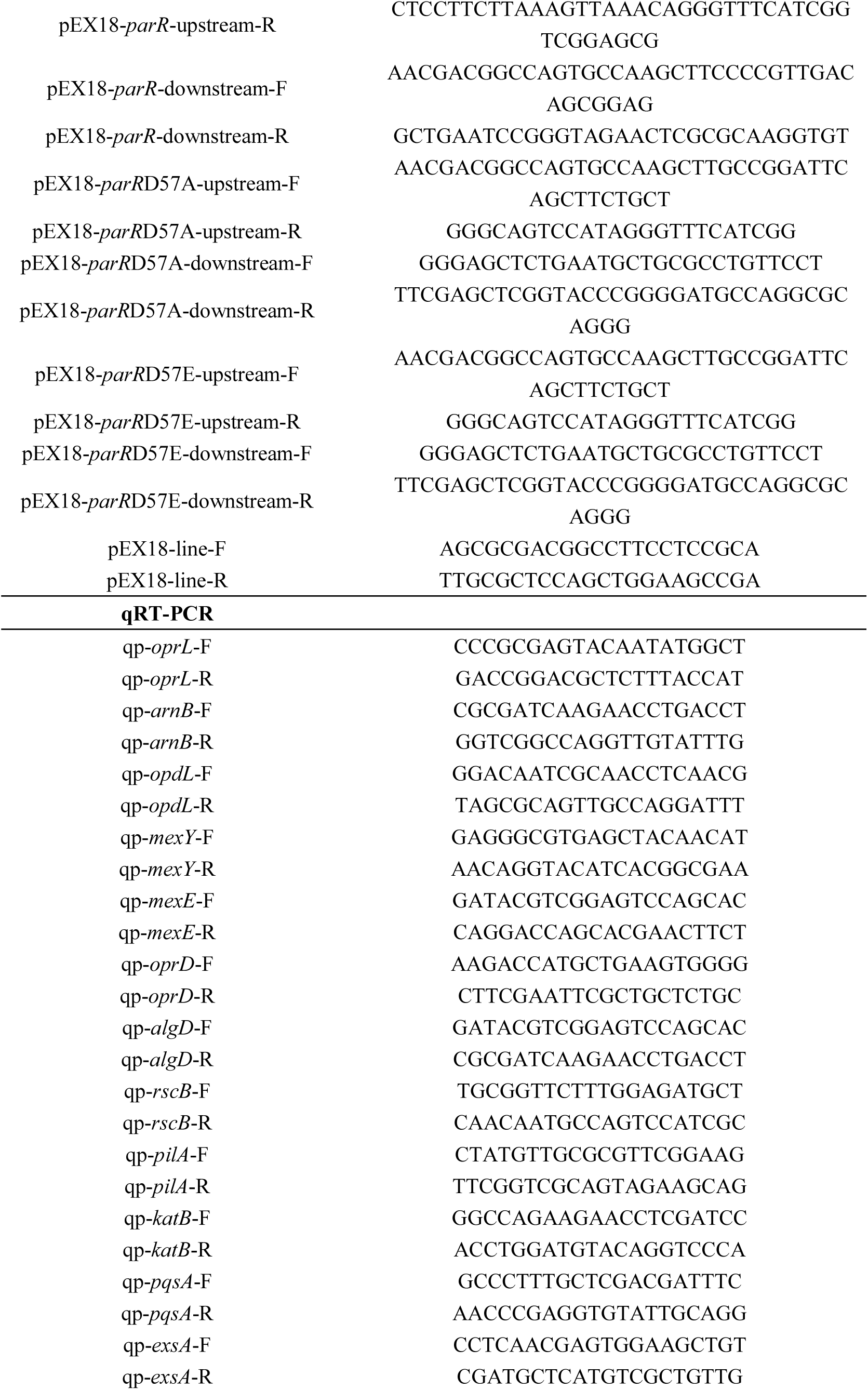
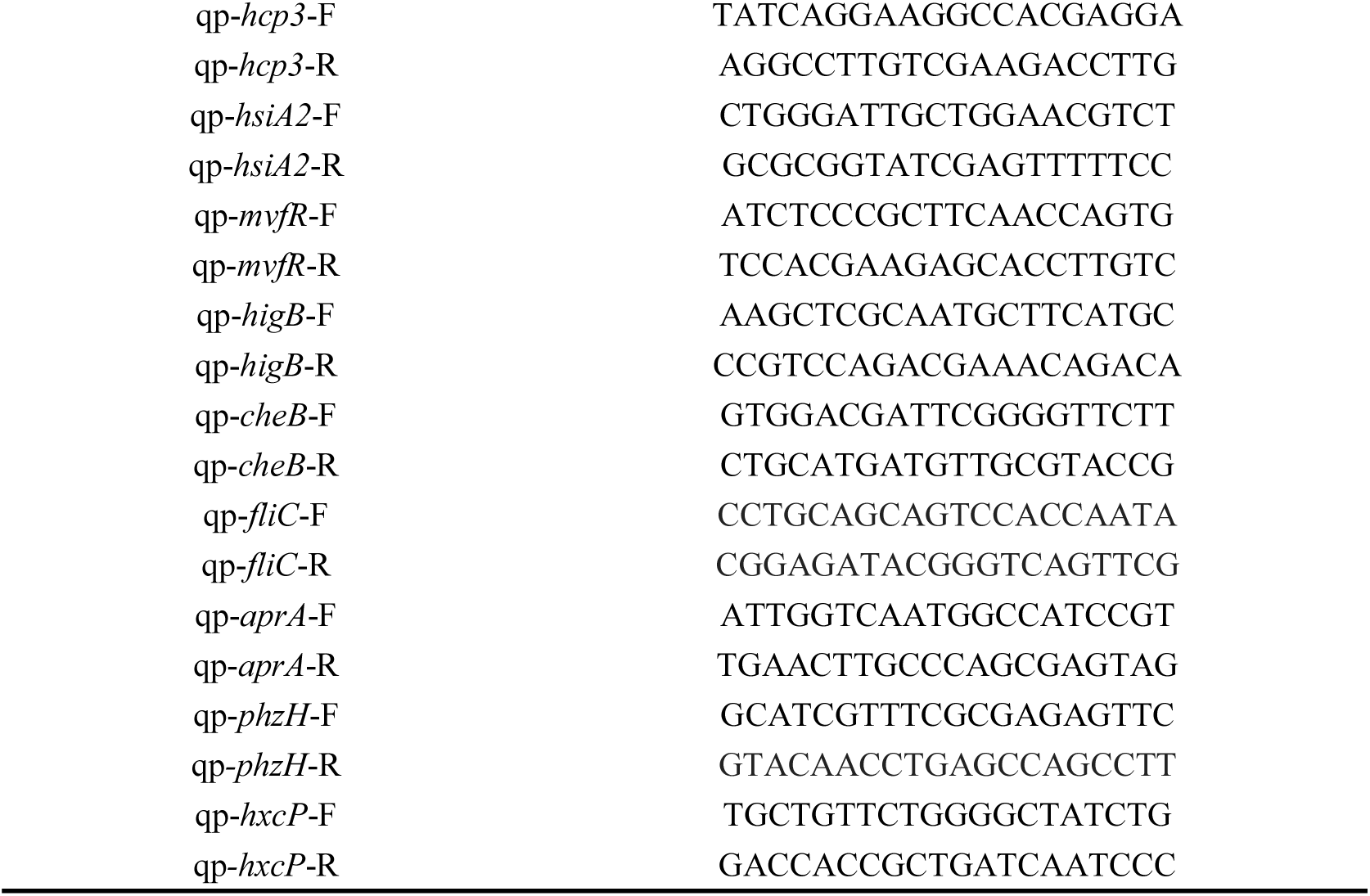
Primers used in this study.

**Table S3.**
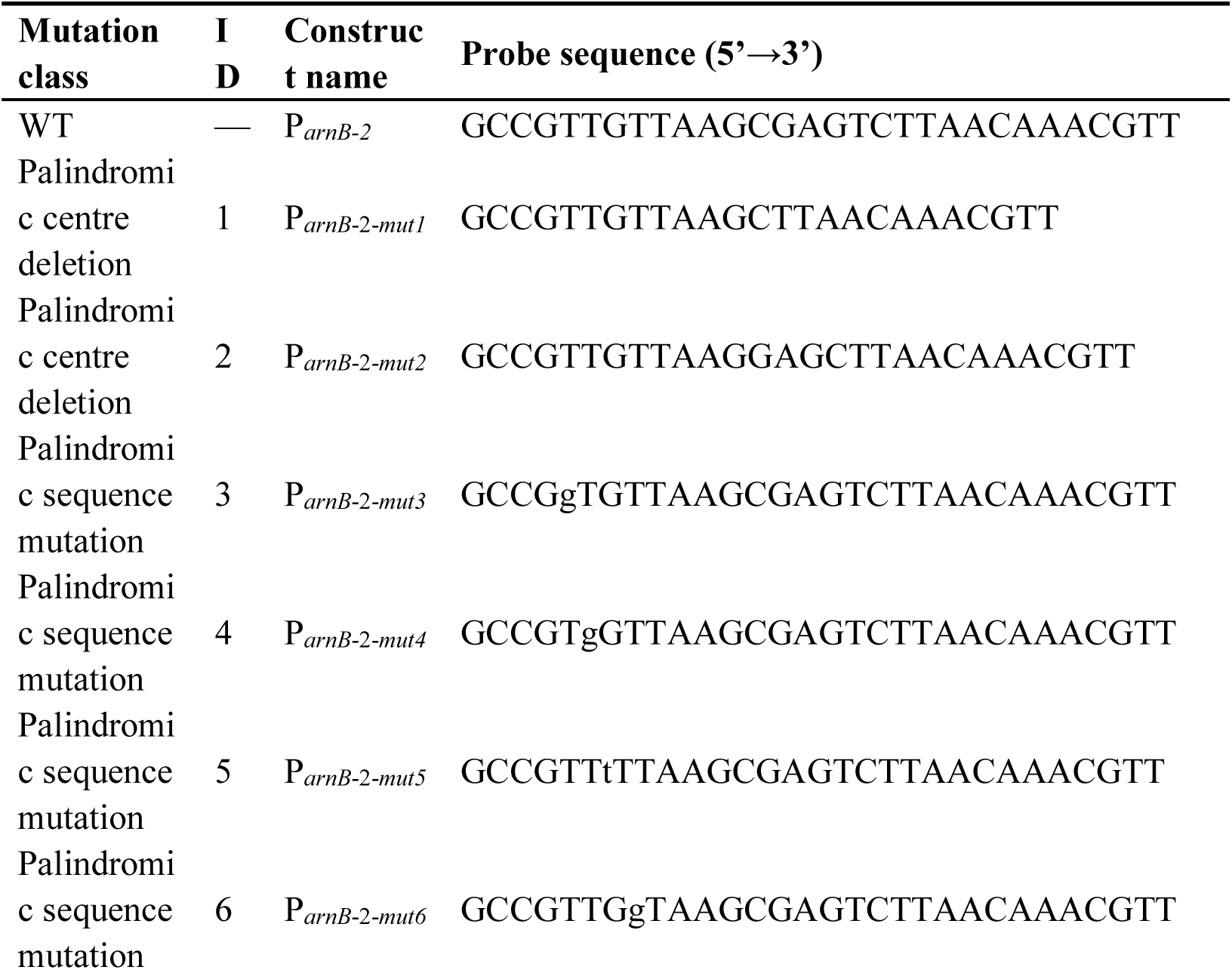
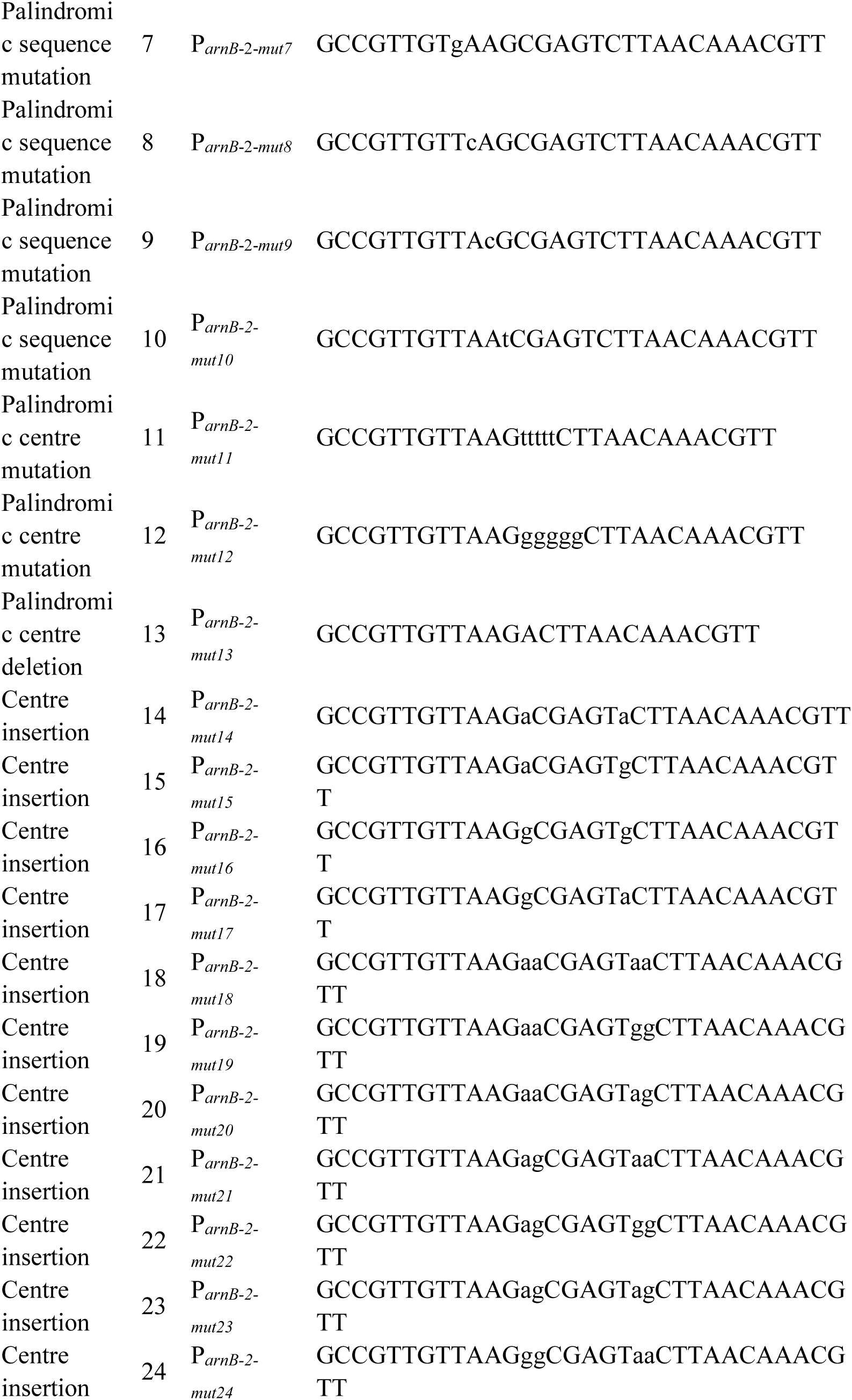
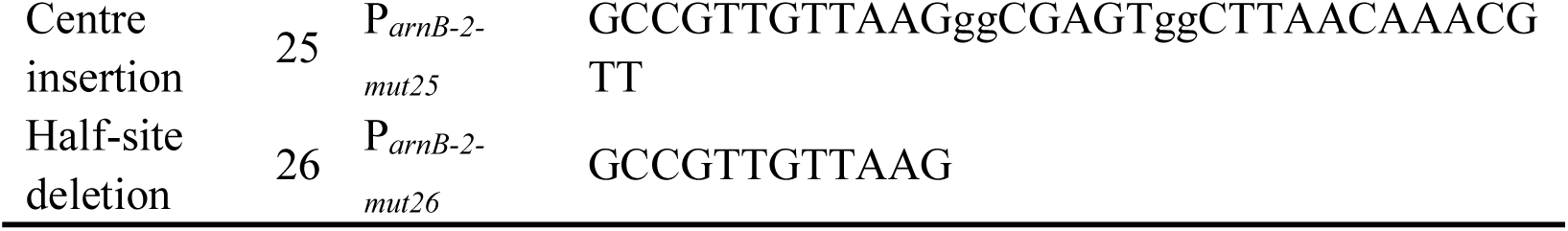
Complete probe set and sequences for the P*_arnB-2_* mutational series used for EMSA in Fig. 5B.

